# Peptide molecular glues select between BET paralogues by exploiting allosteric sites and conformational dynamics

**DOI:** 10.1101/2025.09.24.678353

**Authors:** Karishma Patel, Biswaranjan Mohanty, Alexander Norman, Charlotte Franck, Petr Pachl, Lihua Yang, Xuefei Jing, Xavier J. Reid, James L. Walshe, Paul Solomon, Daniel H. Tran, Daniel Ford, Jason K. K. Low, Lorna Wilkinson-White, Toby Passioura, Richard J. Payne, Hiroaki Suga, Louise J. Walport, Joel P. Mackay

**Affiliations:** School of Life and Environmental Sciences, The University of Sydney, Sydney, NSW 2006, Australia; Sydney Analytical Core Research Facility, The University of Sydney, Sydney, NSW 2006, Australia; School of Chemistry, The University of Sydney, Sydney, NSW 2006, Australia; Australian Research Council Centre of Excellence for Innovations in Peptide and Protein Science, The University of Sydney, Sydney, NSW 2006, Australia; Department of Chemistry, Graduate School of Science, The University of Tokyo, Tokyo 113-0033, Japan; Protein–Protein Interaction Laboratory, The Francis Crick Institute, London NW1 1AT, United Kingdom; Department of Chemistry, Molecular Sciences Research Hub, Imperial College London, London W12 0BZ, United Kingdom

**Keywords:** mRNA-display, RaPID, cyclic peptides, BET family proteins, BRD2, BRD4, bromodomain, drug discovery, protein dynamics

## Abstract

Achieving selective target inhibition is critical for minimising drug side effects. This can be especially challenging when targeting individual members of protein families with high sequence similarity. A well-recognised example is the Bromodomain and Extraterminal domain (BET) family of proteins. Chemical inhibition of the acetyllysine (AcK)-binding bromodomains (BDs) of BET proteins has shown considerable promise in a range of disease models. However, despite over a decade of medicinal chemistry efforts, it has proven challenging to develop BET BD inhibitors that exhibit high selectivity between BET family paralogues. Cyclic peptides are versatile scaffolds for therapeutic development and often exhibit high selectivity and affinity for their targets. To explore their potential as selective BET BD inhibitors, we have used mRNA display to identify cyclic peptide ligands for the *N*-terminal BD of BRD2 and BRD4. The structurally diverse cyclic peptides enriched from the selections boast superior selectivity and affinity to previously developed inhibitors. Most strikingly, we isolated cyclic peptides with ∼1000-fold higher affinity for their target BD over the paralogous BDs, far surpassing selectivities reported to date. Our biochemical and structural data suggest that paralogue-selective cyclic peptides act as molecular glues, exploiting both subtle sequence differences at locations far from the AcK-binding pocket and differences in conformational dynamics between BET BD paralogues to achieve this unprecedented level of specificity. This work provides a blueprint for the development of new classes of selective BET inhibitors and, more generally, underscores the potential of exploiting protein dynamics in the design of selective ligands.

## INTRODUCTION

Selectivity is a critical parameter in the design of pharmaceuticals, agrichemicals, and other protein-targeting molecules. Poor selectivity causes efficacy and safety/toxicity issues that derail many clinical trials and leads to adverse effects on beneficial non-target species in the pesticide space^1,2^. In many cases, achieving selectivity can be a challenge because of the high level of conservation that is observed at orthosteric binding sites, such as the ATP-binding pocket of many kinases and the active site of histone deacetylases (HDACs)^3,4^. Molecules that target alternative – allosteric – sites on a target protein, which are often less well-conserved, are considered a solution to the selectivity problem in some cases and have the potential to provide higher effective potency and safety profiles^5–7^. In general, such sites have been difficult to identify and molecules that bind these sites, which can be shallower than typical small-molecule binding sites, are challenging to design rationally.

One class of therapeutic targets for which selectivity has proven a significant issue is the Bromodomain and Extraterminal domain (BET) family of transcriptional coregulators (**Fig. S1**)^8^. These proteins use tandem bromodomains (BDs) to recognize the acetyllysine (AcK) modification that decorates both histones and other gene regulatory proteins (**Fig. S2**)^9–12^. BET proteins have been linked with a growing list of human diseases, ranging from cancer to inflammation, cardiovascular disease, and neurological disorders^13–22^, and numerous drug discovery campaigns have sought BET BD inhibitors over the last two decades^18,23–29^. Approximately 20 BD inhibitors have advanced to clinical trials, mainly targeting haematological cancers and inflammation. Most of these trials, however, have been terminated due to severe dose limiting toxicity, likely arising, at least in part, from the inability of these molecules to distinguish between BET paralogues^23,30–33^. The design of paralogue specific inhibitors has been impaired by the nearly 100% sequence identity of residues in the AcK-binding pocket, although the first few molecules that display some selectivity have recently emerged^34–37^.

To explore new binding modalities beyond small molecules that might provide access to higher selectivity, we have recently used mRNA display to screen genetically reprogrammed cyclic peptide libraries against BET BDs^38–40^. Screens against several of these domains have yielded highly potent peptides that, in some cases, display ∼10–50-fold selectivity for the specific BD against which they were screened.

Here we describe two peptides – 4.1A and 2.1C – isolated in screens against BRD4-BD1 or BRD2-BD1, respectively, that each display >1,000-fold selectivity for their target BD over other BET paralogues. Both peptides act as molecular glues, creating filamentous assemblies of the BD by simultaneously contacting both the orthosteric AcK-binding site and an allosteric site formed by the αB and αC helices. Despite the similar architecture of the complexes, the two peptides exploit entirely distinct mechanisms to achieve their selectivity. 4.1A draws on a single glutamine residue in the αB helix that is unique to BRD2, and which forms a double-headed hydrogen bond to the peptide backbone that controls selectivity. In contrast, the peptide 2.1C makes no BRD2-specific contacts but unexpectedly appears to achieve its selectivity by sensing differences in conformational dynamics between BRD2 and its paralogues.

Our findings highlight several unexplored paths to BET ligand selectivity, namely the use of non-conserved allosteric binding surfaces, the formation of multivalent interactions driven by molecular glues and, most remarkably, the possibility of exploiting differences in conformational dynamics that are often not apparent from analysis of ground-state structures. Currently, these are all difficult avenues to pursue using rational design approaches, underscoring the value of combinatorial display approaches such as mRNA display.

## RESULTS

### RaPID screens against BRD2-BD1 and BRD4-BD1 yield high-affinity peptides that exhibit significant intra-family selectivity

We performed RaPID selections to identify potent and selective cyclic peptide ligands for BRD2-BD1 and BRD4-BD1 (**Fig. S3A**). As described previously,^38,39^ we used a semi-randomised 10^13^-member library of 10–17-residue peptides, which featured initiation with *N*-chloroacetyl-L-tryptophan (*N*-ClAc-Trp) to drive spontaneous cyclisation with the thiol group of a downstream cysteine (**Fig. S3B**). Genetic code reprogramming allowed us to encode a fixed AcK residue at the centre of the library and the flanking randomised sequences could also potentially encode additional AcKs. After five rounds of selection, the enriched peptide pools from the final three rounds were characterised by next-generation DNA sequencing (**Table S1** for the BRD2-BD1 selection and **Table S2** for the BRD4-BD1 selection). Six diverse peptides from each RaPID screen were chosen and synthesised using solid-phase peptide synthesis (**Table S3**; peptide characterisation data are provided in the methodology section of the **Supporting Information**).

We used surface plasmon resonance (SPR) to assess the ability of these peptides to bind the BDs of BRD2, BRD3, and BRD4 (**Fig. 1** and **S4**, and **Table S3**). All peptides bound to the BD against which they were selected with *K*_D_ values of 0.1–10 nM, with the exception of peptide 2.1D, which bound more weakly to its target BRD2-BD1 (*K*_D_ =130 nM) but had a *K*_D_ of 20 nM for BRD3-BD1 (**Fig. 1B** and **Table S3**). All cyclic peptides, apart from 4.1C, exhibited marked selectivity for the BD1 domains, showing ∼40–10,000-fold reduced binding to the BD2s. 2.1A, 2.1D, 2.1E, 4.1B, 4.1D, 4.1E, and 4.1F also displayed significant intra-BD1 selectivity, showing ∼8–300-fold stronger binding to their cognate BD1 (**Fig. 1A** and **1B**). Peptides 2.1C and 4.1A demonstrated even greater selectivity; they bound only their target BD under the conditions of the SPR assay, indicating paralogue and BD selectivity of at least ∼1,000-fold (**Fig. 1C** and **1D**).

**Figure 1.**
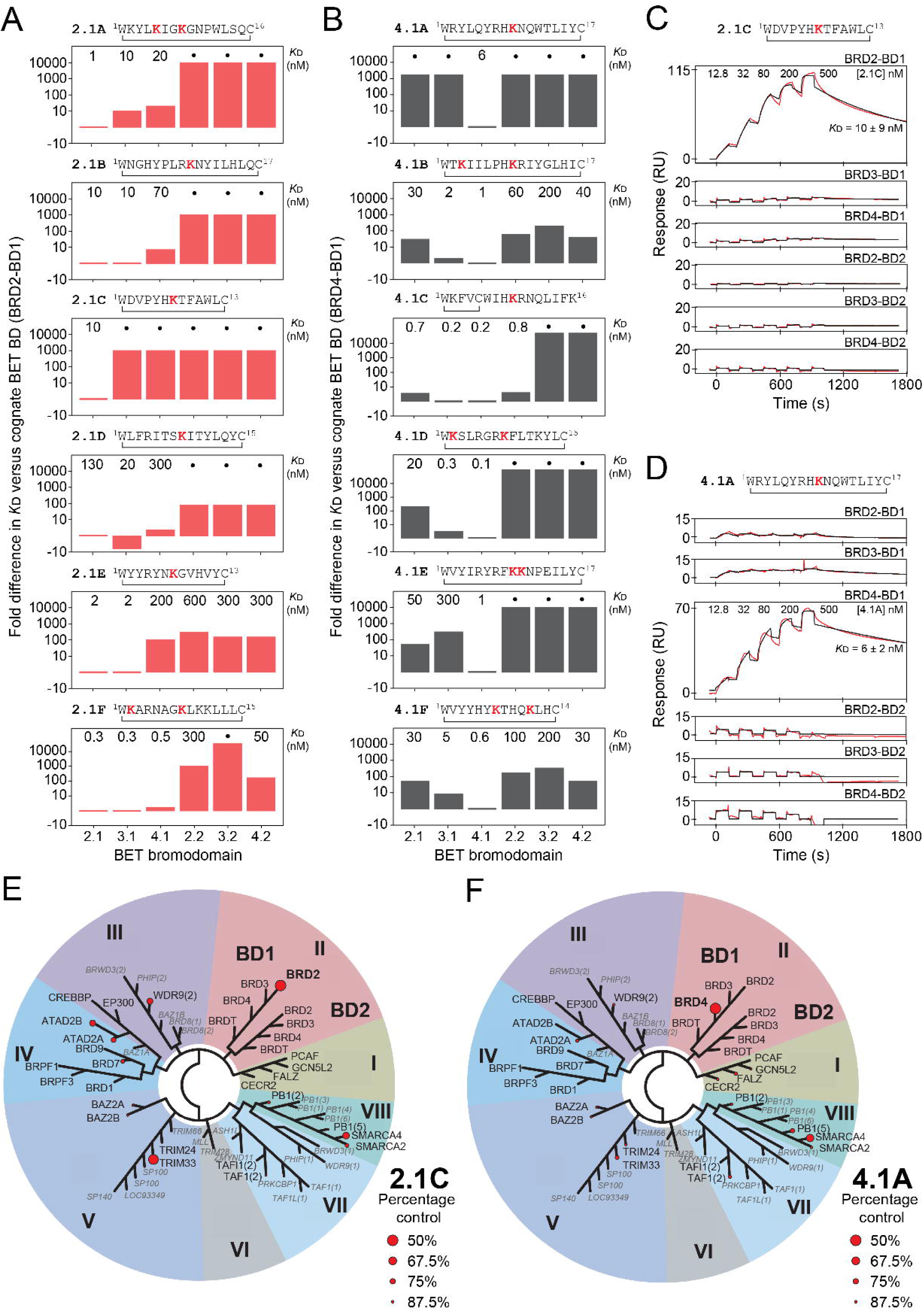
Binding analysis of CPs discovered from RaPID selections against BRD2-BD1 and BRD4-BD1. Bar graphs showing fold change in *K*_D_ for the interactions between **A.** the BRD2-BD1 RaPID CPs and all other BDs from BRD2, BRD3, and BRD4, relative to the binding affinity of the CPs against BRD2-BD1 and **B.** the BRD4-BD1 RaPID CPs and all other BDs from BRD2, BRD3, and BRD4, relative to the binding affinity of the CPs against BRD4-BD1. Fold changes were calculated by normalising the *K*_D_ of the CPs against their affinity for BRD2-BD1/BRD4-BD2 (set to 1). The individual *K*_D_ values for each interaction are provided above the graph and given as the geometric mean of a minimum of three independent measurements. Uncertainties are shown in **Table S3**. Black circles are used to indicate interactions that were too weak to be detected (and are therefore weaker than >10 μM). The names and sequences of the CPs are given above each graph (AcK residues are shown in *red* and the residues that are cyclised are indicated). **C.** Representative SPR sensorgrams (*red*) for binding of 2.1C to all the BDs of BRD2, BRD3, and BRD4. Fits to a 1:1 binding model (*black*) are shown for each sensorgram. BDs other than BRD2-BD1 don’t bind measurably to 2.1C. **D.** Representative SPR sensorgrams (*red*) for the binding of 4.1A to all BDs of BRD2, BRD3, and BRD4. Fits to a 1:1 binding model (*black*) are shown for each sensorgram. BDs other than BRD4-BD1 don’t bind measurably to 4.1A. **E,F**. *TREE*spot phylogenetic tree showing the binding of **E.** 1 μM 2.1C and **F.** 1 μM 4.1A to 32 diverse human BDs in a BROMO*scan* competition binding assay (Eurofins DiscoverX). The larger spot size and lower percentage control correlate with stronger affinity. BDs that were tested are labelled in black and untested BDs are in grey. The BD1 and BD2 branches of the tree are labelled.

To further assess the striking paralogue specificity of 2.1C and 4.1A, we conducted a BROMO*scan* assay (performed by Eurofins DiscoverX) to profile the relative affinity of these peptides for 32 diverse human BDs. 2.1C displayed marked selectivity for BRD2-BD1 (**Fig. 1E**). No binding was observed for any of the other seven BET family BDs, consistent with our SPR data, although comparable binding was observed for the unrelated BD from TRIM33 (26% sequence identity with BRD2-BD1). Likewise, 4.1A demonstrated a clear preference for BRD4-BD1 among the eight BET-family BD, although some binding to the unrelated BD from SMARCA4 (24% sequence identity with BRD4-BD1) was observed (**Fig. 1F**).

### 4.1A acts as a head-to-tail molecular glue and selects BRD4-BD1 by recognising a non-canonical BD surface

We determined the X-ray crystal structure of the BRD4-BD1:4.1A complex to define the mechanism underlying the paralogue selectivity of the peptide (2.9-Å resolution, PDB ID: 9MPI, **Fig. 2A**, **Table S4**). 4.1A forms a highly ordered β-hairpin that engages two molecules of BRD4-BD1, effectively acting as a molecular glue (**Fig. 2A** *left* and **Fig. S5A**). One BD (BRD4-BD1-A) is recruited via an AcK residue (K9ac) that forms the canonical hydrogen bond with N140 in the AcK-binding pocket of the BD (**Fig. 2A** *bottom right inset*). However, the second BRD4-BD1 (BRD4-BD1-B) binds 4.1A via a completely different surface, which is formed by the αB-αC helices (**Fig. 2A** *left* and *top right inset*). Because of this architecture, both BRD4-BD1 and 4.1A can simultaneously bind two molecules of peptide and BD, respectively (**Fig. 2A** and **Fig. S5B**). 4.1A therefore has the potential to drive the formation of a filamentous structure and, indeed, we observe the formation of a helical filament in the crystal of the complex (**Fig. 2B** and **S5C**). Consistent with this observation, the ^15^N-heteronuclear single-quantum coherence (HSQC) nuclear magnetic resonance (NMR) NMR spectrum of BRD4-BD1 displays widespread broadening following the addition of 4.1A, suggesting the formation of a higher molecular weight species in solution (**Fig. 2C** *top*), and extended incubation leads to precipitation. In contrast, no change to the ^15^N-HSQC spectrum of BRD4-BD2 is observed upon the addition of 4.1A and no precipitation is observed, consistent with our SPR data and suggesting that 4.1A does not induce non-specific aggregation of BET bromodomains (**Fig. 2C** *bottom*).

**Figure 2.**
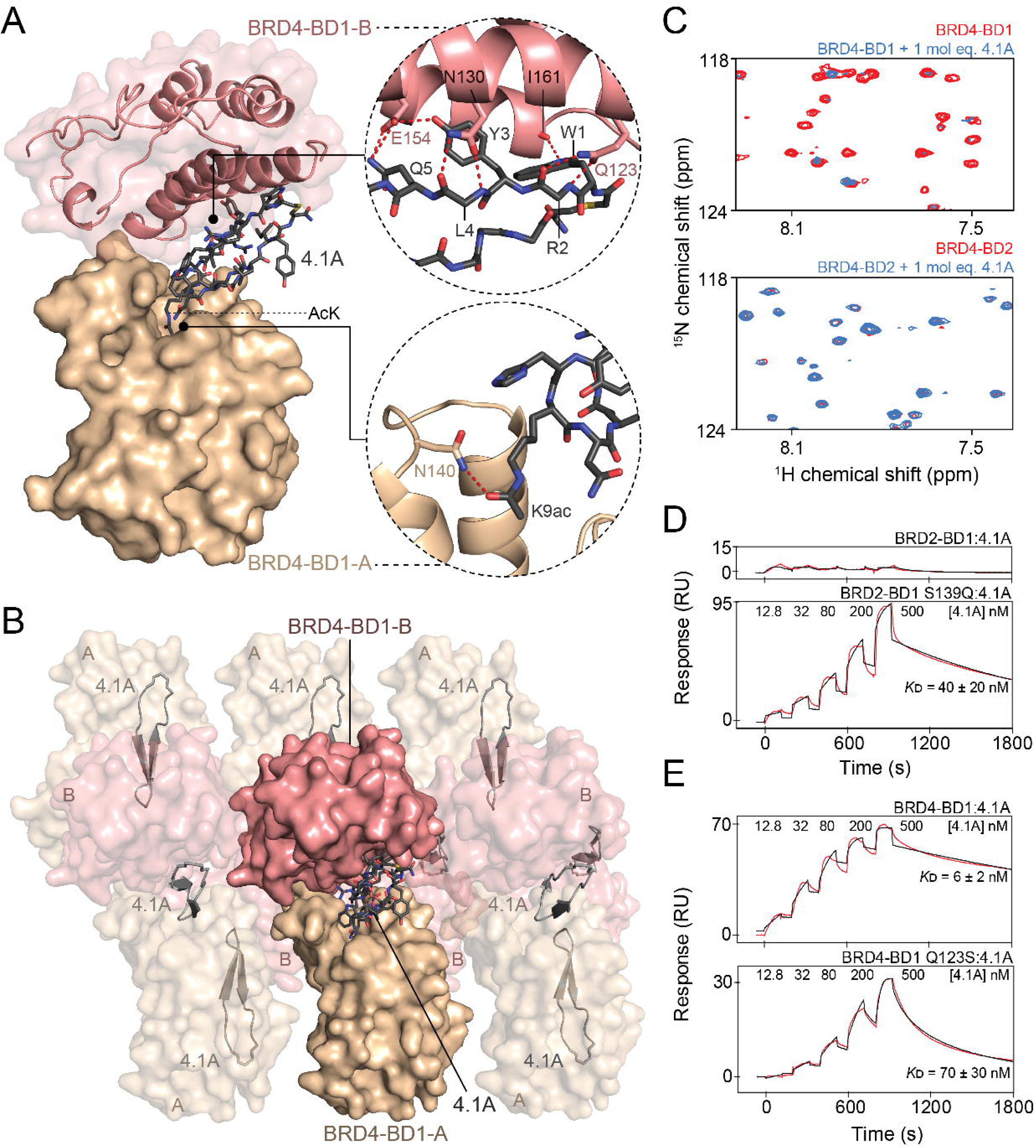
Characterisation of the interaction between BRD4-BD1 and 4.1A. **A.** A side view of the X-ray crystal structure of the complex formed between 4.1A and two molecules of BRD4-BD1 (2.9-Å resolution, PDB ID: 9MPI). The peptide is represented as sticks in *dark grey* and the two BDs bound to 4.1A are shown in *wheat* (BRD4-BD1-A) and *salmon* (BRD4-BD1-B) as surfaces. *Top right inset*: Close up of the AcK-independent interactions between 4.1A and BRD4-BD1-B. Hydrogen bonds formed between BRD4-BD1-B and 4.1A residues are indicated by *red dashed lines*. *Bottom right inset*: Close-up of the hydrogen bond formed between the sidechain carbonyl of K9ac of 4.1A and N140 of BRD4-BD1-A (*red dashed line*). **B**. A short section of the filamentous structure formed by the BRD4-BD1:4.1A complex. The BDs (BRD4-BD1-A labelled as A and BRD4-BD1-B labelled as B) and peptide (4.1A) are indicated. **C.** Sections of ^15^N-HSQC spectra of BRD4-BD1 (*top*) and BRD4-BD2 (*bottom*) alone (*red*) or in the presence of one molar equivalent of 4.1A (*blue*). Widespread signal loss is observed only for BRD4-BD1. **D.** Representative SPR sensorgrams (*red*) for the binding of 4.1A to the BRD2-BD1-S139Q mutant. SPR data for the native BRD2-BD1:4.1A interaction (from **Fig. 1D**) are also shown for comparison. Fits to a 1:1 binding model (*black*) are shown. The *K*_D_ for the interaction between 4.1A and BRD2-BD1-S139Q is given as the geometric mean (± standard error) of a minimum of three independent measurements. **E.** Representative SPR sensorgrams (*red*) for the binding of 4.1A to the BRD4-BD1-Q123S mutant. The SPR data for the native BRD4-BD1:4.1A interaction are also shown for comparison. Fits to a 1:1 binding model (*black*) are shown. The *K*_D_ for the interaction between 4.1A and BRD4-BD1-Q123S is provided as the geometric mean (± standard error) of a minimum of three independent measurements.

To corroborate our crystal structure, we assessed the binding of 4.1A variants in which BD-contacting residues were mutated to alanine. Substitution of 4.1A W1 and Y3, which contact the αB-αC surface of BRD4-BD1-B, reduced binding by ∼17-and ∼650-fold, respectively (**Fig. S5D**). However, substitution of 4.1A Q5, which also interacts with the αB-αC helices, did not significantly impact binding (**Fig. S5D**). Surprisingly, a K9ac to alanine mutant (4.1A K9acA) displayed no reduction in binding (*K*_D_ = 2 nM) and retained selectivity for BRD4-BD1, indicating that the AcK-pocket interaction does not make a significant contribution to affinity (**Fig. S5D** and **S6A**). Titration of the 4.1A K9acA peptide into ^15^N-labelled BRD4-BD1 led to selective reductions in ^15^N-HSQC signal intensity and the appearance of new signals (**Fig. S6B**). The most significant reductions in signal intensity induced by the addition of 0.5 molar equivalents of 4.1A K9acA map to the αB-αC helices, rather than the AcK binding site, suggesting the formation of a 1:1 complex with the BRD4-BD1-B subunit from the crystal structure (**Fig. S6C**).

We next sought to determine the basis of the selectivity of 4.1A for BRD4-BD1. All residues in proximity to the 4.1A contact surface on BRD4-BD1-A are conserved in the BD1s of BRD2 and BRD3, suggesting that this part of the interaction does not bestow paralogue specificity. On the BRD4-BD1-B contact surface (the αB-αC helices), only a single residue – Q123 – is unique to BRD4-BD1; this residue is a serine in the other BET BD1s (**Fig. 2A** *top right inset*). We therefore introduced a glutamine at this location in BRD2-BD1 (BRD2-BD1 S139Q). This single-residue substitution was sufficient to install robust binding of 4.1A (*K*_D_ = 40 nM; **Fig. 2D**). Conversely, replacement of Q123 with a serine in BRD4-BD1 decreased binding by ∼10-fold (**Fig. 2E**). Whilst not the sole determinant of specificity, these data suggest that 4.1A exploits a single-residue difference on a non-canonical surface of the BDs to derive exceptional paralogue specificity.

### 2.1C also exploits bivalent binding to selectively engage its target BD

To explore whether 2.1C achieves selectivity in a similar manner to 4.1A, we next determined the X-ray crystal structure of 2.1C in complex with BRD2-BD1 (2.9 Å resolution, PDB ID: 9MPJ, **Fig. 3A**, **Table S4**). The overall architecture of the complex closely resembles that of the 4.1A-BRD4-BD1 complex, even though the structure of 2.1C is completely different to 4.1A. 2.1C forms a compact irregular structure with five internal hydrogen bonds (**Fig. S7A**) and also engages two BDs in a way that, in this case, gives rise to a considerable interaction interface between the two BDs (**Fig. 3A** *centre* and *top left inset* and **S7B**). An AcK (K7ac) in 2.1C binds the canonical AcK-binding pocket of one BRD2-BD1 (BRD2-BD1-A; **Fig. 3A** *bottom right inset*), and the peptide simultaneously contacts the αB-αC surface of a second BRD2-BD1, using residues Y5, W11 and L12 (**Fig. 3A** *top right inset*). As with the BRD4-BD1:4.1A complex, this architecture means that BRD2-BD1 can bind two molecules of 2.1C at the same time and, accordingly, a continuous helical filament is observed in the crystal structure (**Fig. 3B** and **S7C**). Also mirroring our observations for 4.1A, titration of one molar equivalent of 2.1C into ^15^N-labelled BRD2-BD1 caused a widespread disappearance of signals in a ^15^N-HSQC spectrum (**Fig. 3C** *top*), as well as the formation of visible precipitate. In contrast, a ^15^N-HSQC of BRD2-BD2 was unchanged following addition of 2.1C (**Fig. 3C** *bottom*). The formation of these filaments is consistent with the formation of soluble aggregates and precipitates in the ^15^N-HSQC NMR titration.

**Figure 3.**
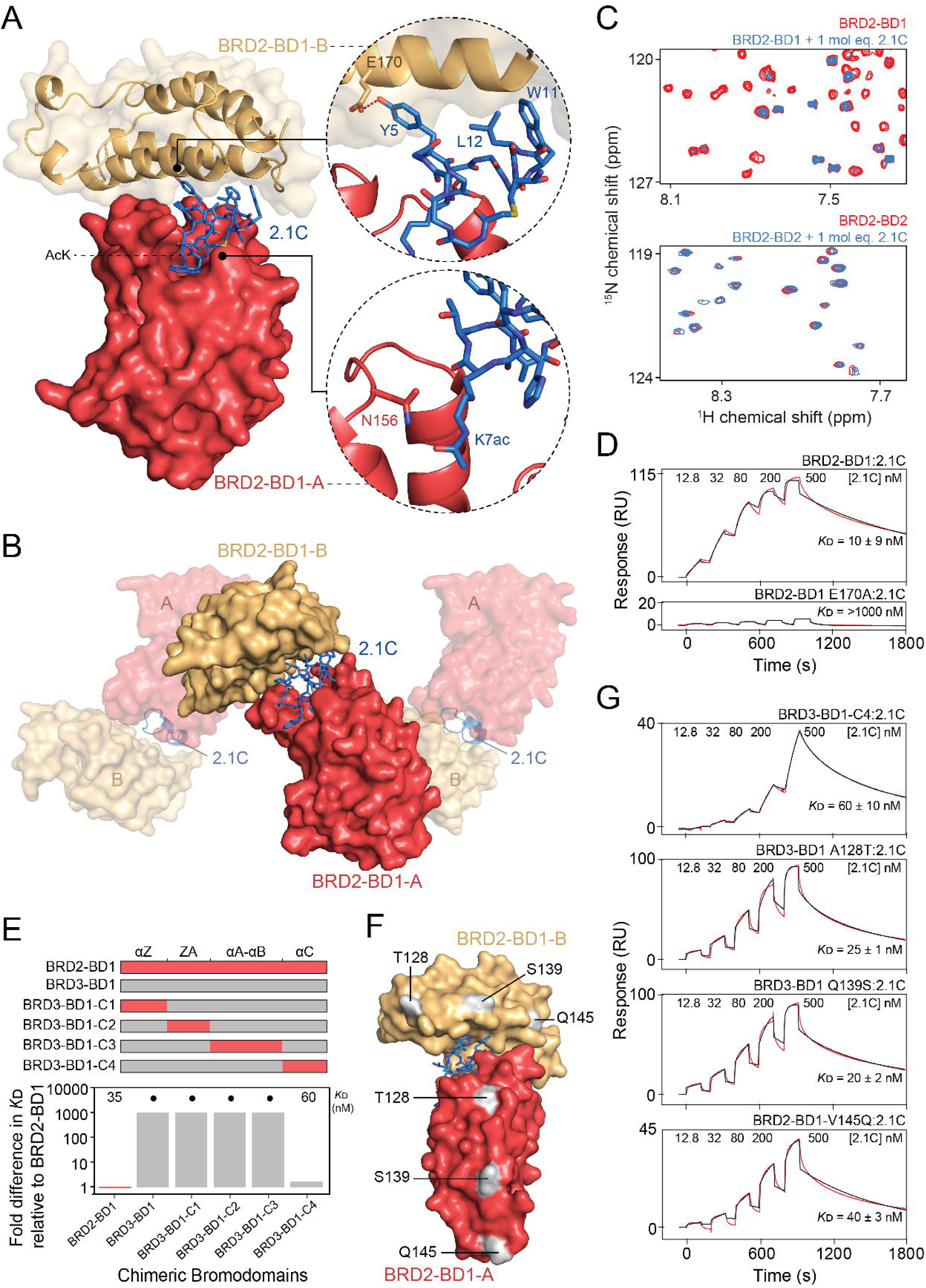
Characterisation of the interaction between BRD2-BD1 and 2.1C. **A.** A side view of the X-ray crystal structure of the complex formed between 2.1C and two molecules of BRD2-BD1 (2.9 Å resolution, PDB ID: 9MPJ). The peptide is represented as sticks in *blue* and the two BDs bound to 2.1C are shown in *coral* (BRD2-BD1-A) and *light orange* (BRD2-BD1-B) as surface representation. *Top right inset*: Close up of the AcK-independent interactions between 2.1C and BRD2-BD1-B. The hydrogen bond formed between E170 of the BD and Y5 of 2.1C is indicated by the *red dashed line*. *Bottom right inset*: The hydrogen bond formed between the sidechain carbonyl of K7ac of 2.1C and the sidechain amide of N156 of BRD2-BD1-A is represented by the *red dashed line*. **B.** A section of the filamentous structure formed by the BRD2-BD1:2.1C complex. The BDs (BRD2-BD1-A labelled as A and BRD2-BD1-B labelled as B) and peptide are indicated. **C.** Sections of ^15^N-HSQC spectra of BRD2-BD1 (*top*) and BRD2-BD2 (*bottom*) alone (*red*) or in the presence of one molar equivalent of 2.1C (*blue*). **D.** Representative SPR sensorgram (*red*) for the binding of 2.1C to the BRD2-BD1-E170A mutant. SPR data for the native BRD2-BD1:2.1C interaction (from **Fig. 1C**) are shown for comparison. **E.** *Top*: Schematic showing the BRD2-BD1 and BRD3-BD1 chimeric BDs. BRD2-BD1 is coloured in *coral* and BRD3-BD1 is coloured in *grey*. *Bottom*: Fold change in *K*_D_ (measured by SPR) for the interactions between the chimeric BDs and 2.1C relative to the binding of the peptide to BRD2-BD1. *K*_D_ values are given as the geometric mean of a minimum of three independent measurements. Black circles are used to indicate interactions that could not be detected. **F.** The position and identity of the three residues from BRD2-BD1 present in BRD3-BD1-C4 mapped onto the structure of the BRD2-BD1:2.1C complex. **G.** Representative SPR sensorgrams (*red*) for the binding of 2.1C to BRD3-BD1-C4, BRD3-BD1 A128T, BRD3-BD1 Q139S, and BRD3-BD1 V145Q. Fits to a 1:1 binding model (*black*) are shown. *K*_D_ values are provided as the geometric mean (± standard error) of at least three independent measurements.

To establish the energetic drivers for complex formation, we first examined the BD-contacting residues in 2.1C. Mutation of K7ac to alanine abolished binding of the peptide altogether and mutation of any of Y5, W11 or L12 to alanine reduced the affinity for BRD2-BD1 by 60–100-fold, establishing the importance of both peptide-protein interfaces (**Fig. S7D**). Consistent with these data, an E170A mutation in BRD2-BD1, disrupting the hydrogen bond with 2.1C Y5, also reduced affinity by ∼100-fold, reinforcing the importance of the αB-αC surface for the interaction (**Fig. 3D**). We also noted that, although 2.1C and 4.1A have distinct structures and specificity, both peptides use a tyrosine to make a hydrogen bond to this conserved glutamate residue (**Fig. 2A** *top right inset*, **3A** *top right inset*, and **S7E**).

Structural analyses found that the 2.1C W11A mutation disrupts the architecture of the BRD2-BD1:2.1C complex. A structure of 2.1C W11A bound to BRD2-BD1 (**Fig. S8A**, 3.1 Å resolution, PDB ID: 9MPL, **Table S5**) adopts the same conformation as the wildtype 2.1C peptide (**Fig. S8B**) and mirrors the AcK-dependent interaction. However, the peptide engages a second copy of BRD2-BD1 in a very different manner that leads to the lower observed affinity (**Fig. S8A-D**). Interestingly, 2.1C W11A gains affinity for BRD3-BD1, such that there is less than a two-fold difference in affinity for BRD2-BD1 and BRD3-BD1 (600 nM vs 1 μM, respectively; **Fig. S8C**). In line with this observation, a crystal structure of BRD3-BD1 bound to 2.1C W11A (**Figure S8A**, 2.6 Å resolution, PDB ID: 9MPM, **Table S5**) is essentially identical to the structure featuring BRD2-BD1. Surprisingly, the conserved glutamate described above again plays a key role in the interaction – this time forming two hydrogen bonds with the peptide backbone (**Fig. S8D** *inset*).

Together, these data show that 2.1C, like 4.1A, forms a bivalent complex by both inserting an AcK into the canonical AcK-binding pocket and contacting the αB-αC surface of BRD2-BD1.

### Chimeric BDs reveal individual residues that allosterically drive affinity and specificity

We next asked which BRD2-BD1 residues drive the high paralogue selectivity of 2.1C. Unexpectedly, we found that there are no amino acids within 5 Å of 2.1C at either binding interface that are unique to BRD2. We did note a single residue unique to BRD2-BD1 (G109; asparagine in BRD3 and BRD4) that lies at the BD:BD interface of the complex on the ZA loop (**Fig. S9A** and **S9B**). A BRD2-BD1-G109N mutant, however, bound 2.1C with the same affinity as wildtype BRD2-BD1, albeit with a slightly faster off-rate (**Fig. S9C**). Similarly, mutating the only two other ZA-loop residues unique to BRD2-BD1 to the corresponding residue in BRD3-BD1 (R100Y and V106I) did not significantly impact affinity (**Fig. 9D** and **S9E**).

Given that no 2.1C-contacting residues endowed BRD2-BD1 with selectivity for 2.1C, we probed paralogous sequence differences across the entire domain. We created a set of chimeric BDs in which we introduced sets of residues from BRD2-BD1 into BRD3-BD1 (**Fig. 3E** *top panel*). Introduction of BRD2-BD1 residues from the αZ helix, the ZA loop, αA helix, or αB helix did not allow BRD3-BD1 to bind 2.1C (**Fig. 3E**, **S9A**, and **S10**). However, a chimera bearing the αC helix from BRD2 (BRD2-BD1-C4; three changes **Fig. 3F**, **S9A**, and **S10**), bound 2.1C with high affinity (*K*_D_ = 60 nM; within 2-fold of the affinity for BRD2-BD1 measured on the day; **Fig. 3E** and **3G**).

The sidechains of all three of the substituted residues in the αC helix are surface exposed, and none contact 2.1C (**Fig. 3F**). To pinpoint the critical residue(s) that bestow the ability to bind 2.1C, we made the individual single-site mutations to BRD3-BD1 (A128T, Q139S and V145Q). Any one of the three substitutions allowed BRD3-BD1 to bind 2.1C with similar affinity to BRD2-BD1 (within ∼2-fold; **Fig. 3G**), which represents increases of ∼200–500-fold in affinity relative to BRD3-BD1. ^15^N-HSQC spectra and chemical shift assignment for BRD3-BD1 A128T, BRD3-BD1 Q139S, and BRD3-BD1 V145Q show that the overall structure is conserved in these mutants, with chemical shift changes largely localized to the sites of mutation (**Fig S11A**-**B**, **Fig. S12A**, and **Tables S6-9**) and an X-ray crystal structure of BRD3-BD1-A128T (**Fig. S12B**, 1.6 Å resolution, PDB ID: 9MPN, **Table S5**) is essentially identical to BRD3-BD1 (RMSD = 0.25 Å over structured backbone residues).

These data show that any of three specific point mutations in BRD3-BD1 – none of which introduce significant structural changes nor contact 2.1C – can essentially abolish the dramatic paralogue-level specificity that peptide 2.1C displays for BRD2-BD1, by allowing high affinity binding of the peptide to BRD3-BD1.

### Specificity-determining residues modulate milli-to microsecond timescale dynamics

Given that there was no clear structural basis for the specificity of 2.1C, we hypothesized that differences in conformational dynamics might underlie our observations. We therefore probed μs– ms backbone dynamics in BRD2-BD1, BRD3-BD1 and BRD4-BD1 by measuring H^N^-detected Carr-Purcell-Meiboom-Gill (CPMG) relaxation dispersion data for each domain. Relaxation dispersion data have been used extensively to detect structural fluctuations that involve a minor state with a population too small to be directly observable in NMR spectra^41,42^. **Fig. 4A** and **B** show data and plots of *R_ex_*versus residue number for each domain; *R_ex_* measures the magnitude of residue-level conformational dynamics and is typically sensitive to the presence of minor conformations down to ∼1% or less. Remarkably, despite pairwise sequence identities of ∼80–90%, the three domains display distinct conformational landscapes. The dynamics in BRD3-BD1 are dominated by motion of the ∼25-residue ZA loop that forms a major part of the AcK-binding pocket (**Fig. 4C**). BRD4-BD1 also exhibits conformational flexibility in the ZA loop, but additionally shows significant dynamics at the opposite end of the structure where the *N*-terminal part of the αZ helix contacts the *C*-terminal portions of αA and αC (**Fig. 4C**). In stark contrast, the data for BRD2-BD1 show no indication of μs–ms timescale dynamics at any position (**Fig. 4C**).

**Figure 4.**
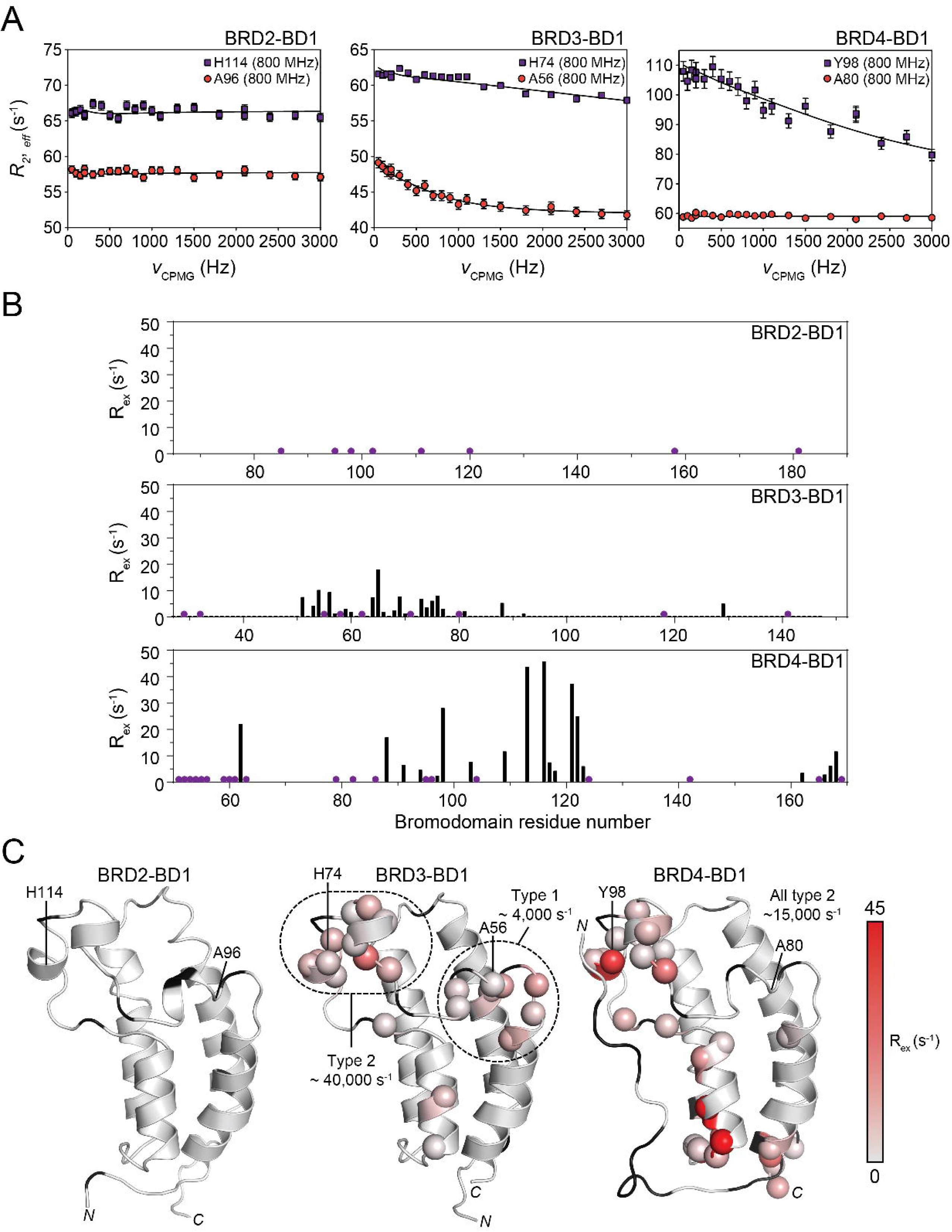
BET-BD1 domains display distinct. μ**s–ms dynamics. A.** Exemplar H^N^ relaxation dispersion profiles for two corresponding residues (shown in blue and red) from BRD2-BD1, BRD3-BD1 and BRD4-BD1. Neither residue shows measurable dynamics in BRD2; in BRD3, H74 exhibits fast-timescale dynamics while A56 undergoes much slower motion; in BRD4, Y98 undergoes fast motion and A80 shows no measurable dynamics. **B.** Plots of the magnitude of *R_ex_*(the contribution of μs–ms dynamics to the *transverse* relaxation rate) as a function of residue number for each BD1 domain. *Purple* circles indicate residues that could not be measured (either proline or unassigned residues). No dynamics were observed for BRD2-BD1. **C.** Mapping of μs–ms dynamics onto each BD. Cα atoms of residues with measurable H^N^ relaxation dispersion are shown as spheres and are coloured on the indicated *grey* to *red* scale according to the magnitude of *R_ex_* displayed by that residue. The timescale of the motion, obtained from fitting to a two-site exchange model, is indicated. Residues in *black* could not be measured.

Fitting of the relaxation dispersion data also provides estimates of both the timescale of the local motion at each residue and the populations of the interconverting states. For BRD3-BD1, conformational exchange at the *N*-terminal end of the ZA loop (which we label type 1 dynamics, and also includes the nearby L129 in the αC helix) takes place with an overall rate constant of ∼4,000 s^-1^, whereas a separate (type 2) motional regime exists at the central portion of the ZA loop with a rate constant of ∼40,000 s^-1^ (**Fig. 4C**, **S13**, and **S14**). For BRD4-BD1, all dynamics takes place on the faster type 2 timescale with a rate constant of ∼15,000 s^-1^ (**Fig. 4C** and **S15**).

To determine whether a causal link exists between BD conformational dynamics and 2.1C-binding capacity, we measured H^N^ relaxation dispersion data for each of the BRD3-BD1 point mutants that bound strongly to 2.1C (**Fig. S16**). **Fig. 5A** and **B** shows that the A128T mutation almost eliminates conformational dynamics at the αA site and significantly dampens the dynamics in the αZ helix and ZA loop. Similarly, the Q139S and V145Q mutations broadly dampen BD dynamics, though to a smaller extent. To further test the idea that changes in BD dynamics are causally linked to 2.1C affinity, we sought a BRD2-BD1 mutant with reduced affinity for 2.1C. Mutation of three residues in the ZA loop (R100Y, V106I, and G109N) reduces the *K*_D_ by ∼1000-fold to >1 μM (**Fig. 6A**). In accord with our hypothesis, H^N^ relaxation dispersion data reveal the appearance of conformational dynamics with a spatial distribution that overall resembles both BRD3-BD1 and BRD4-BD1 (**Fig. 6B** and **C**) and that occurs on the type 2 timescale (*k_ex_* ∼35,000 s^-1^) (**Fig. S17**). Changes in the dynamics of the ZA loop triple mutant are accompanied by differences in the HSQC relative to wildtype BRD2-BD1 which largely map to the ZA loop region (**Fig. S18**).

**Figure 5.**
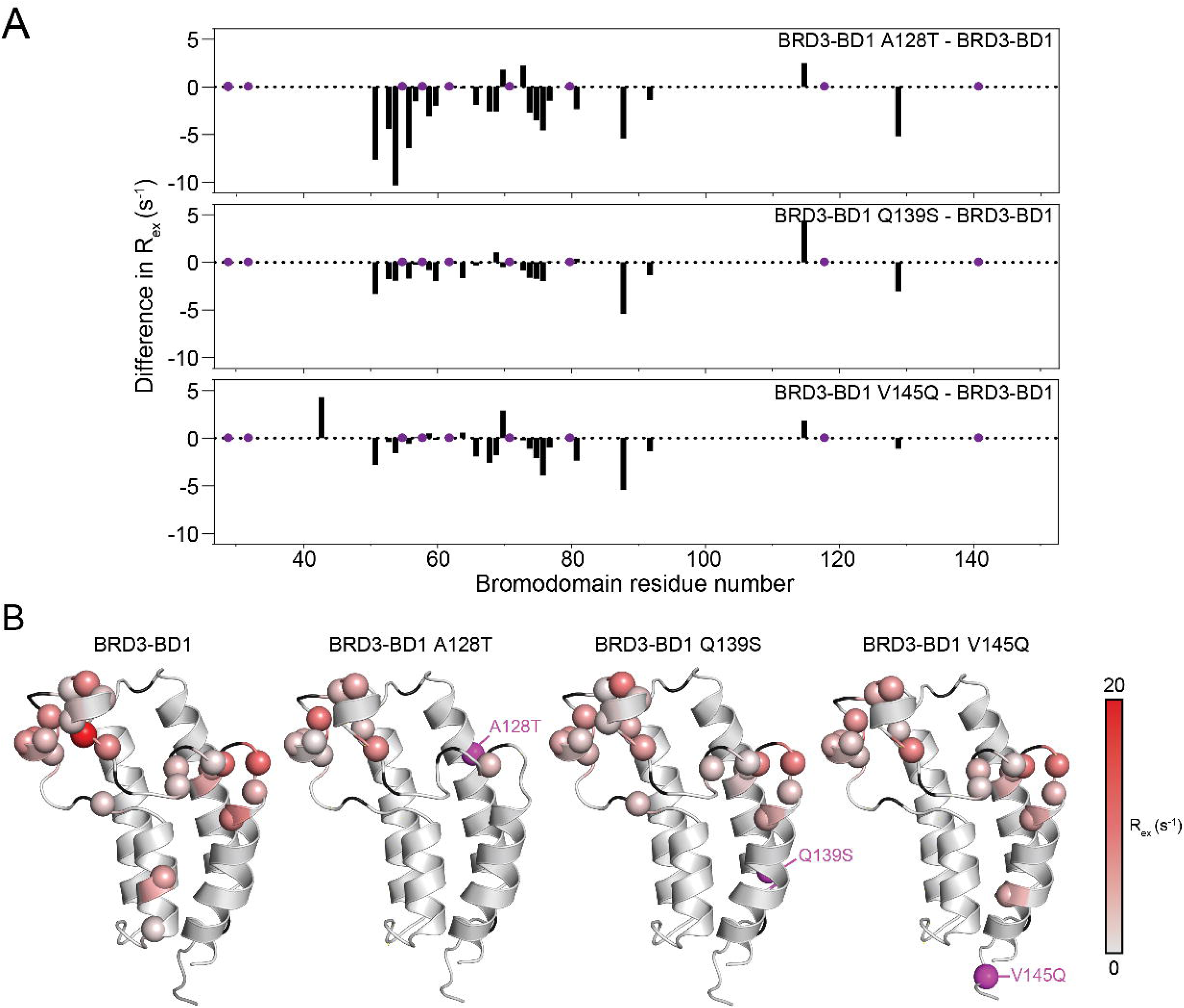
Single-residue mutants of BRD3-BD1 that bind 2.1C have significantly dampened. μ**s–ms dynamics. A.** Difference plots showing the change in magnitude of *R_ex_* (the contribution of μs–ms dynamics to the *transverse* relaxation rate) as a function of residue number for the A128T, Q139A and V145Q point mutants of BRD3-BD1. Plots show *R_ex_*(mutant) – *R_ex_*(BRD3-BD1). **B.** Mapping of μs–ms dynamics onto each BD. Cα atoms of residues with measurable H^N^ relaxation dispersion are shown as spheres and are coloured on the indicated *grey* to *red* scale according to the magnitude of *R_ex_* displayed by that residue. Residues in *black* could not be measured. The position of each mutation is shown as a *purple* sphere. Wildtype BRD3-BD1 data are taken from **Fig. 4C**.

**Figure 6.**
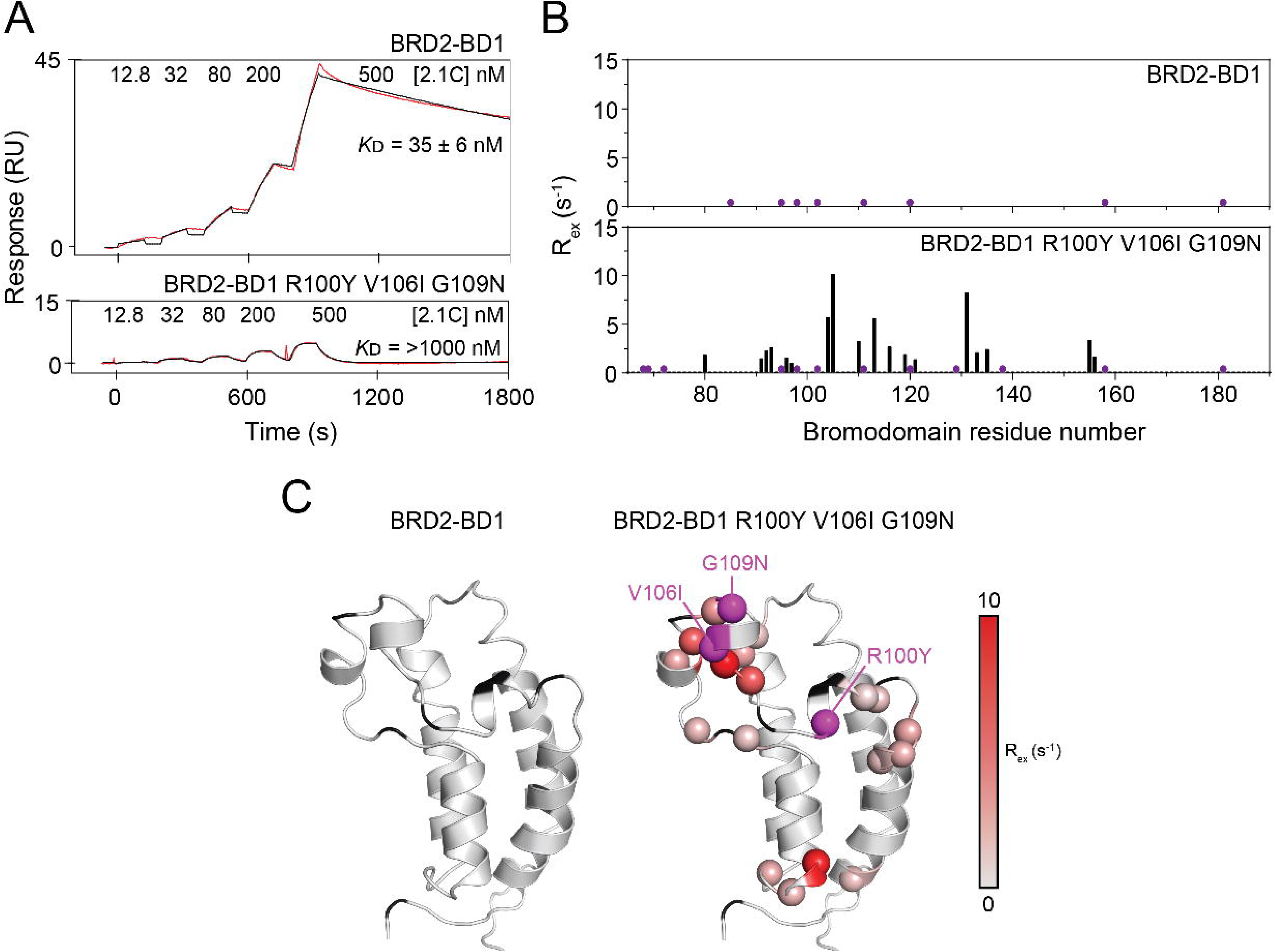
A BRD2-BD1 mutant with reduced affinity for 2.1C displays significant. μ**s–ms dynamics. A.** Representative SPR sensorgrams (*red*) for the binding of 2.1C to BRD2-BD1 (*top*) and to the indicated triple point mutant (*bottom*). Fits to a 1:1 binding model (*black*) are shown. The *K*_D_ for each interaction is given as the geometric mean (± standard error) of a minimum of three independent measurements. **B.** Plots of the magnitude of *R_ex_* as a function of residue number for BRD2-BD1 (from **Fig. 4B**) and the triple mutant. Purple circles indicate residues that could not be measured (either proline or unassigned residues). Substantial dynamics was observed in the mutant. **C.** Mapping of μs–ms dynamics onto BRD2-BD1 and the triple mutant. Cα atoms of residues with measurable H^N^ relaxation dispersion are shown as spheres and are coloured on the indicated *grey* to *red* scale according to the magnitude of *R_ex_* displayed by that residue. Residues in *black* could not be measured. The pattern of dynamics was very similar to that observed for wildtype BRD3-BD1 and BRD4-BD1.

Overall, these data are consistent with the idea that 2.1C achieves a high level of specificity between the BD1 domain of BET paralogues by exploiting differences in μs–ms conformational dynamics between the three paralogues. Perhaps surprisingly, residues that are key determinants of these dynamic properties are distant from both of the two distinct surfaces that mediate BD-peptide interactions.

## DISCUSSION

### The cyclic peptide 4.1A exploits a single-residue difference on a non-canonical surface to achieve high paralogue-level specificity

Specificity is a fundamental parameter in the development of bioactive molecules, and perhaps the most challenging to achieve. Selectivity between closely related paralogues is a particularly difficult challenge. Though multiple examples exist^43–48^, these typically rely on exploiting one or more amino acid differences that give rise to altered ground-state structures of the paralogues. For example, the EZH2 inhibitor GSK126, which exploits sequence differences within the binding site (only six differences within 10 Å of the binding site) to derive selectivity over the closely related methyltransferases EZH1, or inhibitors of glycogen synthetase kinases that derive paralogue selectivity by exploiting a single-residue difference.

The cyclic peptide 4.1A binds BRD4-BD1 with a selectivity of at least 1,000-fold over the BDs of BRD2 and BRD3. An X-ray crystal structure complemented by mutagenesis revealed that this specificity derives from hydrogen bonds made by 4.1A with a single amino acid – Q123 – that is not conserved in the other BDs (including BRDT). A similar observation has been made in the case of cyclic peptides that can distinguish BCL-2 from BCL-X_L_ by targeting a single residue difference at a critical position in the binding groove^49^. However, 4.1A is unique in that not only does it exploit a single residue difference, but because this difference is located on a surface that is a considerable distance from the orthosteric AcK-binding pocket.

The bivalent binding mode of 4.1A also drives the formation of a fibrillar structure that is observed in the structure of its complex with BRD4-BD1. However, this bivalent architecture is not a necessary condition for high-affinity and selective binding; the Kac9A mutant of 4.1A retains high affinity and selectivity and yet does not bind appreciably to the AcK pocket, meaning that selectivity is driven primarily by the αB-αC surface.

### 2.1C reads differences in the conformational dynamics of BET-family BDs

The BRD2-BD1:2.1C complex displays both striking similarities with and striking differences from the BRD4-BD1:4.1A complex. In both cases, each BD binds two copies of the peptide, one copy via a canonical AcK-pocket interaction and the second via a surface formed by the αB and αC helices. Both 2.1C and 4.1A use two distinct surfaces to make these two interactions, giving rise to polymeric complexes.

However, whereas 4.1A exploits a sequence difference on the αB-αC binding surface to achieve selectivity, 2.1C remarkably appears to distinguish BRD2-BD1 from its paralogues by sensing the lack of μs–ms dynamics in its target. Mutations that introduce such dynamics to BRD2-BD1 weaken its affinity for 2.1C and point mutations in the αC helix of BRD3-BD1 that dampen its μs– ms dynamics endow the domain with robust 2.1C-binding ability. Each of these latter mutations – A128T, S139Q and V145Q – alter the specificity of 2.1C by at least ∼200–500-fold.

Although the idea that protein dynamics can impact ligand affinity – and can do so allosterically – is well-established, the specificity observed here is notable for several reasons. First, none of the three mutations appears to significantly alter the ground-state structure of BRD3-BD1. Second, none of these residues directly contact either copy of the peptide in the complex. Third, these residues do not themselves display observable dynamics on the μs–ms timescale and, fourth, the changes in BRD3-BD1 dynamics that are induced by each mutation occur in many cases a significant distance (up to ∼25 Å) from the mutations. Thus, residues in these three positions act to broadly regulate dynamics across the BD1 domains.

As well as providing insight into the mechanism underlying selective binding, it is likely that the significant differences in μs–ms dynamics between BET family members is also translated to functional differences between the paralogues *in vivo*, especially given that the AcK-binding ZA loop is one focal point of the observed dynamics. For example, out of a total of 425 unique proteins identified as binding partners of BRD2, BRD3 or BRD4 in one study, only 105 were shared between all three proteins^11^. This selectivity could arise in part from the distinct dynamical properties of the paralogues.

### A new binding surface on BET bromodomains

Of the thousands of small molecules and proteins that have been discovered or designed to bind BET BD1 domains, only one experimental structure shows a ligand that binds to a location other than the AcK pocket: a small molecule that bound the ‘outside’ surface of the AcK pocket^50^. However, 2.1C and 4.1A contact a surface formed by the αB and αC helices, and this interaction is essential for the selectivity that each peptide displays. The αB-αC contact surface is highly similar in the two complexes, despite the completely different structures of the two peptides; the only shared contact is a hydrogen bond between a peptide tyrosine and a glutamate conserved in both BDs.

These interactions suggest the possibility that the αB-αC surface might be an as yet unrecognized interaction surface on BET BDs, perhaps even one that drives paralogue-specific interactions *in vivo*. A recently reported Alphafold2-multimer prediction of a complex formed between BRD3-BD1 and the splicing related protein ACIN1 shows ACIN1 packing against the αB-αC surface of the BD – and using a tyrosine (Y140) to form a hydrogen bond to the same glutamate discussed above (E130; **Fig. S19**)^51^. Intriguingly, ACIN1 was previously identified as a BRD2, BRD3, BRD4, BRDT interaction partner from mass spectrometry data and, in the same study, only the interaction with BRD3 was retained following treatment with the BET inhibitor JQ1 (which binds the AcK pocket)^11^. These data hint at a scenario in which (a) each BET protein can bind ACIN1 through the αB-αC surface, and (b) there is allosteric communication between the AcK pocket and the αB-αC surface – such that JQ1 binding abrogates binding of ACIN1 to BRD2, BRD4 and BRDT but retains affinity for BRD3. We speculate that the differences in dynamics that we observe between paralogues could be the basis for such specificity

### New directions for BET inhibition?

Despite intense efforts over the last 15 years, the design of paralogue-specific BET inhibitors has proven a challenge that has only recently begun to be overcome. Our data suggest a new avenue for exploration. Allosteric inhibitors can provide specificity when orthosteric binding sites are highly homologous, and several allosteric small-molecule drugs are already on the market^7^. Molecules that target the BET family BDs beyond the highly conserved AcK-binding pocket might offer enhanced selectivity and allosterically modulate ligand binding, given the clear differences observed between paralogues in ZA-loop dynamics.

As well as targeting a new contact surface for BET modulators, our paralogue-selective cyclic peptides induce filament formation of their target BD. A similar phenomenon has been observed for a small-molecule inhibitor of BCL6 (BI-3802), which acts as a molecular glue to induce the cellular polymerisation and subsequent proteasomal degradation of its target^52^. Cell permeable derivatives of 2.1C and 4.1A might act in a corresponding way, either inducing degradation by the proteosome or simply sequestering the protein to block its normal activity.

Finally, the differences in dynamics that we observe could be exploited in inhibitor design. As noted above, crystal structures of the BRD4-BD2 selective inhibitors described by Li *et al*. did not reveal any features that could explain the selectivity, and it was speculated that differences in binding pocket dynamics might explain their observations^50^. Although high-throughput measurements of protein-ligand dynamics are likely to be a challenge, there is an argument that mid-pipeline analysis of protein dynamics should perhaps be a more routine weapon in the arsenal of drug discovery programs.

## EXPERIMENTAL PROCEDURES

### Protein expression

The wildtype BDs from human BRD2 (BD1: 65–194; BD2: 347–455), human BRD3 (BD1: 25– 147), rat BRD3 (BD2: 307–419), and human BRD4 (BD1: 42–168; BD2: 348–464) were cloned into pGEX-6P for expression as *N*-terminal GST-HRV3C-fusion proteins for NMR, X-ray crystallography, and SEC-MALS and into pQE80L-*N*avi for expression as *N*-terminally His-tagged and Avitagged™ proteins for the production of biotinylated BDs for RaPID screening and SPR as previously described^38^. Mutations to these constructs were introduced using site-directed mutagenesis. Chimeric and mutant BDs used for SPR were cloned into pQE80L-GST-HRV3C-*N*avi for expression as GST-HRV3C-Avitagged™ proteins.

All constructs were transformed into Rosetta2(DE3) *Escherichia coli* (*E*. *coli*) cells in preparation for recombinant protein expression. Saturated starter cultures grown overnight from single colonies from fresh transformation plates were used to inoculate large scale 2xYT expression cultures (1:100 dilution; all media supplemented with the appropriate antibiotics). Expression cultures were incubated at 37 °C, with shaking at 150 rotations per minute (rpm), until an OD_600_ of ∼0.6–0.8 was reached. For proteins used for RaPID, SPR, X-ray crystallography, and SEC-MALS, expression cultures were then transferred to 18 °C, supplemented with 250 μM isopropyl β-D-1-thiogalactopyranoside (IPTG; 200 μM biotin was also added at this point for Avitagged™ BDs), and incubated with shaking at 150 rpm for a further ∼20–24 hours to enable expression. Expression cultures were harvested via centrifugation at 4500 x*g* for 25 minutes and stored at –20 °C.

Uniformly ^15^N/^13^C-labelled proteins used for NMR were produced as above but cultures were harvested via centrifugation (4000 x*g* for 10 minutes) upon reaching an OD_600_ of ∼0.6–0.8. The cell pellets were washed in a small volume M9 minimal media salts and harvested again to remove any residual 2xYT media and finally resuspended in a volume of minimal media (containing ^15^NH_4_Cl and/or ^13^C-glucose) half the volume of the original 2xYT culture. The minimal media cultures were then transferred to 18 °C, with shaking at 150 rpm, for ∼45 minutes before expression was induced with 250 μM IPTG. The cultures were incubated for a further ∼20–24 hours before being harvested.

### Protein purification

#### Purification of His-tagged and Avitagged™ proteins

A procedure consisting of nickel-ion (Ni^2+^) affinity chromatography followed by size exclusion chromatography (SEC) was used to prepare *N*-terminally His-tagged and Avitagged™ proteins. Purification steps and protein purity were assessed by SDS-PAGE and concentrations were determined using *A*_280_ nm measurements. All steps were performed at 4 °C or using pre-chilled buffers.

Cell pellets were resuspended in a buffer consisting of 25 mM HEPES pH 7.2, 500 mM NaCl, 20 mM imidazole, 0.5 mM tris(2-carboxyethyl)phosphine hydrochloride (TCEP), 0.1% (v/v) Triton-X-100, 1 x cOmplete EDTA-free protease inhibitor, 10 μg/mL DNase I, 10 μg/mL RNase I, and 100 μg/mL lysozyme. The resuspended cells were lysed via sonication and the lysate was clarified via centrifugation at 15,000 x*g* for ∼0.5-1 hour. The soluble fraction of the lysate was then applied to Ni^2+^-nitrilotriacetic acid (NTA) resin (Cytiva) preequilibrated in a wash buffer comprising 25 mM HEPES pH 7.2, 500 mM NaCl, 20 mM imidazole, and 0.5 mM TCEP. The resin was then washed with wash buffer and bound proteins were eluted using a buffer comprising 25 mM HEPES pH 7.2, 150 mM NaCl, 300 mM imidazole, and 0.5 mM TCEP. The eluates were concentrated to ∼ 5 mL and subjected to SEC using a HiLoad 16/600 Superdex 75 column (Cytiva) preequilibrated in a SEC buffer containing 25 mM HEPES pH 7.2, 150 mM NaCl, and 0.5 mM TCEP. The protein was eluted using SEC buffer and protein containing fractions were pooled and either aliquoted directly or concentrated beforehand. Protein aliquots were snap frozen using liquid nitrogen and stored at – 80 C.

#### Purification of GST-HRV3C and GST-HRV3C-Avitagged™ proteins

A procedure consisting of GSH-affinity chromatography, on-resin HRV3C cleavage of the tags, and SEC was used to prepare the proteins expressed as GST-HRV3C and GST-HRV3C-Avitagged™ fusions in this study. Purification steps and protein purity were assessed by SDS-PAGE and concentrations were determined using *A*_280_ nm measurements. All steps were performed at 4 °C or using pre-chilled buffers.

Cell pellets were resuspended in a buffer comprising 25 mM HEPES pH 7.2, 500 mM NaCl, 0.5 mM TCEP, 0.1% (v/v) Triton-X-100, 1 x cOmplete EDTA-free protease inhibitor, 10 μg/mL DNase I, 10 μg/mL RNase I, and 100 μg/mL lysozyme. The resuspension was lysed via sonication and then clarified via centrifugation at 15,000 x*g* for ∼0.5-1 hour. The soluble fraction of the lysate was then applied to GSH resin preequilibrated in a wash buffer containing 25 mM HEPES pH 7.2, 500 mM NaCl, and 0.5 mM TCEP. The resin was washed extensively with the wash buffer before being resuspended in ∼5 mL of a buffer containing 25 mM HEPES pH 7.2, 150 mM NaCl, and 0.5 mM TCEP supplemented with HRV3C (∼1:50 ratio of HRV3C:estimated bound protein concentration). The resuspensions were gently rotated overnight before the cleaved proteins were separated from the resin. The cleaved protein was subjected to SEC using a HiLoad 16/600 Superdex 75 column preequilibrated in SEC buffer. The protein was eluted using SEC buffer and protein containing fractions were either aliquoted directly or concentrated prior to aliquoting and snap freezing using liquid nitrogen. Protein aliquots were stored at –80 C

#### RaPID screening

RaPID screens were performed as described previously^38,39^. Briefly, AcK-focussed DNA libraries were constructed by extension reactions and PCR from oligos purchased from Eurofins Genomics K.K. (Japan). Library DNA – TAATACGACTCACTATAGGGTTGAACTTTAAGTAGGAGATATATCCATG(NNK)m=3-7ATG(NNK)n=4-7TGTGGGTCTGGGTCTGGGTCTTAGGTAGGTAGGCGGAAA T4 RNA polymerase was used to transcribe DNA libraries to mRNA and these mRNA libraries were ligated to a puromycin-PEG-DNA splint using T4 RNA ligase. All reactions followed standard reaction conditions. Initial libraries were pooled as follows: (m=3,n=4):(m=4,n=4):(m=4,n=5):(m=5,n=5):(m=5,n=6):(m=6,n=6):(m=6,n=7):(m=7,n=7) = 0.01425:0.45:10:10:7.5:7.5:7.5:7.5

During translation, peptides were initiated with N-(chloroacetoxy)-L-tryptophan (ClAc-L-Trp), which was coupled to tRNAfMetCAU as the cyanomethyl ester using eFx. Elongator methionine codons were reprogrammed to Nε-acetyl-lysine by coupling to tRNAAsnCAU as the 3,5-dinitrobenzyl ester using dFx. Flexizymes were prepared as described previously^53^.

Puromycin-ligated AcK-focussed mRNA libraries were in vitro translated (30 min, 37 °C then 12 min, 25 °C) using a custom transcription/translation mixture containing additional 12.5 μM ClAc-L-Trp-tRNAfMetCAU and 25 μM AcK-tRNAAsnCAU and lacking methionine and 10-formyl-5,6,7,8-tetrahydrofolic acid. First-round translations were performed on a 150 μL scale, with subsequent rounds on a 5 μL scale. Following addition of 200 mM EDTA, pH 8.0 (15 μL) and reverse transcription with M-MLV RTase, RNase H minus (Promega), blocking buffer was added (50 mM HEPES, 150 mM NaCl, 2 mM DTT, 0.1% Tween-20, 0.2% (w/v) acetylated bovine serum albumin, pH 7.5) and libraries were incubated with magnetic streptavidin bead-immobilised bromodomain (Promega) (200 nM, 30 min). Following ice-cold washing (50 mM HEPES, 150 mM NaCl, 2 mM DTT, 0.1% Tween-20, pH 7.5, 3 × 5 min), PCR solution was added (400 µL for round 1, 100 µL for rounds 2-5) and bound peptide-mRNA/DNA hybrids eluted from the beads by heating (95 °C, 5 min). Library enrichment was assessed by quantitative real-time PCR relative to standards and the input DNA library using primers T7g10M_F46 and CGS3-CH.R22. Enriched pools were amplified using the same primers and used as the input DNA for subsequent selection rounds.

T7g10M_F46-TAATACGACTCACTATAGGGTTGAACTTTAAGTAGGAGATATATCC CGS3-CH.R22 – TTTCCGCCTACCTACCTAAGAC

Recovered library DNA from rounds 3-5 was converted into double indexed libraries (Nextera XT indices) which were sequenced on a MiSeq platform (Illumina) using a v3 chip as single 151 cycle reads. DNA sequences were converted to peptide sequences and ranked by total read number (**Tables S1** and **S2**).

### Peptide synthesis

#### General procedure A; Automated Peptide Synthesis (SYRO I peptide synthesizer)

The resin (164 mg, 100 µmol, 0.61 mmol g-1, 1 eq.) was treated with 40 vol.% piperidine (1.6 mL) in DMF for 3 min, drained, and then treated with 20 vol.% piperidine in DMF for 10 min (1.6 mL), drained, and washed with DMF (4 x 1.6 mL). The resin was then treated with a solution of Fmoc-Xaa-OH (400 µmol, 4 eq.) and Oxyma (57 mg, 400 µmol, 4 eq.) in DMF (800 µL), followed by a solution of DIC (63 µL, 400 µmol, 4 eq.) in DMF (800 µL) and shaken at rt for 1 h. The resin was then drained and washed with DMF (4 x 1.6 mL) before being treated with a solution of 5 vol.% Ac_2_O and 10 vol.% iPr_2_NEt in DMF (1.6 mL) for 5 min at rt, drained, washed with DMF (4 x 1.6 mL) and drained.

#### General procedure B; Manual Peptide Synthesis (standard amino acids)

The resin was treated with 20 vol.% piperidine in DMF (5 mL) for 2 x 5 min, filtered, then washed with DMF (5 x 3 mL), CH_2_Cl_2_ (5 x 3 mL) and DMF (5 x 3 mL). The resin was then treated with a solution of Fmoc-Xaa-OH (4 eq.), Oxyma (4 eq.) and DIC (4 eq.) in DMF (0.1 M with respect to resin loading) and shaken at rt for 1 h. The resin was then drained and washed with DMF (5 x 3 mL), CH_2_Cl_2_ (5 x 3 mL) and DMF (5 x 3 mL). The resin was then treated with a solution of 5 vol.% Ac_2_O and 10 vol.% iPr_2_NEt in DMF (2.5 mL) for 5 min at rt before being washed with DMF (5 x 3 mL), CH_2_Cl_2_ (5 x 3 mL) and DMF (5 x 3 mL).

#### General procedure C: Manual cleavage

The resin was thoroughly washed with CH_2_Cl_2_ (5 x 5 mL) before being treated with 90:5:5 v/v/v TFA:triisopropylsilane:H_2_O and shaken at rt for 2 h. The resin was filtered and washed with TFA (2 x 3 mL). The filtrate was concentrated under a stream of nitrogen before addition of diethyl ether (40 mL) to precipitate the peptide. The peptide was pelleted by centrifugation (5 min at 5000 rcf), the ether decanted, and the resulting crude peptide dried under a gentle stream of nitrogen.

#### Preparative chromatography

Preparative and semi-preparative reversed-phase high performance liquid chromatography (HPLC) was performed using a Waters 600E multisolvent delivery system with a Rheodyne 7725i injection valve (5 mL loading loop) with a Waters 500 pump and a Waters 490E programmable wavelength detector operating at 214 nm and 280 nm. Preparative reversed phase HPLC was performed using a Waters X-Bridge® C18 OBDTM super Prep Column (5 µm, 30 x 150 mm) at a flow rate of 30 mL min-1 using a mobile phase of 0.1% TFA in water (solvent A) and 0.1% TFA in MeCN (solvent B) on linear gradients, unless otherwise specified.

#### Semi-preparative chromatography

was performed using a Waters X-Bridge® BEH C18 OBDTM Prep Column (300 Å, 5 µm, 10 x 250 mm) at a flow rate of 4 mL min-1 using a mobile phase of 0.1% TFA in water (solvent A) and 0.1% TFA in MeCN (solvent B) on linear gradients, unless otherwise specified.

#### Analytical Chromatography-Mass Spectrometry Liquid Chromatography-Mass

Spectrometry (LC-MS) was performed on a Shimadzu 2020 UPLC-MS instrument with a Nexera X2 LC-30AD pump, Nexera X2 SPD-M30A UV/Vis diode array detector and a Shimadzu 2020 (ESI) mass spectrometer operating in positive and negative mode. Separations were performed on a Waters Acquity BEH300 1.7 μm, 2.1 x 50 S6 mm (C18) column at a flow rate of 0.6 mL min-1. All separations were performed using a mobile phase of 0.1 vol.% formic acid in water (solvent A) and 0.1 vol.% formic acid in MeCN (solvent B) using a linear gradient of 0 to 100% solvent B over 5 min, unless otherwise specified.

#### Analytical reversed-phase HPLC

was performed on a Waters Acquity UPLC system equipped with a PDA eλ detector (λ = 210 – 400 nm), a sample manager FAN and Quaternary Solvent Manager (H-Class) modules. Separations were performed on a Waters Acquity BEH300 1.7 μm, 2.1 x 50 mm (C18) column at a flow rate of 0.6 mL min-1. All separations were performed using a mobile phase of 0.1% TFA in water (Solvent A) and 0.1% TFA in MeCN (Solvent B) using linear gradients, unless otherwise specified.

#### Surface plasmon resonance

Surface plasmon resonance experiments were performed using a Biacore™ T200 (Cytiva) instrument. Experiments were conducted by immobilising biotinylated BET BDs on a Biacore™ Biotin CAPture chip (Cytiva) with a target density of ∼1000-1500 response units (RU). A running buffer comprising 20 mM HEPES pH 7.5, 150 mM NaCl, and 0.05% (v/v) Tween-20 was used and experiments were conducted at a 50 μL/min flow rate. Binding experiments were performed at 4 °C using single-cycle kinetic mode and fit using a 1:1 binding mode using the Biacore Insight Evaluation Software (Cytiva). The chip was regenerated as per manufacturer’s protocol as required.

### NMR spectroscopy

#### Sequence-specific backbone (H^N^, N, C**^α^**) and C**^β^** resonance assignments

To assign backbone (H^N^, N, C^α^) and C^β^ resonance assignments of BRD2-BD1 R100Y V106I G109N, BRD3-BD1 A128T, BRD3-BD1 S139Q, and BRD3-BD1 V145Q mutants at 298 K, we used the previously reported H^N^, N, C^α^, and C^β^ assignments of BRD2-BD1 (BMRB ID: 50143) and BRD3-BD1 (BMRB ID: 50148) as references. The wild-type backbone H^N^ and N assignments were initially transferred to the mutant [^15^N,^1^H]-HSQC spectra by overlaying them with the WT HSQC spectra in CARA (http://cara.nmr.ch/)^54^. However, significant chemical shift changes in the amide resonances near the mutation sites made it difficult to unambiguously transfer several amide resonances from WT to mutant spectra. Therefore, the sequence-specific assignments of these mutant proteins were further confirmed using 3D HNCA and CBCA(CO)NH spectra. These triple resonance experiments were acquired using non-uniform sampling (NUS) from [U-^13^C,^15^N]-labelled protein samples (400-600 μM) in a HEPES buffer (50 mM HEPES pH 7.2, 100 mM NaCl, 1 mM TCEP) at 298 K on an 800 MHz Bruker spectrometer with a CryoProbe. NUS data were acquired using a Poisson Gap sampling schedule by Topspin (Bruker Biospin) and reconstructed using hsmIST or MaxEnt algorithms in NMRbox (https://nmrbox.nmrhub.org/)^55^.

#### H^N^-CPMG relaxation dispersion data acquisition and analysis at 288 K

H^N^-single quantum (SQ) pseudo 3D CPMG relaxation dispersion (RD) experiments were recorded on the 600 MHz and 800 MHz spectrometers equipped with CryoProbe at 288 K using the pulse sequence as previously reported^56^. For BRD2-BD1 R100Y V106I G109N, BRD3-BD1 A128T, BRD3-BD1 S139Q, and BRD3-BD1 V145Q samples, H^N^-CPMG RD experiments were collected from [U-^13^C,^15^N]-labelled proteins while [U-^15^N]-labelled proteins were used for the BRD2-BD1, BRD3-BD1, and BRD4-BD1. V_CPMG_ (Hz) values were set between set between 50 - 3000 Hz (T_CPMG_ = 20 ms) with a recycle delay (d1) of 2.5 s. Because these proteins are not perdeuterated, H^N^-CPMG RD data were acquired with a REBURP pulse that selectively inverts the amides without affecting H^α^, as described previously. Before acquiring the relaxation dispersion experiments, H^N^ CPMG and REBURP pulses were manually calibrated for each protein sample. All spectra were acquired using 5 mm Shigemi NMR tubes.

For the H^N^-CPMG RD data analysis of wildtype BDs, we used the previously reported backbone amide resonance assignments from BMRB (BRD2-BD1 – accession number 50143; BRD3-BD1 – accession number 50148 and BRD4-BD1 – accession number 50145). However, we extended the following backbone amide resonance assignments for residues Val63, Asp64, Asn92 and Ala98 of BRD3-BD1 using our previous triple resonance spectra before analyzing the RD data.

To analyse H^N^-CPMG RD data at 288 K, we needed chemical shift assignments at that temperature. Therefore, high-resolution 2D [^15^N,^1^H]-HSQCs of each protein were recorded with 256 complex points in the nitrogen dimension at 288 K, 293 K, and 298 K on an 800 MHz spectrometer. This enabled the transfer of backbone amide resonance assignments from 298 K to 288 K. The spectra were processed using Topspin and overlaid in CARA.

H^N^-CPMG RD data were analyzed using the nmrDraw software package in NMRbox^55^, and the backbone amide cross peak intensities in [^15^N,^1^H]-HSQC spectrum for each V were calculated by PINT^57,58^. R_ex_ values were obtained by subtracting the *R_2,eff_*differences between the lowest (50 Hz) and highest CPMG frequencies (3000 Hz). Chemical exchange parameters (*k_ex_*, *p_b_*and δω) from relaxation dispersion data were extracted using ChemEx (https:// github.com/gbouvignies/ChemEx). The dispersion data were fitted to a two-state exchange model.

### HSQC titrations

Spectra were collected at 298 K using Bruker Avance III 600-and 800-MHz NMR spectrometers fitted with TCI probe heads and using standard pulse sequences from the Bruker library. Topspin and NMRFAM-SPARK were used to analyse spectra NMR spectra were acquired at 298 K using Bruker Avance III 600-or 800-MHz NMR spectrometers fitted with TCI probe heads and using standard pulse sequences from the Bruker library. TOPSPIN3 (Bruker) and NMRFAM-SPARKY. Spectra were internally referenced to 10 μM 4,4-dimethyl-4-silapentane-1 sulfonic acid (DSS). Binding experiments were performed by collecting ^15^N-HSQC spectra of labelled BDs before and after titration of unlabelled peptide into the samples. Interactions were assessed by monitoring the CSPs induced by addition of the peptides to the BDs.

### Secondary shift analysis

The secondary chemical shift values were calculated using UNIO software^59^. The first column shows the residue number. The second and third columns show the difference between the experimental Cα/Cβ chemical shifts and the corresponding random coil shifts. The last column shows the Cα-Cβ value for each residue, which is calculated as the average over three consecutive residues (i-1, i, and i+1), following the method described by Metzler *et al*^60^. A positive Cα-Cβ value of a residue indicates that it is a part of a helix, while a negative value indicates it belongs to a strand. A stretch of three or more neighbouring residues with a Cα-Cβ ≥ 1 ppm is considered as evidence of a regular secondary structure.

### X-ray crystallography

Crystallisation experiments for BD-peptide complexes were initially performed using commercial 96-well crystallisation screens (PACT, JCSG+, and Morpheus I from Molecular Dimensions and Index, Crystal Screen, and PEGRx from Hampton Research) in a sitting-drop vapour diffusion format at 18 °C. BDs (10–15 mg/mL) were either mixed with ∼1.5 molar equivalents of peptide directly or diluted to a ∼1 mg/mL concentration before the addition of ∼1.5 molar equivalents of peptide and then concentrated back to ∼10 mg/mL. BD-peptide complexes were dispensed into MRC two-drop chamber crystallisation plates using a Mosquito crystallisation robot and each condition was screened at a 1:1 or 2:1 protein:precipitant ratio (maintaining a final drop volume of 300 nL). In cases where heavy precipitation was observed when peptide was added to BDs, crystallisation was performed by separately dispensing the BD and peptide directly into the crystallisation plate prior to mixing with the precipitant. Microseeding and gradient refinement were used to optimise initial hits where required. Protein crystals generally took days to several weeks to appear. Crystals were cryoprotected using ∼10% glycerol in the mother liquid from which the crystals grew, harvested, and plunge frozen in liquid nitrogen.

X-ray diffraction data were collected at the Australian Synchrotron (AS) using the Macromolecular Crystallography MX1 (bending magnet) and MX2 (microfocus) beamlines at 100 K and a wavelength of 0.9537 Å^61,62^. The data were processed using the automated XDS pipeline provided by the AS and AIMLESS from the CCP4i suite was used to scale the data^63,64^. The initial phases were calculated using the molecular replacement program PhaserMR within the Phenix suite using existing structures of the BET BDs as MR models (PDB IDs 4UYF for BRD2 BD1, 3ONI for BRD2-BD2, 3S91 for BRD3-BD1, 3S92 for BRD3-BD2, 4LYI for BRD4-BD1, and 5UVV for BRD4-BD2)^65–69^. The models were refined using consecutive rounds of manual model building in COOT followed by Phenix refine^70^. The wwPDB server was used to assess the quality of the final model before being submitted to the PDB. Structure diagrams were generated using PyMOL. The data collection and refinement statistics for all structures described in this study are outlined in **Tables S4 and 5**.

### SEC-MALS

Molecular weight analyses were performed by subjecting samples to SEC coupled with multi-angle light scatter. Proteins of complexes (∼100 μL at ∼100–300 μM concentrations) were subject to to SEC (using a Superdex 75 10/300 Increase) on an Äkta system with inline multi-angle light scattering (MALS), UV absorbance, and refractive index (dRI) detectors. MALS, UV, and dRI data were collected and analysed using the ASTRA software (Wyatt) and molecular weights were determined using the Debye-Zimm model. BSA (100 µL of 2 mg/mL) was run as a standard to calibrate the MALS, UV, and dRI signals. We estimate the uncertainty in the molecular weight determination from this system to be ±10%.

## DATA AVAILABILITY

The chemical shift assignments for the BRD2-BD1 R100Y V106I G109N (BMRB ID: 52788), BRD3-BD1 A128T (BMRB ID: 52785), BRD3-BD1 Q139S (BMRB ID: 52786), and BRD3-BD1 V145Q (BMRB ID: 52787) mutants have been deposited to the Biological Magnetic Resonance Bank (BMRB). The coordinates the structure factors for all structures described in this study have been deposited to the Protein Data Bank (PDB). The PDB IDs are as follows: 9MPI (BRD4-BD1:4.1A), 9MPJ (BRD2-BD1:2.1C), 9MPL (BRD2-BD1:2.1C W11A), 9MPM (BRD3-BD1:2.1C W11A), and 9MPN (BRD2-BD1 A128T).

## SUPPORTING INFORMATION

This article contains supporting information provided as separate documents

## ACKNOWLEDGEMENTS

This research was undertaken using the MX1 and MX2 beamlines at the Australian Synchrotron, part of Australian Nuclear Science and Technology Organization, and made use of the Australian Cancer Research Foundation Eiger 16M detector. We acknowledge the Sydney Analytical Core Facilities (University of Sydney) for providing access to NMR and SPR infrastructure. J.P.M., L.J.W., and R.J.P. received funding from the National Health and Medical Research Council (APP 1161623). L.J.W. received funding from the European Union’s Horizon 2020 research and innovation program under Marie Skłodowska-Curie Grant Agreement 657292. This work is also supported by The Francis Crick Institute which receives its core funding from Cancer Research UK (CC2030), the UK Medical Research Council (CC2030), and the Wellcome Trust (CC2030). K.P. is supported by a European Molecular Biology Organisation (EMBO) Postdoctoral Fellowship (ALTF 733-2024).

## AUTHOR CONTRIBUTIONS

Conceptualisation: J.P.M., L.J.W., K.P., and B.M. Methodology: K.P., B.M., A.N., J.K.K.L., P.P., and L.W. Investigation, Formal Analysis, and Validation: K.P., B.M., A.N., C.F., P.P., L.Y., X.J., X.J.R., J.L.W., D.H.T., D.F., L.W., L.J.W., and J.P.M. Resources: A.N., D.H.T., D.F., and R.J.P.

Writing – Original Draft: J.P.M., L.J.W., K.P., and B.M. Writing – Review & Editing: J.P.M., L.J.W., K.P., and B.M. with input from all authors. Visualisation: K.P. Supervision: J.P.M., L.J.W., T.P., R.J.P., and H.S. Funding Acquisition: J.P.M., L.J.W., and R.J.P.

## REFERENCES

1. Campillos, M., Kuhn, M., Gavin, A. C., Jensen, L. J. & Bork, P. Drug target identification using side-effect similarity. Science (1979) 321, 263–266 (2008).

2. Huggins, D. J., Sherman, W. & Tidor, B. Rational approaches to improving selectivity in drug design. J Med Chem 55, 1424–1444 (2012).

3. Niwa, T. Elucidation of characteristic structural features of ligand binding sites of protein kinases: A neural network approach. J Chem Inf Model 46, 2158–2166 (2006).

4. Marmorstein, R. Structure of Histone Deacetylases: Insights into Substrate Recognition and Catalysis. Structure 9, 1127–1133 (2001).

5. Han, B., Salituro, F. G. & Blanco, M. J. Impact of Allosteric Modulation in Drug Discovery: Innovation in Emerging Chemical Modalities. ACS Med Chem Lett 11, 1810–1819 (2020).

6. Tee, W. V. & Berezovsky, I. N. Allosteric drugs: New principles and design approaches. Curr Opin Struct Biol 84, 102758 (2024).

7. Lu, S., He, X., Ni, D. & Zhang, J. Allosteric Modulator Discovery: From Serendipity to Structure-Based Design. J Med Chem 62, 6405–6421 (2019).

8. Taniguchi, Y. The Bromodomain and Extra-Terminal Domain (BET) Family: Functional Anatomy of BET Paralogous Proteins. Int J Mol Sci 17, (2016).

9. Morinière, J. et al. Cooperative binding of two acetylation marks on a histone tail by a single bromodomain. Nature 461, 664–668 (2009).

10. Wu, S. Y. & Chiang, C. M. The double bromodomain-containing chromatin adaptor Brd4 and transcriptional regulation. Journal of Biological Chemistry 282, 13141–13145 (2007).

11. Lambert, J.-P. et al. Interactome Rewiring Following Pharmacological Targeting of BET Bromodomains. Mol Cell 73, 621–638 (2019).

12. Patel, K. et al. BET-Family Bromodomains Can Recognize Diacetylated Sequences from Transcription Factors Using a Conserved Mechanism. Biochemistry 60, 648–662 (2021).

13. French, C. a et al. BRD4-NUT Fusion Oncogene: A Novel Mechanism in Aggressive Carcinoma Advances in Brief BRD4-NUT Fusion Oncogene: A Novel Mechanism in Aggressive Carcinoma 1. Cancer Res 63, 304–307 (2003).

14. Denis, G. V. Bromodomain coactivators in cancer, obesity, type 2 diabetes, and inflammation. Discovery medicine vol. 10 489–499 Preprint at (2010).

15. Cheung, K. L., Kim, C. & Zhou, M. M. The Functions of BET Proteins in Gene Transcription of Biology and Diseases. Front Mol Biosci 8, 787 (2021).

16. Belkina, A. C. & Denis, G. V. BET domain co-regulators in obesity, inflammation and cancer. Nat Rev Cancer 12, 465–477 (2012).

17. Padmanabhan, B., Mathur, S., Manjula, R. & Tripathi, S. Bromodomain and extra-terminal (BET) family proteins: New therapeutic targets in major diseases. J Biosci 41, 295–311 (2016).

18. Dawson, M. A. et al. Inhibition of BET recruitment to chromatin as an effective treatment for MLL-fusion leukaemia. Nature 478, 529–533 (2011).

19. Delmore, J. E. et al. BET bromodomain inhibition as a therapeutic strategy to target c-Myc. Cell 146, 904–917 (2011).

20. Denis, A. C., Belkina, B. S. & Nikolajczyk, G. V. Macrophage Inflammatory Responses BET Inhibitor JQ1 Impair Mouse Inflammation: Brd2 Genetic Disruption and BET Protein Function Is Required for. (2013) doi:10.4049/jimmunol.1202838.

21. Liu, L., Yang, C. & Candelario-Jalil, E. Role of BET Proteins in Inflammation and CNS Diseases. Front Mol Biosci 8, 906 (2021).

22. Borck, P. C., Guo, L. W. & Plutzky, J. BET Epigenetic Reader Proteins in Cardiovascular Transcriptional Programs. Circ Res 1190–1208 (2020) doi:10.1161/CIRCRESAHA.120.315929.

23. Mita, M. M. & Mita, A. C. Bromodomain inhibitors a decade later: a promise unfulfilled? British Journal of Cancer 2020 123:12 123, 1713–1714 (2020).

24. Collins, M. K., Chau, C. H., Price, D. K. & Figg, W. D. BETting on next-generation bromodomain inhibitors. Am J Clin Exp Urol 8, 129 (2020).

25. Mertz, J. A. et al. Targeting MYC dependence in cancer by inhibiting BET bromodomains. Proc Natl Acad Sci U S A 108, 16669–16674 (2011).

26. Fong, C. Y. et al. BET inhibitor resistance emerges from leukaemia stem cells. Nature 525, 538–542 (2015).

27. Filippakopoulos, P. et al. Selective inhibition of BET bromodomains. Nature 468, 1067–1073 (2010).

28. Tanaka, M. et al. Design and characterization of bivalent BET inhibitors. Nat Chem Biol 12, 1089–1096 (2016).

29. Waring, M. J. et al. Potent and selective bivalent inhibitors of BET bromodomains. Nat Chem Biol 12, 1097–1104 (2016).

30. Andrieu, G., Belkina, A. C. & Denis, G. V. Clinical trials for BET inhibitors run ahead of the science. Drug Discov Today Technol 19, 45–50 (2016).

31. Amorim, S. et al. Bromodomain inhibitor OTX015 in patients with lymphoma or multiple myeloma: a dose-escalation, open-label, pharmacokinetic, phase 1 study. Lancet Haematol 3, e196–e204 (2016).

32. Postel-Vinay, S. et al. First-in-human phase I study of the bromodomain and extraterminal motif inhibitor BAY 1238097: emerging pharmacokinetic/pharmacodynamic relationship and early termination due to unexpected toxicity. Eur J Cancer 109, 103–110 (2019).

33. Falchook, G. et al. Development of 2 Bromodomain and Extraterminal Inhibitors With Distinct Pharmacokinetic and Pharmacodynamic Profiles for the Treatment of Advanced Malignancies. Clin Cancer Res 26, 1247–1257 (2020).

34. Cui, H. et al. A Structure-based Design Approach for Generating High Affinity BRD4 D1-Selective Chemical Probes. J Med Chem 65, 2342–2360 (2022).

35. Liu, Z. et al. Discovery, X-ray Crystallography, and Anti-inflammatory Activity of Bromodomain-containing Protein 4 (BRD4) BD1 Inhibitors Targeting a Distinct New Binding Site. J Med Chem 65, 2388–2408 (2022).

36. Liu, Z. et al. Discovery of Orally Bioavailable Chromone Derivatives as Potent and Selective BRD4 Inhibitors: Scaffold Hopping, Optimization, and Pharmacological Evaluation. J Med Chem 63, 5242–5256 (2020).

37. Liu, Z. et al. Discovery of potent and selective BRD4 inhibitors capable of blocking TLR3-induced acute airway inflammation. Eur J Med Chem 151, 450–461 (2018).

38. Patel, K. et al. Cyclic peptides can engage a single binding pocket through highly divergent modes. Proc Natl Acad Sci U S A 117, 26728–26738 (2020).

39. Franck, C. et al. Discovery and characterization of cyclic peptides selective for the C-terminal bromodomains of BET family proteins. Structure 31, 912–923.e4 (2023).

40. Hurd, C. A., Bush, J. T., Powell, A. J. & Walport, L. J. mRNA Display in Cell Lysates Enables Identification of Cyclic Peptides Targeting the BRD3 Extraterminal Domain. Angewandte Chemie International Edition 63, e202406414 (2024).

41. Skrynnikov, N. R., Mulder, F. A. A., Hon, B., Dahlquist, F. W. & Kay, L. E. Probing slow time scale dynamics at methyl-containing side chains in proteins by relaxation dispersion NMR measurements: Application to methionine residues in a cavity mutant of T4 lysozyme. J Am Chem Soc 123, 4556–4566 (2001).

42. Loria, J. P., Rance, M. & Palmer, A. G. A relaxation-compensated Carr-Purcell-Meiboom-Gill sequence for characterizing chemical exchange by NMR spectroscopy [13]. J Am Chem Soc 121, 2331–2332 (1999).

43. Brand, M. et al. Homolog-Selective Degradation as a Strategy to Probe the Function of CDK6 in AML. Cell Chem Biol 26, 300–306.e9 (2019).

44. Chen, X. et al. Paralogue-Selective Degradation of the Lysine Acetyltransferase EP300. JACS Au 4, 3094–3103 (2024).

45. Gewirth, D. T. Paralog Specific Hsp90 Inhibitors – A Brief History and a Bright Future. Curr Top Med Chem 16, 2779–2791 (2016).

46. Dvorak, V. et al. Paralog-dependent isogenic cell assay cascade generates highly selective SLC16A3 inhibitors. Cell Chem Biol 30, 953–964.e9 (2023).

47. McCabe, M. T. et al. EZH2 inhibition as a therapeutic strategy for lymphoma with EZH2-activating mutations. Nature 2012 492:7427 492, 108–112 (2012).

48. Miller, M. S., Thompson, P. E. & Gabelli, S. B. Structural Determinants of Isoform Selectivity in PI3K Inhibitors. Biomolecules 2019, Vol. 9, Page 82 9, 82 (2019).

49. Li, F. et al. Cyclic peptides discriminate BCL-2 and its clinical mutants from BCL-XL by engaging a single-residue discrepancy. Nature Communications 2024 15:1 15, 1–17 (2024).

50. Liu, Z. et al. Discovery, X-ray Crystallography, and Anti-inflammatory Activity of Bromodomain-containing Protein 4 (BRD4) BD1 Inhibitors Targeting a Distinct New Binding Site. J Med Chem 65, 2388–2408 (2022).

51. Zhang, J. et al. Computing the Human Interactome. bioRxiv 2024.10.01.615885 (2024) doi:10.1101/2024.10.01.615885.

52. Słabicki, M. et al. Small-molecule-induced polymerization triggers degradation of BCL6. Nature 2020 588:7836 588, 164–168 (2020).

53. Goto, Y., Goto, Y., Katoh, T. & Suga, H. Preparation of materials for flexizyme reactions and genetic code reprogramming. Protoc Exch (2011) doi:10.1038/protex.2011.209.

54. Keller, R. L. J. The Computer Aided Resonance Assignment Tutorial. (2004).

55. Maciejewski, M. W. et al. NMRbox: A Resource for Biomolecular NMR Computation. Biophys J 112, 1529–1534 (2017).

56. Yuwen, T. & Kay, L. E. Revisiting 1HN CPMG relaxation dispersion experiments: a simple modification can eliminate large artifacts. J Biomol NMR 73, 641–650 (2019).

57. Ahlner, A., Carlsson, M., Jonsson, B. H. & Lundström, P. PINT: A software for integration of peak volumes and extraction of relaxation rates. J Biomol NMR 56, 191–202 (2013).

58. Niklasson, M. et al. Comprehensive analysis of NMR data using advanced line shape fitting. J Biomol NMR 69, 93–99 (2017).

59. Guerry, P. & Herrmann, T. Comprehensive Automation for NMR Structure Determination of Proteins. Methods in Molecular Biology 831, 429–451 (2012).

60. Metzler, W. J. et al. Characterization of the three-dimensional solution structure of human profilin: proton, carbon-13, and nitrogen-15 NMR assignments and global folding pattern. Biochemistry 32, 13818–13829 (2002).

61. Cowieson, N. P. et al. MX1: A bending-magnet crystallography beamline serving both chemical and macromolecular crystallography communities at the Australian Synchrotron. J Synchrotron Radiat 22, 187–190 (2015).

62. Aragão, D. et al. MX2: a high-flux undulator microfocus beamline serving both the chemical and macromolecular crystallography communities at the Australian Synchrotron. J Synchrotron Radiat 25, 885–891 (2018).

63. Winn, M. D. et al. Overview of the CCP4 suite and current developments. Acta Crystallographica Section D: Biological Crystallography vol. 67 235–242 Preprint at 10.1107/S0907444910045749 (2011).

64. Potterton, E., Briggs, P., Turkenburg, M. & Dodson, E. A graphical user interface to the CCP4 program suite. Acta Crystallogr D Biol Crystallogr 59, 1131–1137 (2003).

65. Adams, P. D. et al. PHENIX: A comprehensive Python-based system for macromolecular structure solution. Acta Crystallogr D Biol Crystallogr 66, 213–221 (2010).

66. Zwart, P. H. et al. Automated structure solution with the PHENIX suite. Methods Mol Biol 426, 419–435 (2008).

67. Lucas, X. et al. 4-Acyl pyrroles: Mimicking acetylated lysines in histone code reading. Angewandte Chemie - International Edition 52, 14055–14059 (2013).

68. Filippakopoulos, P. et al. Histone recognition and large-scale structural analysis of the human bromodomain family. Cell 149, 214–231 (2012).

69. Gosmini, R. et al. The discovery of I-BET726 (GSK1324726A), a potent tetrahydroquinoline ApoA1 up-regulator and selective BET bromodomain inhibitor. J Med Chem 57, 8111–8131 (2014).

70. Emsley, P. & Cowtan, K. Coot: Model-building tools for molecular graphics. Acta Crystallogr D Biol Crystallogr 60, 2126–2132 (2004).

