## Supporting Information for "Peptide molecular glues select between BET paralogues by exploiting allosteric sites and conformational dynamics"

**LIST OF MATERIALS INCLUDED**

**INCLUDED IN THIS DOCUMENT:**

- Supporting Methodology
- Supporting Tables S3–S9
- Supporting Figures S1–S19

**PROVIDED AS SEPARATE FILES:**

**Table S1.** Enriched sequences from the final three rounds (3–5) of RaPID screening against BRD2-BD1.

**Table S2.** Enriched sequences from the final three rounds (3–5) of RaPID screening against BRD4-BD1.

**METHODOLOGY**

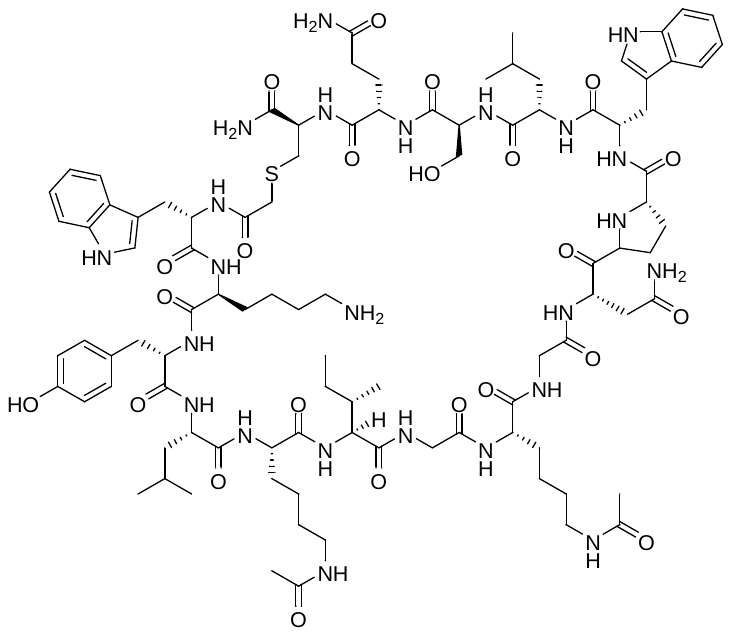

**2.1A** was synthesised via Fmoc-strategy SPPS as specified in general methods. The linear sequence *N’*- WKYLK(Ac)IGK(Ac)GNPWLSQC -*C’* was generated by automated SPPS on Rink amide resin (86 mg, 50µmol, capacity: 0.58 mmolg^-1^). Coupling of chloroacetic acid to the N-terminus, cleavage from resin and cyclisation were performed as described in the general methods. The crude cyclic peptide was purified by semi-preparative RP-HPLC (0 to 40 % B + 0.1 % formic acid over 40 min). The appropriate fractions were combined and lyophilised to afford **2.1A** as a white solid **UPLC:** *R_t_* = 6.46 min. (0 to 60 vol.% B over 5 min, 0.1 vol.% TFA, λ = 214 nm). **LRMS** (ESI+): *m/z =* 1022.8 [M + 2H]^+^.

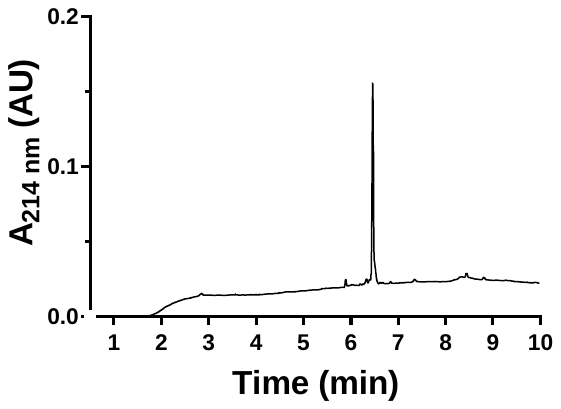

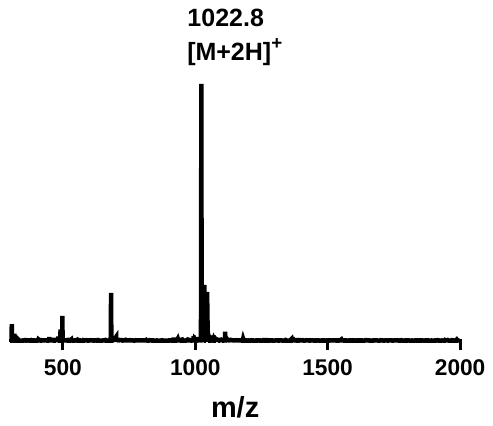

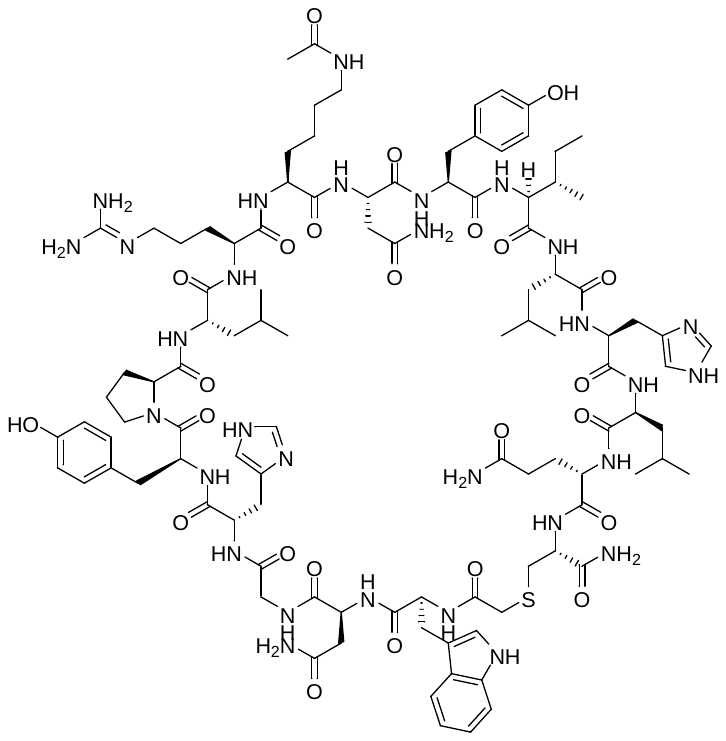

**2.1B** was synthesised via Fmoc-strategy SPPS as specified in general methods. The linear sequence *N’*- WNGHYPLRK(Ac)NYILHLQC -*C’* was generated by automated SPPS on Rink amide resin (86 mg, 50µmol, capacity: 0.58 mmolg^-1^). Coupling of chloroacetic acid to the N-terminus, cleavage from resin and cyclisation were performed as described in the general methods. The crude cyclic peptide was purified by semi-preparative RP-HPLC (0 to 20 % B + 0.1 % formic acid over 40 min). The appropriate fractions were combined and lyophilised to afford **2.1B** as a white solid. **UPLC:** *R_t_* = 7.1 min. (0 to 60 vol.% B over 5 min, 0.1 vol.% TFA, λ = 214 nm). **LRMS** (ESI+): *m/z =* 1747.4 [M + 2H]^+^.

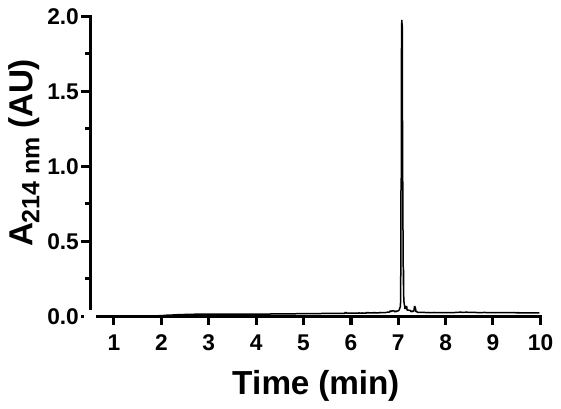

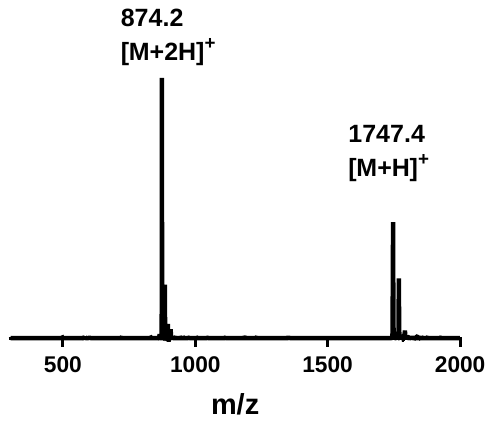

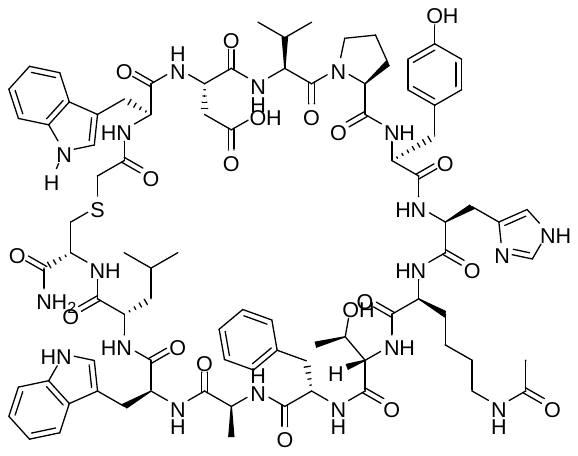

**2.1C** was synthesised via Fmoc-strategy SPPS as specified in general methods. The linear sequence *N’*- WDVPYHK(Ac)TFAWLC -*C’* was generated by automated SPPS on Rink amide resin (86 mg, 50µmol, capacity: 0.58 mmolg^-1^). Coupling of chloroacetic acid to the N-terminus, cleavage from resin and cyclisation were performed as described in the general methods. The crude cyclic peptide was purified by semi-preparative RP-HPLC (0 to 20 % B + 0.1 % formic acid over 40 min). The appropriate fractions were combined and lyophilised to afford **2.1C** as a white solid. **UPLC:** *R_t_* = 7.11 min. (0 to 60 vol.% B over 5 min, 0.1 vol.% TFA, λ = 214 nm). **LRMS** (ESI+): *m/z =* 1747.6 [M + H]^+^.

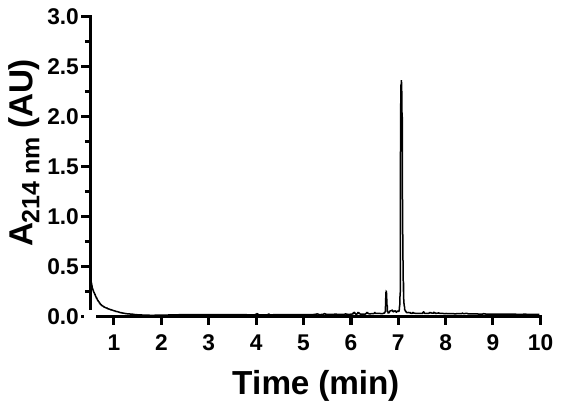

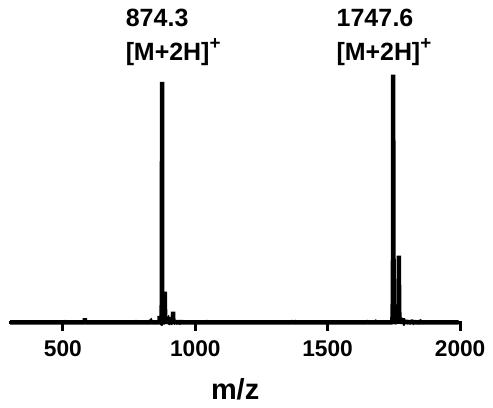

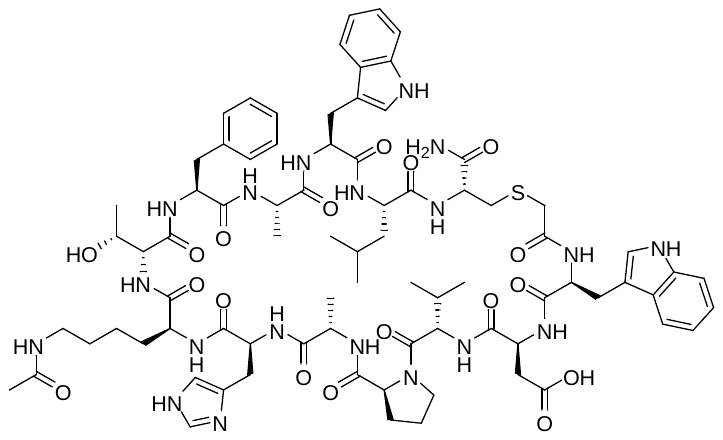

**2.1C Y5A** was synthesised via Fmoc-strategy SPPS as specified in general methods. The linear sequence *N’*- WDVPAHK(Ac)TFAWLC -*C’* was generated by automated SPPS on Rink amide resin (86 mg, 50µmol, capacity: 0.58 mmolg^-1^). Coupling of chloroacetic acid to the N-terminus, cleavage from resin and cyclisation were performed as described in the general methods. The crude cyclic peptide was purified by semi-preparative RP-HPLC (0 to 20 % B + 0.1 % formic acid over 40 min). The appropriate fractions were combined and lyophilised to afford **2.1C_Y5A** as a white solid **UPLC:** *R_t_* = 6.96 min. (0 to 60 vol.% B over 5 min, 0.1 vol.% TFA, λ = 214 nm). **LRMS** (ESI+): *m/z =* 1656.0 [M + H]^+^.

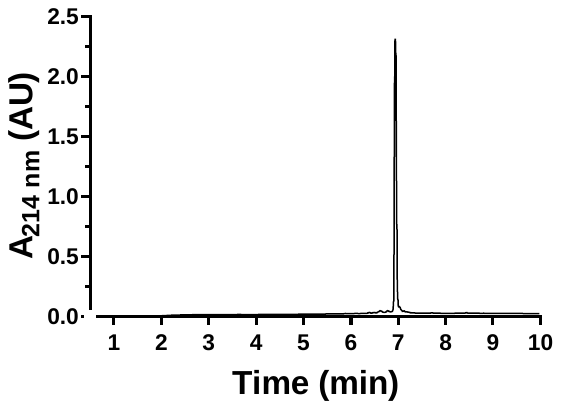

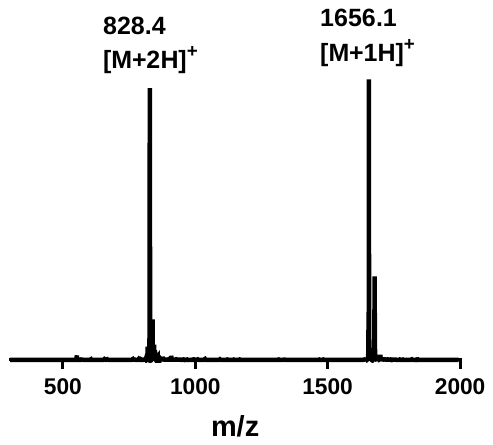

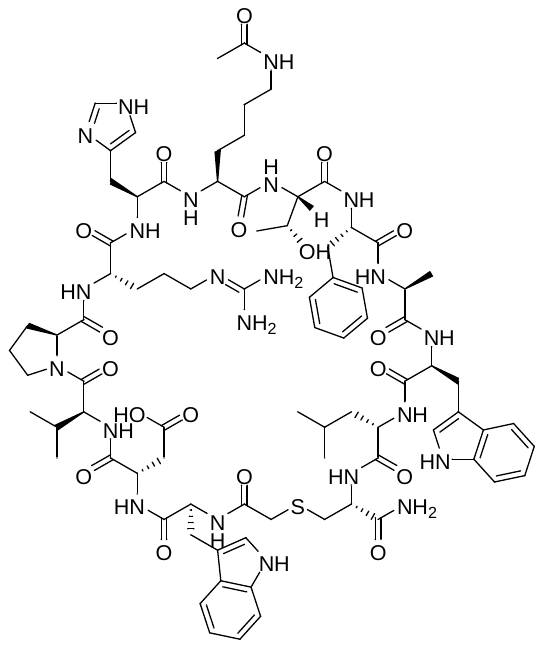

**2.1C Y5R** was synthesised via Fmoc-strategy SPPS as specified in general methods. The linear sequence *N’*- 1 WDVPRHK(Ac)TFAWLC -*C’* was generated by automated SPPS on Rink amide resin (86 mg, 50µmol, capacity: 0.58 mmolg^-1^). Coupling of chloroacetic acid to the N-terminus, cleavage from resin and cyclisation were performed as described in the general methods. The crude cyclic peptide was purified by semi-preparative RP-HPLC (0 to 20 % B + 0.1 % formic acid over 40 min). The appropriate fractions were combined and lyophilised to afford **2.1C_Y5R** as a white solid. **UPLC:** *R_t_* = 7.21 min. (0 to 60 vol.% B over 5 min, 0.1 vol.% TFA, λ = 214 nm). **LRMS** (ESI+): *m/z =* 1741.1 [M + H]^+^.

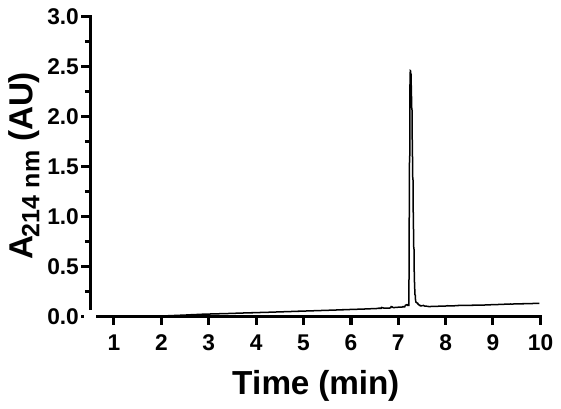

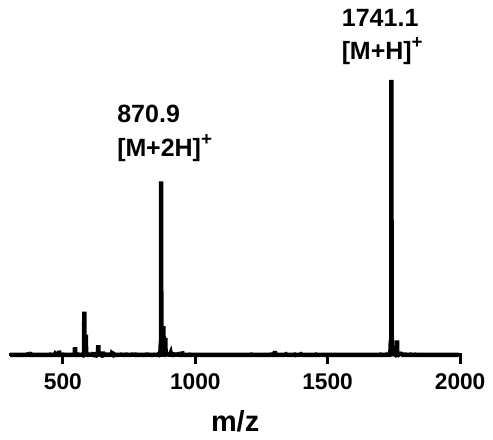

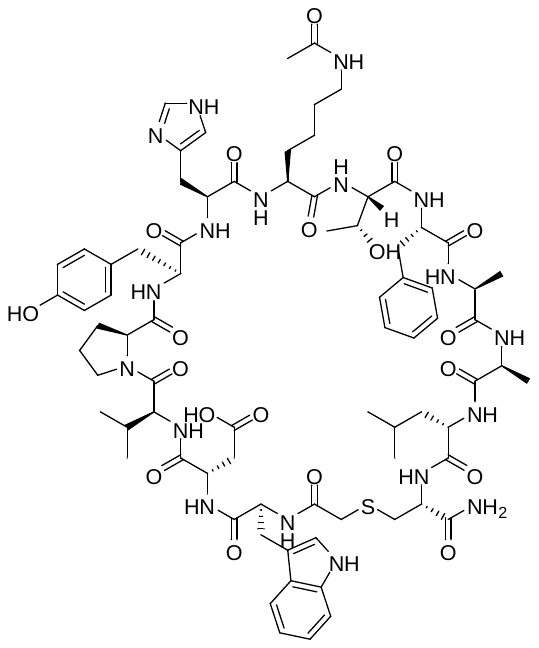

**2.1C W11A** was synthesised via Fmoc-strategy SPPS as specified in general methods. The linear sequence *N’*- WDVPYHK(Ac)TFAALC -*C’* was generated by automated SPPS on Rink amide resin (86 mg, 50µmol, capacity: 0.58 mmolg^-1^). Coupling of chloroacetic acid to the N-terminus, cleavage from resin and cyclisation were performed as described in the general methods. The crude cyclic peptide was purified by semi-preparative RP-HPLC (0 to 20 % B + 0.1 % formic acid over 40 min). The appropriate fractions were combined and lyophilised to afford **2.1C_W11A** as a white solid. **UPLC:** *R_t_* = 7.01 min. (0 to 60 vol.% B over 5 min, 0.1 vol.% TFA, λ = 214 nm). **LRMS** (ESI+): *m/z =* 1633.2 [M + H]^+^.

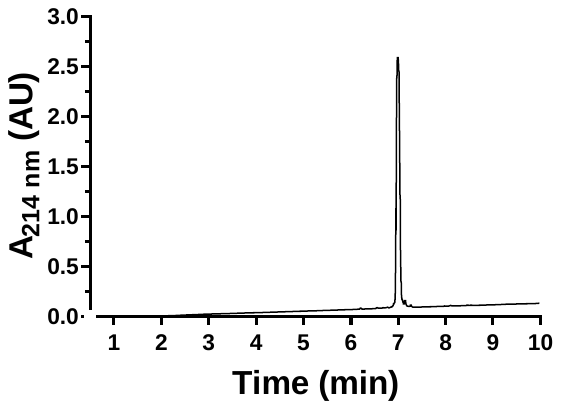

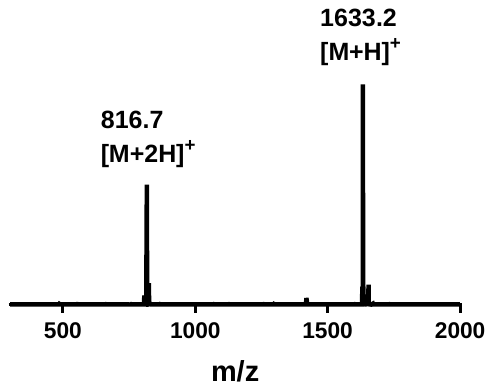

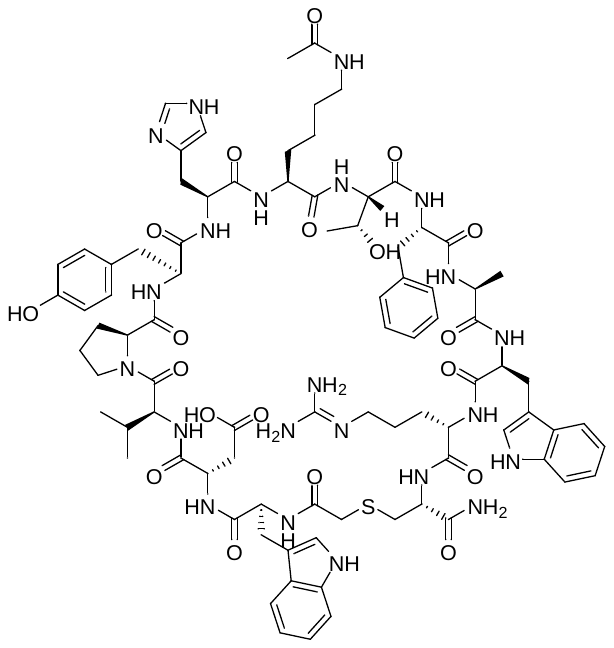

**2.1C L12R** was synthesised via Fmoc-strategy SPPS as specified in general methods. The linear sequence *N’*-WDVPYHK(Ac)TFAWRC -*C’* was generated by automated SPPS on Rink amide resin (86 mg, 50µmol, capacity: 0.58 mmolg^-1^). Coupling of chloroacetic acid to the N-terminus, cleavage from resin and cyclisation were performed as described in the general methods. The crude cyclic peptide was purified by semi-preparative RP-HPLC (0 to 20 % B + 0.1 % formic acid over 40 min). The appropriate fractions were combined and lyophilised to afford **2.1C_L12R** as a white solid. **UPLC:** *R_t_* = 6.65 min. (0 to 60 vol.% B over 5 min, 0.1 vol.% TFA, λ = 214 nm). **LRMS** (ESI+): *m/z =* 895.6 [M + 2H]^+^.

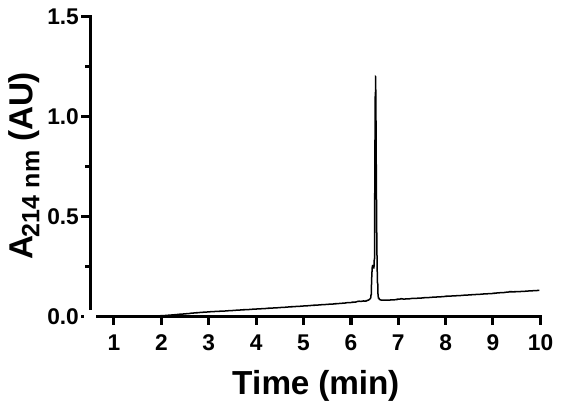

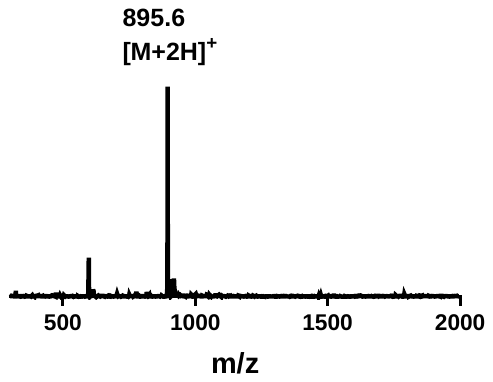

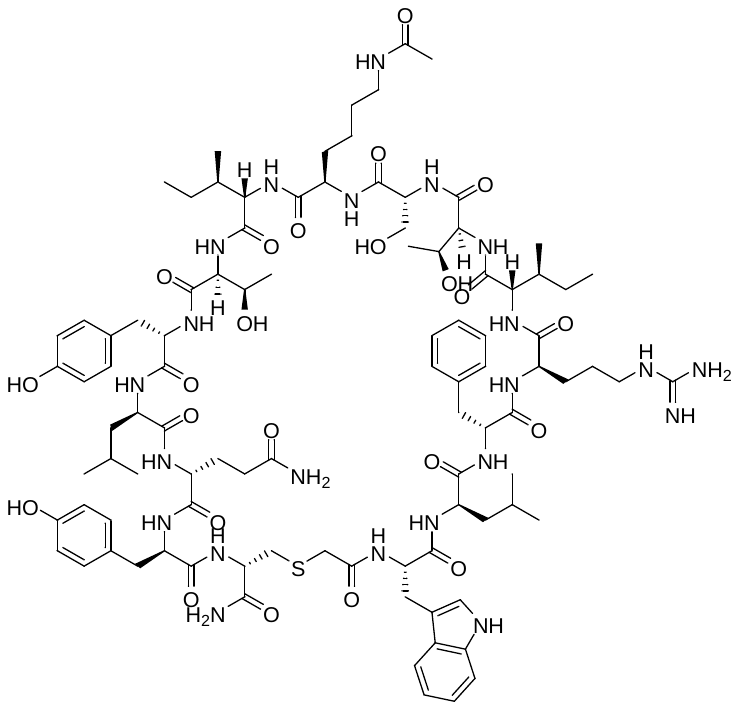

**2.1D** was synthesised via Fmoc-strategy SPPS as specified in general methods. The linear sequence *N’*- WLFRITSK(Ac)ITFLQYC -*C’* was generated by automated SPPS on Rink amide resin (86 mg, 50µmol, capacity: 0.58 mmolg^-1^). Coupling of chloroacetic acid to the N-terminus, cleavage from resin and cyclisation were performed as described in the general methods. The crude cyclic peptide was purified by semi-preparative RP-HPLC (0 to 20 % B + 0.1 % formic acid over 40 min). The appropriate fractions were combined and lyophilised to afford **2.1D** as a white solid. UPLC: *R_t_* = 5.35 min. (0 to 60 vol.% B over 5 min, 0.1 vol.% TFA, λ = 214 nm). LRMS (ESI+): *m/z =* 1009.0 [M + 2H]^2+^.

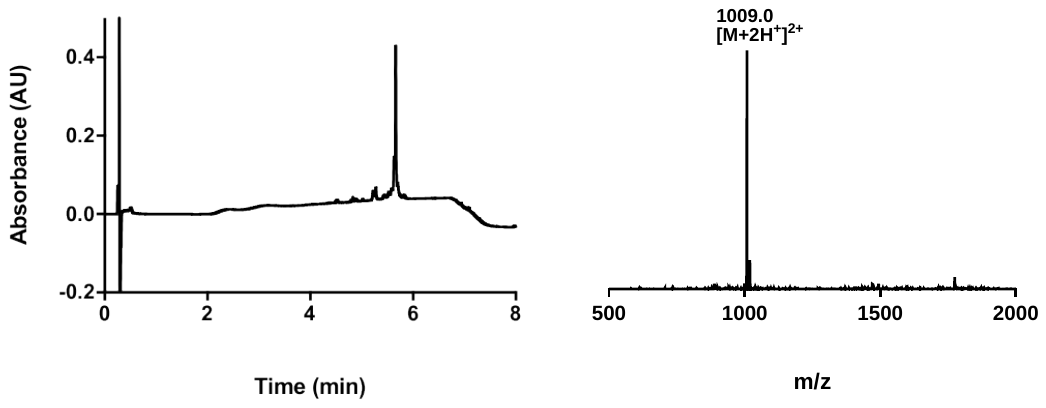

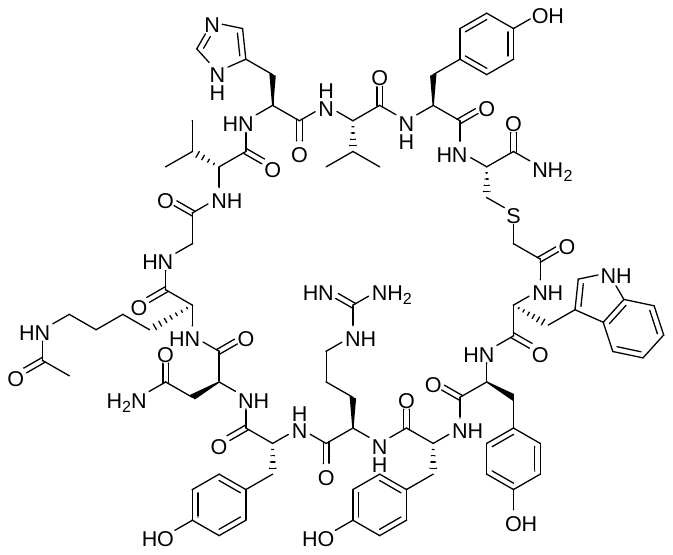

**2.1E** was synthesised via Fmoc-strategy SPPS as specified in general methods. The linear sequence *N’*- WYYRYNK(Ac)GVHVY -*C’* was generated by automated SPPS on Rink amide resin (86 mg, 50µmol, capacity: 0.58 mmolg^-1^). Coupling of chloroacetic acid to the N-terminus, cleavage from resin and cyclisation were performed as described in the general methods. The crude cyclic peptide was purified by semi-preparative RP-HPLC (0 to 20 % B + 0.1 % formic acid over 40 min). The appropriate fractions were combined and lyophilised to afford **2.1E** as a white solid. UPLC: *R_t_* = 4.33 min. (0 to 60 vol.% B over 5 min, 0.1 vol.% TFA, λ = 214 nm). LRMS (ESI+): *m/z =* 1832.5 [M + H]^+^.

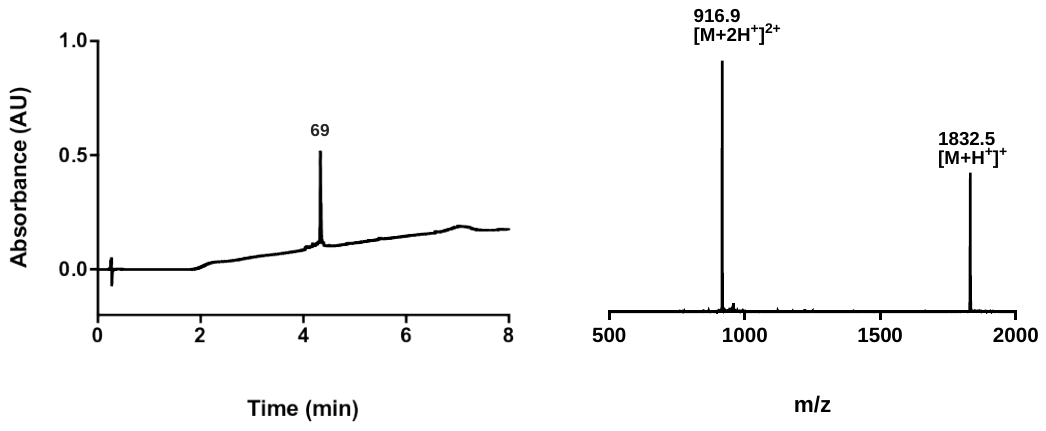

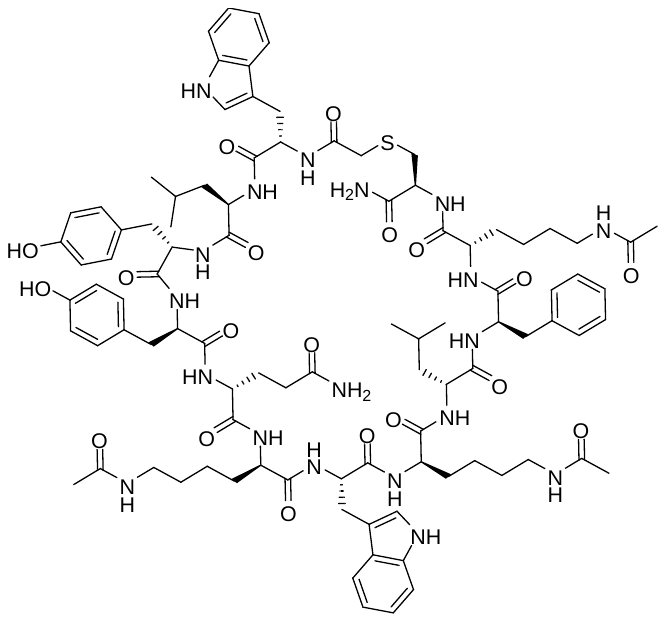

**2.1F** was synthesised via Fmoc-strategy SPPS as specified in general methods. The linear sequence *N’*- WLYYQK(Ac)WK(Ac)LFK(Ac) -*C’* was generated by automated SPPS on Rink amide resin (86 mg, 50µmol, capacity: 0.58 mmolg^-1^). Coupling of chloroacetic acid to the N-terminus, cleavage from resin and cyclisation were performed as described in the general methods. The crude cyclic peptide was purified by semi-preparative RP-HPLC (0 to 20 % B + 0.1 % formic acid over 40 min). The appropriate fractions were combined and lyophilised to afford **2.1F** as a white solid. UPLC: *R_t_* = 7.07 min. (0 to 60 vol.% B over 5 min, 0.1 vol.% TFA, λ = 214 nm). LRMS (ESI+): *m/z =* 1907.9 [M + H]^+^.

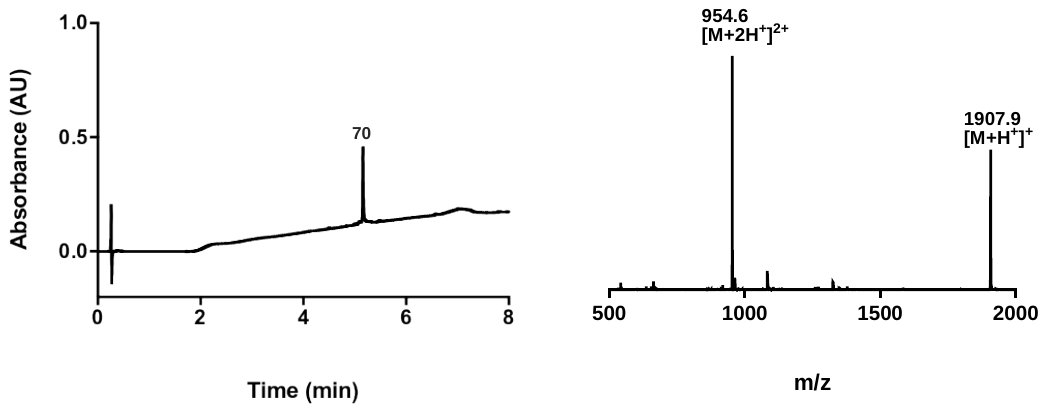

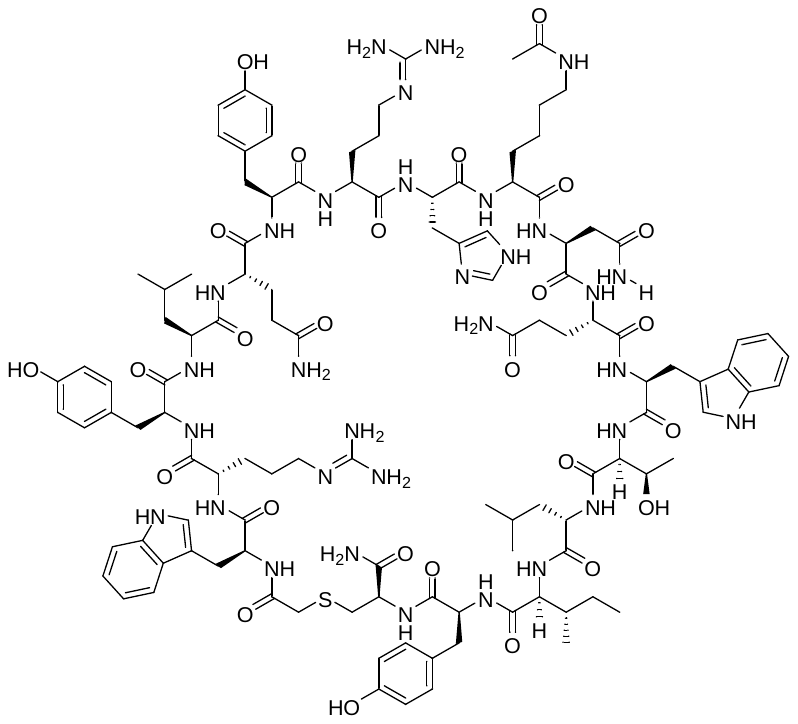

**4.1A** was synthesised via Fmoc-strategy SPPS as specified in general methods. The linear sequence *N’*- WRYLQYRHK(Ac)NQWTLIYC -*C’* was generated by automated SPPS on Rink amide resin (86 mg, 50µmol, capacity: 0.58 mmolg^-1^). Coupling of chloroacetic acid to the N-terminus, cleavage from resin and cyclisation were performed as described in the general methods. The crude cyclic peptide was purified by semi-preparative RP-HPLC (0 to 20 % B + 0.1 % formic acid over 40 min). The appropriate fractions were combined and lyophilised to afford **4.1A** as a white solid. **UPLC:** *R_t_* = 7.61 min. (0 to 60 vol.% B over 5 min, 0.1 vol.% TFA, λ = 214 nm). **LRMS** (ESI+): *m/z* = 1227.1 [M + 2H]^+^.

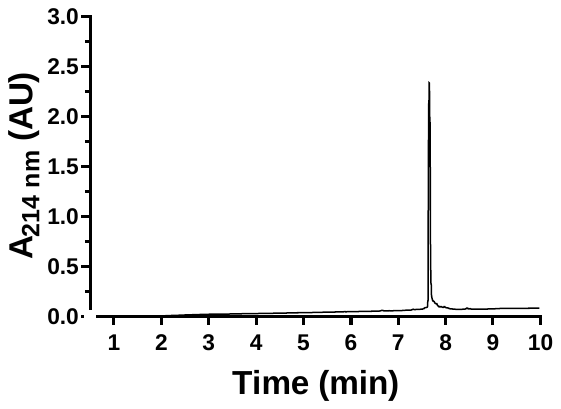

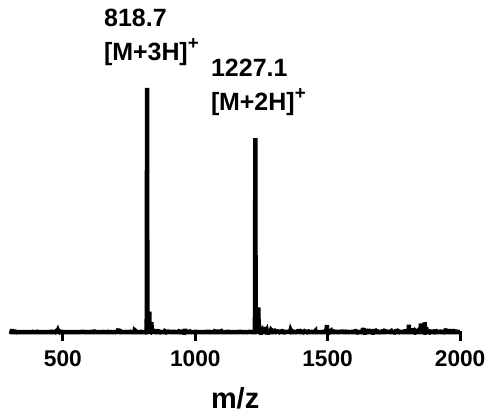

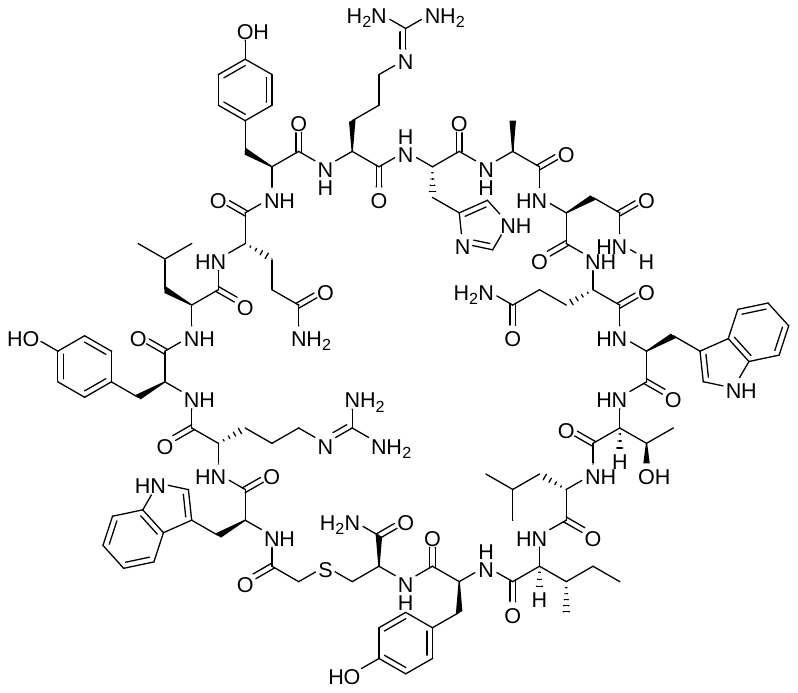

**4.1A Kac9A** was synthesised via Fmoc-strategy SPPS as specified in general methods. The linear sequence *N’*- WRYLQYRHKNQWTLIYC -*C’* was generated by automated SPPS on Rink amide resin (86 mg, 50µmol, capacity: 0.58 mmolg^-1^). Coupling of chloroacetic acid to the N-terminus, cleavage from resin and cyclisation were performed as described in the general methods. The crude cyclic peptide was purified by semi-preparative RP-HPLC (0 to 20 % B + 0.1 % formic acid over 40 min). The appropriate fractions were combined and lyophilised to afford **4.1A_Kac9A** as a white solid. **UPLC:** *R_t_* = 6.09 min. (0 to 60 vol.% B over 5 min, 0.1 vol.% TFA, λ = 214 nm). **LRMS** (ESI+): *m/z =* 1177.6 [M + 2H]^+^.

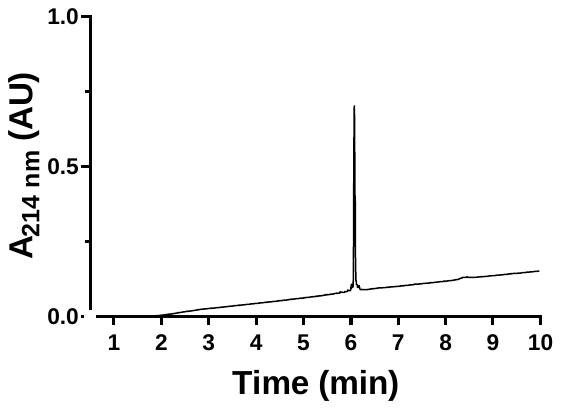

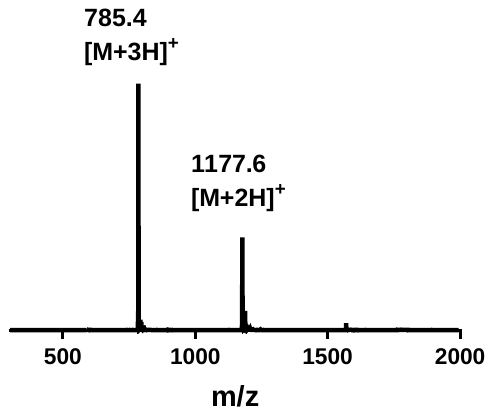

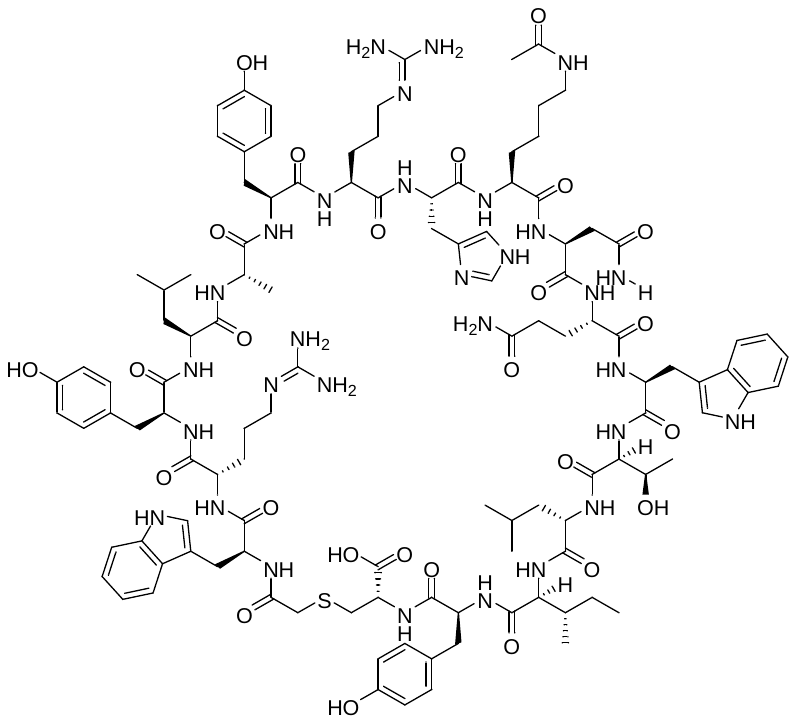

**4.1A Q5A** was synthesised via Fmoc-strategy SPPS as specified in general methods. The linear sequence *N’*- WRYLAYRHK(Ac)NQWTLIYC -*C’* was generated by automated SPPS on Rink amide resin (86 mg, 50µmol, capacity: 0.58 mmolg^-1^). Coupling of chloroacetic acid to the N-terminus, cleavage from resin and cyclisation were performed as described in the general methods. The crude cyclic peptide was purified by semi-preparative RP-HPLC (0 to 20 % B + 0.1 % formic acid over 40 min). The appropriate fractions were combined and lyophilised to afford **4.1A_Q5A** as a white solid **UPLC:** *R_t_* = 6.22 min. (0 to 60 vol.% B over 5 min, 0.1 vol.% TFA, λ = 214 nm). **LRMS** (ESI+): *m/z =* 1198.8 [M + 2H]^+^.

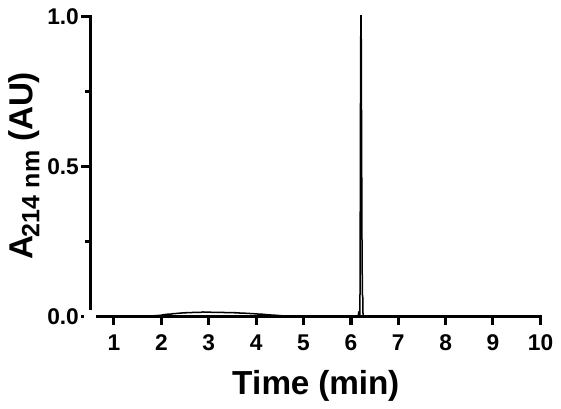

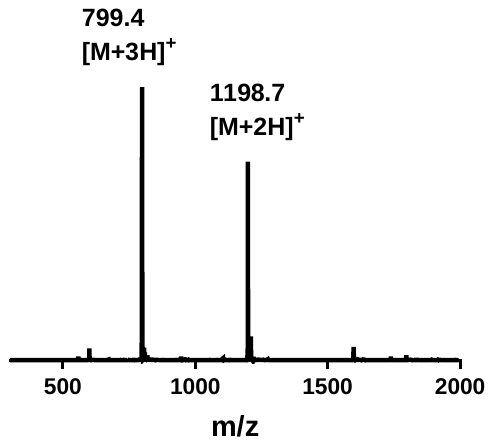

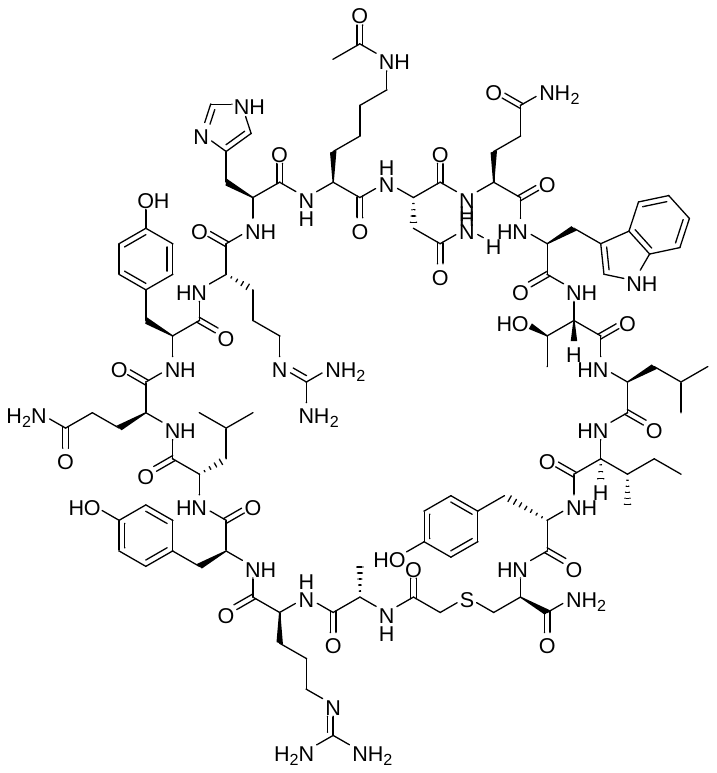

**4.1A W1A** was synthesised via Fmoc-strategy SPPS as specified in general methods. The linear sequence *N’*-ARYLQYRHKNQWTLIYC -*C’* was generated by automated SPPS on Rink amide resin (86 mg, 50µmol, capacity: 0.58 mmolg^-1^). Coupling of chloroacetic acid to the N-terminus, cleavage from resin and cyclisation were performed as described in the general methods. The crude cyclic peptide was purified by semi-preparative RP-HPLC (0 to 20 % B + 0.1 % formic acid over 40 min). The appropriate fractions were combined and lyophilised to afford **4.1A_W1A** as a white solid **UPLC:** *R_t_* = 5.65 min. (0 to 60 vol.% B over 5 min, 0.1 vol.% TFA, λ = 214 nm). **LRMS** (ESI+): *m/z =* 1169.8 [M + 2H]^+^.

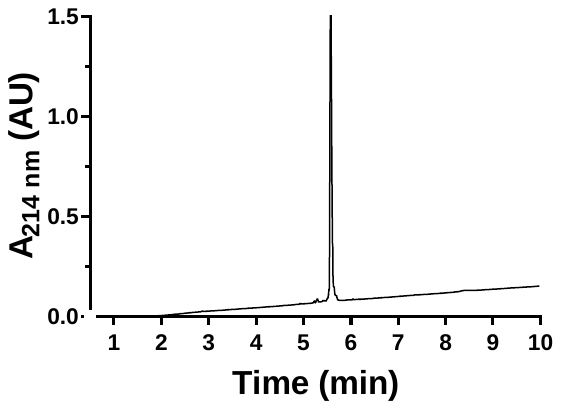

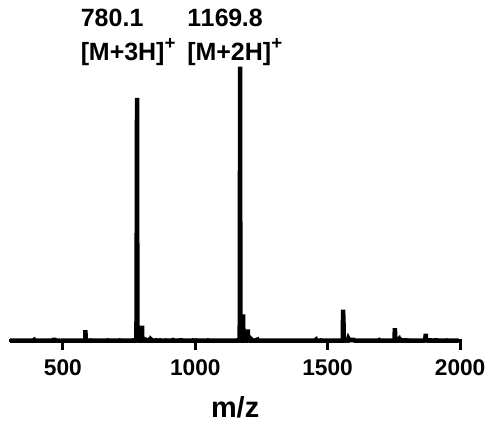

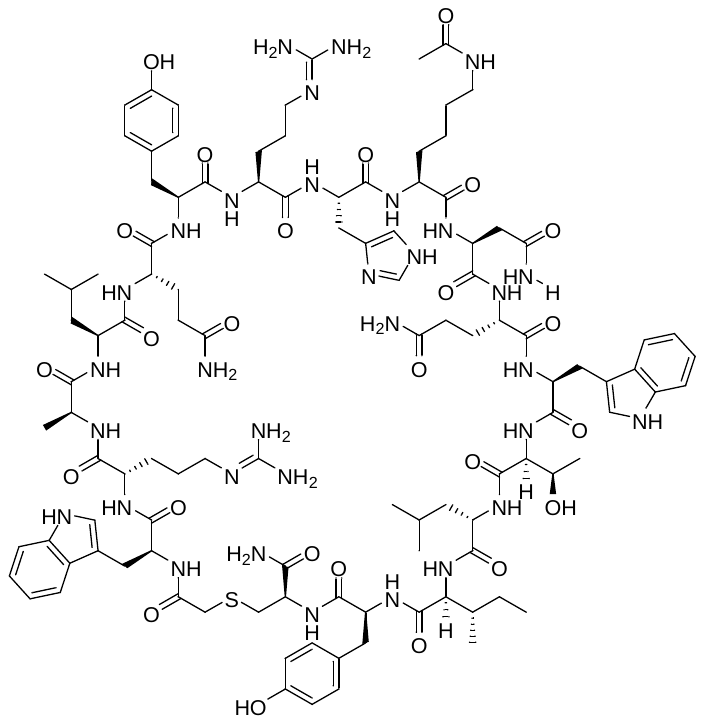

**4.1A Y3A** was synthesised via Fmoc-strategy SPPS as specified in general methods. The linear sequence *N’*-WRALQYRHK(Ac)NQWTLIYC -*C’* was generated by automated SPPS on Rink amide resin (86 mg, 50µmol, capacity: 0.58 mmolg^-1^). Coupling of chloroacetic acid to the N-terminus, cleavage from resin and cyclisation were performed as described in the general methods. The crude cyclic peptide was purified by semi-preparative RP-HPLC (0 to 20 % B + 0.1 % formic acid over 40 min). The appropriate fractions were combined and lyophilised to afford **4.1A_Y3A** as a white solid **UPLC:** *R_t_* = 6.01min. (0 to 60 vol.% B over 5 min, 0.1 vol.% TFA, λ = 214 nm). **LRMS** (ESI+): *m/z =* 1181.1 [M + 2H]^+^.

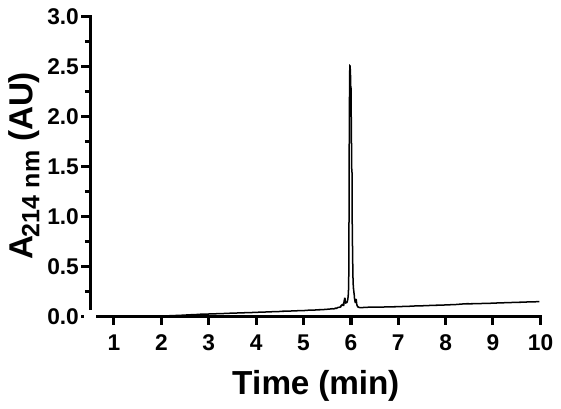

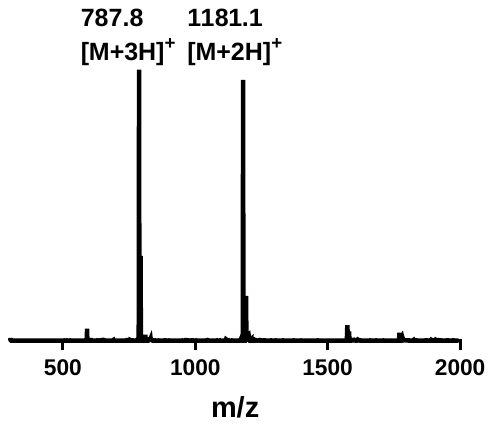

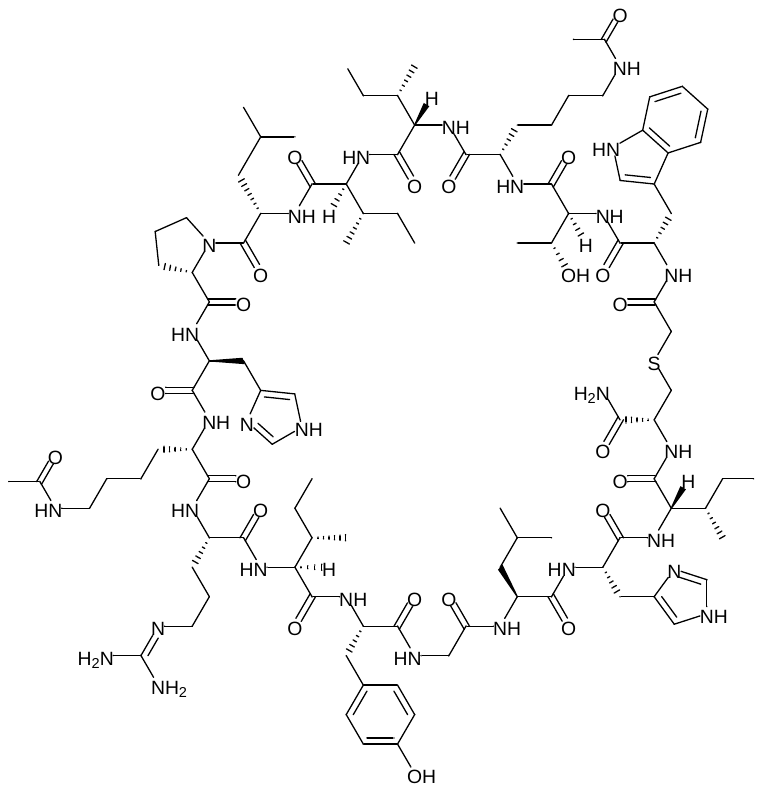

**4.1B** was synthesised via Fmoc-strategy SPPS as specified in general methods. The linear sequence *N’*-WTK(Ac)IILPHK(Ac)RIYGLHIC -*C’* was generated by automated SPPS on Rink amide resin (86 mg, 50µmol, capacity: 0.58 mmolg^-1^). Coupling of chloroacetic acid to the N-terminus, cleavage from resin and cyclisation were performed as described in the general methods. The crude cyclic peptide was purified by semi-preparative RP-HPLC (0 to 20 % B + 0.1 % formic acid over 40 min). The appropriate fractions were combined and lyophilised to afford **4.1B** as a white solid **UPLC:** *R_t_* = 6.31 min. (0 to 60 vol.% B over 5 min, 0.1 vol.% TFA, λ = 214 nm). **LRMS** (ESI+): *m/z =* 1108.2 [M + 2H]^+^.

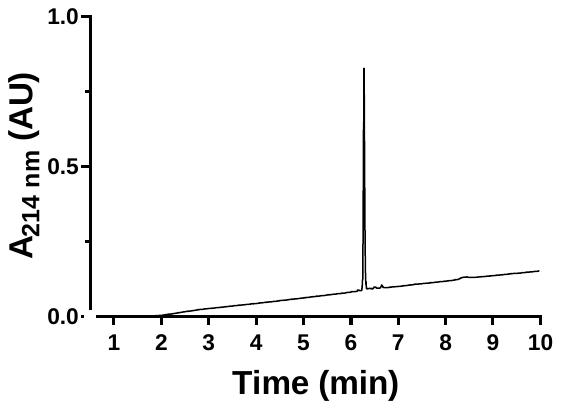

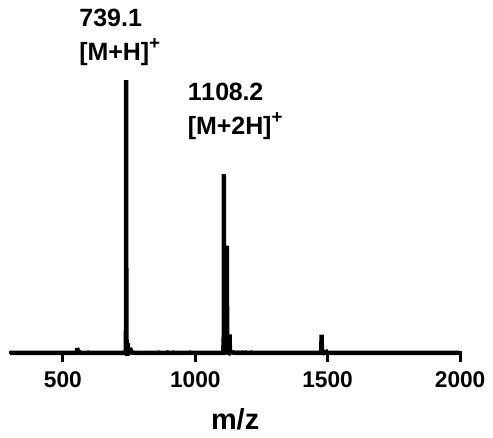

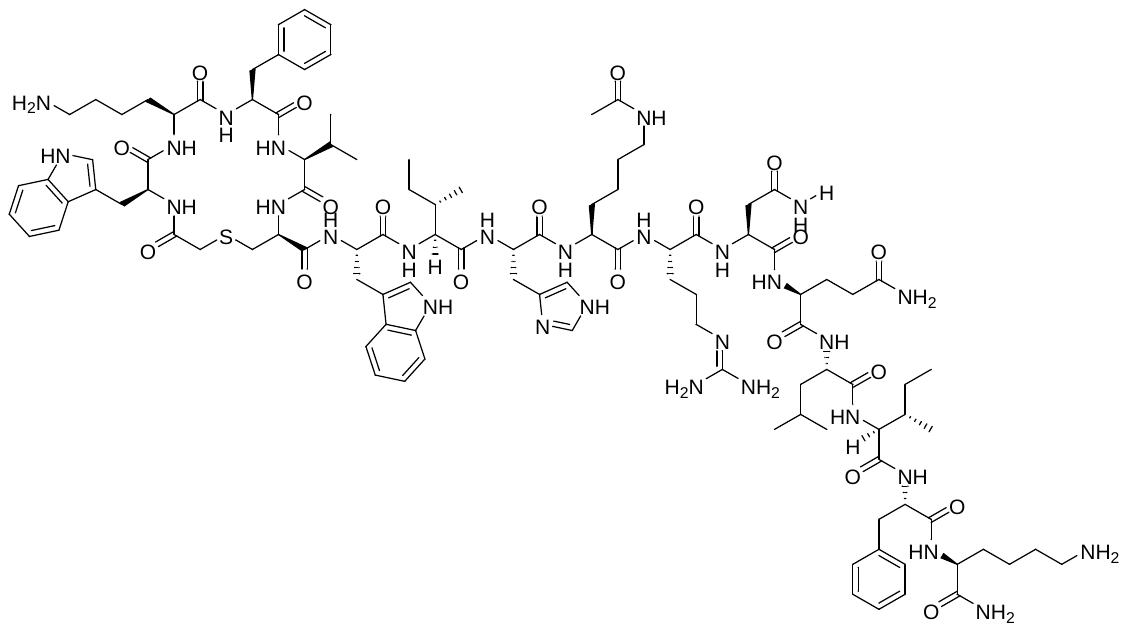

**4.1C** was synthesised via Fmoc-strategy SPPS as specified in general methods. The linear sequence *N’*-WKFVCWIHK(Ac)RNQLIFK -*C’* was generated by automated SPPS on Rink amide resin (86 mg, 50µmol, capacity: 0.58 mmolg^-1^). Coupling of chloroacetic acid to the N-terminus, cleavage from resin and cyclisation were performed as described in the general methods. The crude cyclic peptide was purified by semi-preparative RP-HPLC (0 to 20 % B + 0.1 % formic acid over 40 min). The appropriate fractions were combined and lyophilised to afford **4.1C** as a white solid **UPLC:** *R_t_* = 6.51 min. (0 to 60 vol.% B over 5 min, 0.1 vol.% TFA, λ = 214 nm). **LRMS** (ESI+): *m/z =* 1114.5 [M + 2H]^+^.

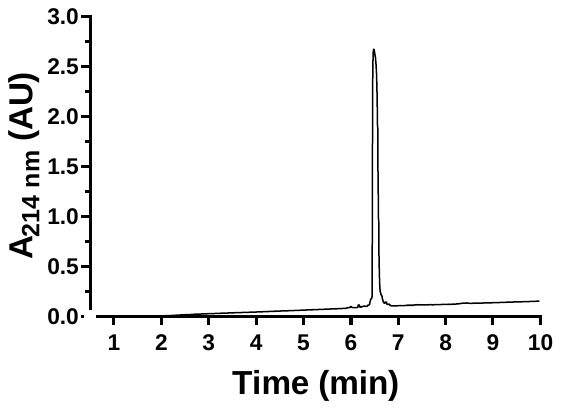

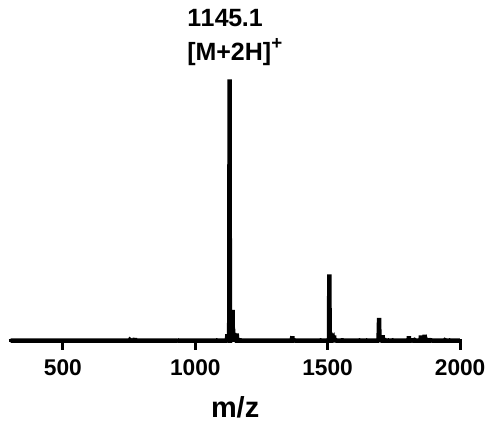

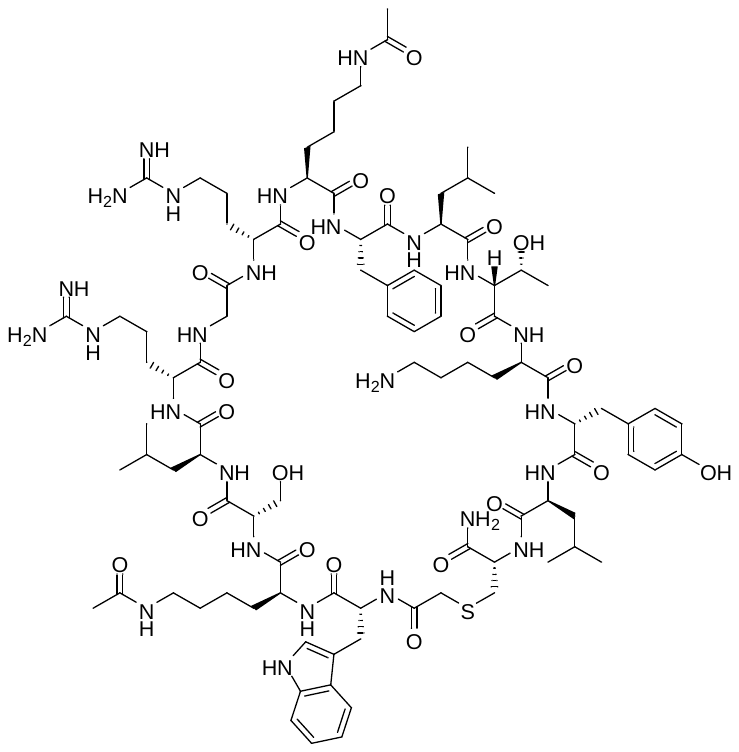

**4.1D** was synthesised via Fmoc-strategy SPPS as specified in general methods. The linear sequence *N’*- WK(Ac)SLRGRK(Ac)FLTKYLC -*C’* was generated by automated SPPS on Rink amide resin (86 mg, 50µmol, capacity: 0.58 mmolg^-1^). Coupling of chloroacetic acid to the N-terminus, cleavage from resin and cyclisation were performed as described in the general methods. The crude cyclic peptide was purified by semi-preparative RP-HPLC (0 to 20 % B + 0.1 % formic acid over 40 min). The appropriate fractions were combined and lyophilised to afford **4.1D** as a white solid. UPLC: *R_t_* = 6.54 min. (0 to 60 vol.% B over 5 min, 0.1 vol.% TFA, λ = 214 nm). LRMS (ESI+): *m/z =* 1907.9 [M + H]^+^.

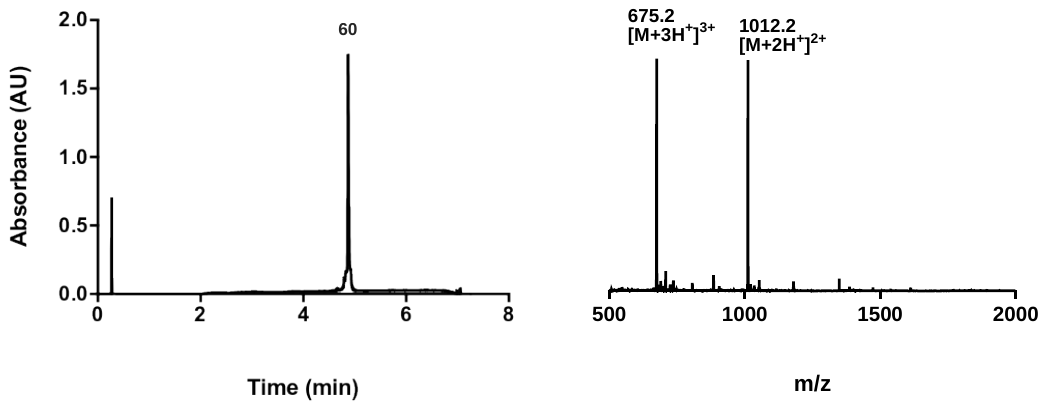

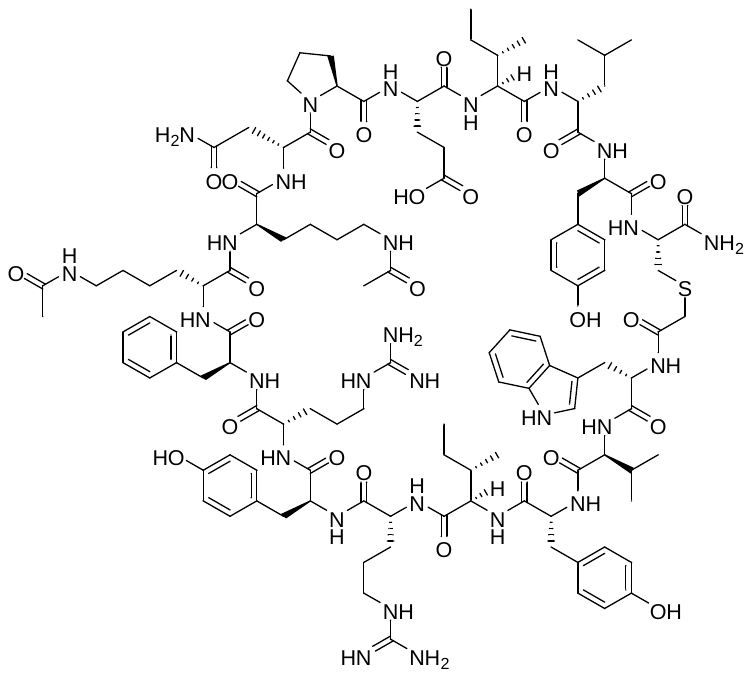

**4.1E** was synthesised via Fmoc-strategy SPPS as specified in general methods. The linear sequence *N’*- WVYIRYRFK(Ac)K(Ac)NPEILYC -*C’* was generated by automated SPPS on Rink amide resin (86 mg, 50µmol, capacity: 0.58 mmolg^-1^). Coupling of chloroacetic acid to the N-terminus, cleavage from resin and cyclisation were performed as described in the general methods. The crude cyclic peptide was purified by semi-preparative RP-HPLC (0 to 20 % B + 0.1 % formic acid over 40 min). The appropriate fractions were combined and lyophilised to afford **4.1E** as a white solid. UPLC: *R_t_* = 4.98 min. (0 to 60 vol.% B over 5 min, 0.1 vol.% TFA, λ = 214 nm). LRMS (ESI+): *m/z =* 1208.2 [M + 2H]^2+^.

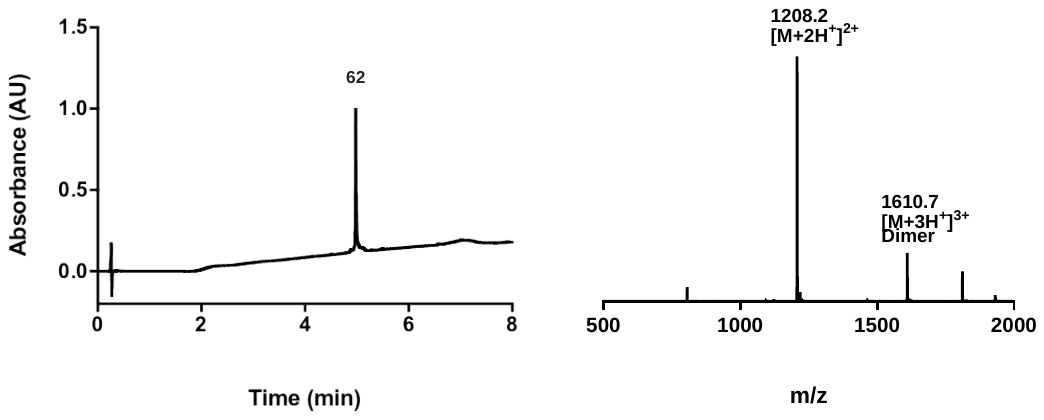

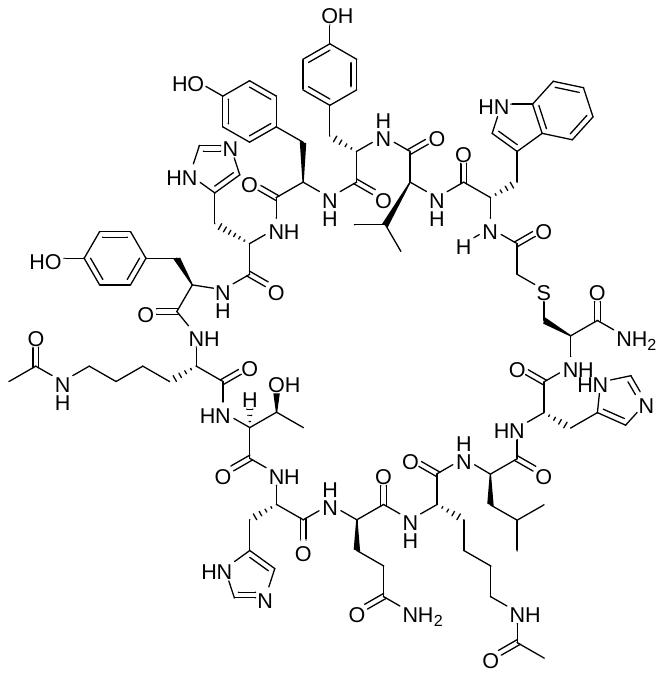

**4.1F** was synthesised via Fmoc-strategy SPPS as specified in general methods. The linear sequence *N’*- WVYYHYK(Ac)THQK(Ac)LHC -*C’* was generated by automated SPPS on Rink amide resin (86 mg, 50µmol, capacity: 0.58 mmolg^-1^). Coupling of chloroacetic acid to the N-terminus, cleavage from resin and cyclisation were performed as described in the general methods. The crude cyclic peptide was purified by semi-preparative RP-HPLC (0 to 20 % B + 0.1 % formic acid over 40 min). The appropriate fractions were combined and lyophilised to afford **4.1F** as a white solid. UPLC: *R_t_* = 4.24 min. (0 to 60 vol.% B over 5 min, 0.1 vol.% TFA, λ = 214 nm). LRMS (ESI+): *m/z =* 1015.5 [M + 2H]^2+^.

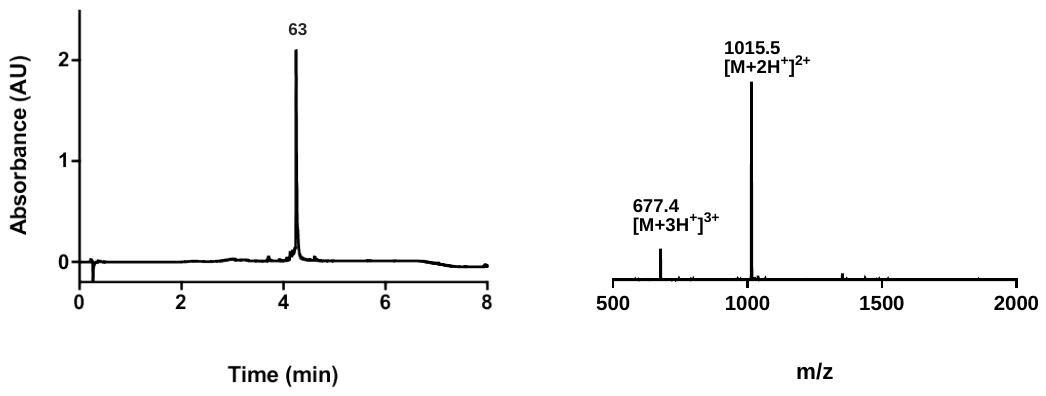

**TABLES**

**Table S3.** Peptide sequences and equilibrium dissociation constants (*K*ᴅ) measured by SPR for the binding of the cyclic peptides (CPs) selected for study from the RaPID selections against BRD2-BD1 and BRD4-BD1.

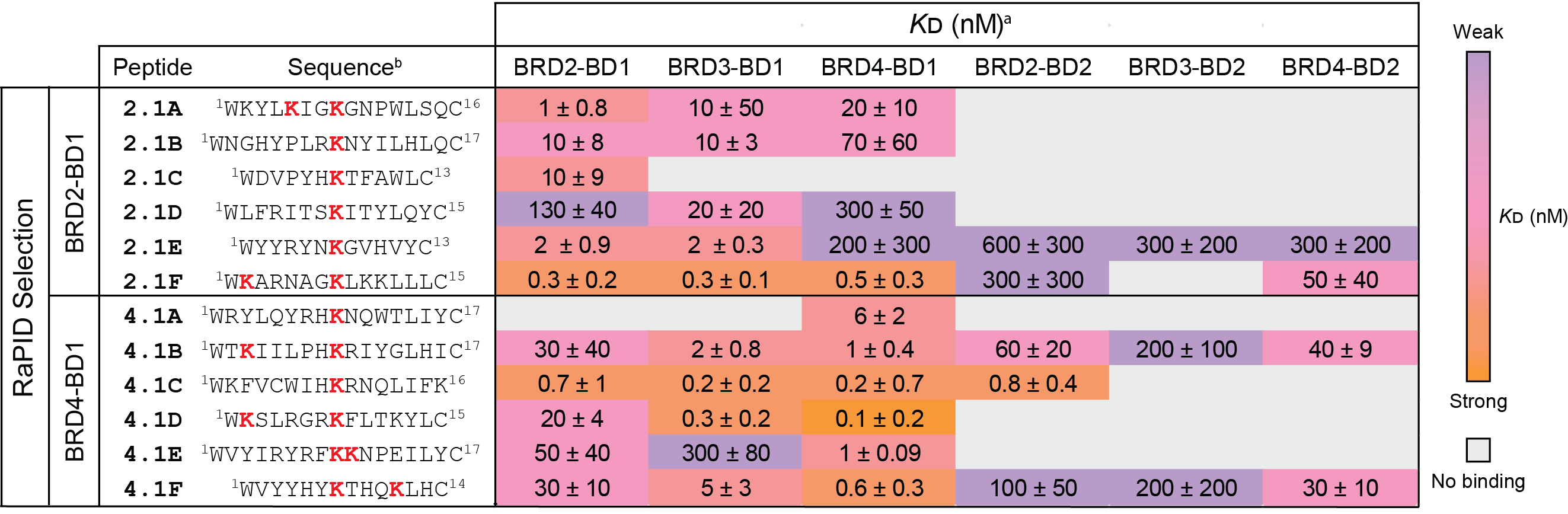

^a^ The *K*ᴅ values are given as the geometric means (± standard deviation) of at least three independent measurements.

^b^ Acetyllysine (AcK) residues are coloured in *red*.

^c^ The interactions are coloured according to binding strength using a *purple* (weak) to *orange* (strong) colour gradient.

^d^ No binding was observed for the combinations coloured in *grey*.

**Table S4.** Data collection and refinement statistics for the crystal structures of BRD4-BD1 in complex with 4.1A (PDB ID: 9MPI) and BRD2-BD1 in complex with 2.1C (9MPJ) solved using molecular replacement.

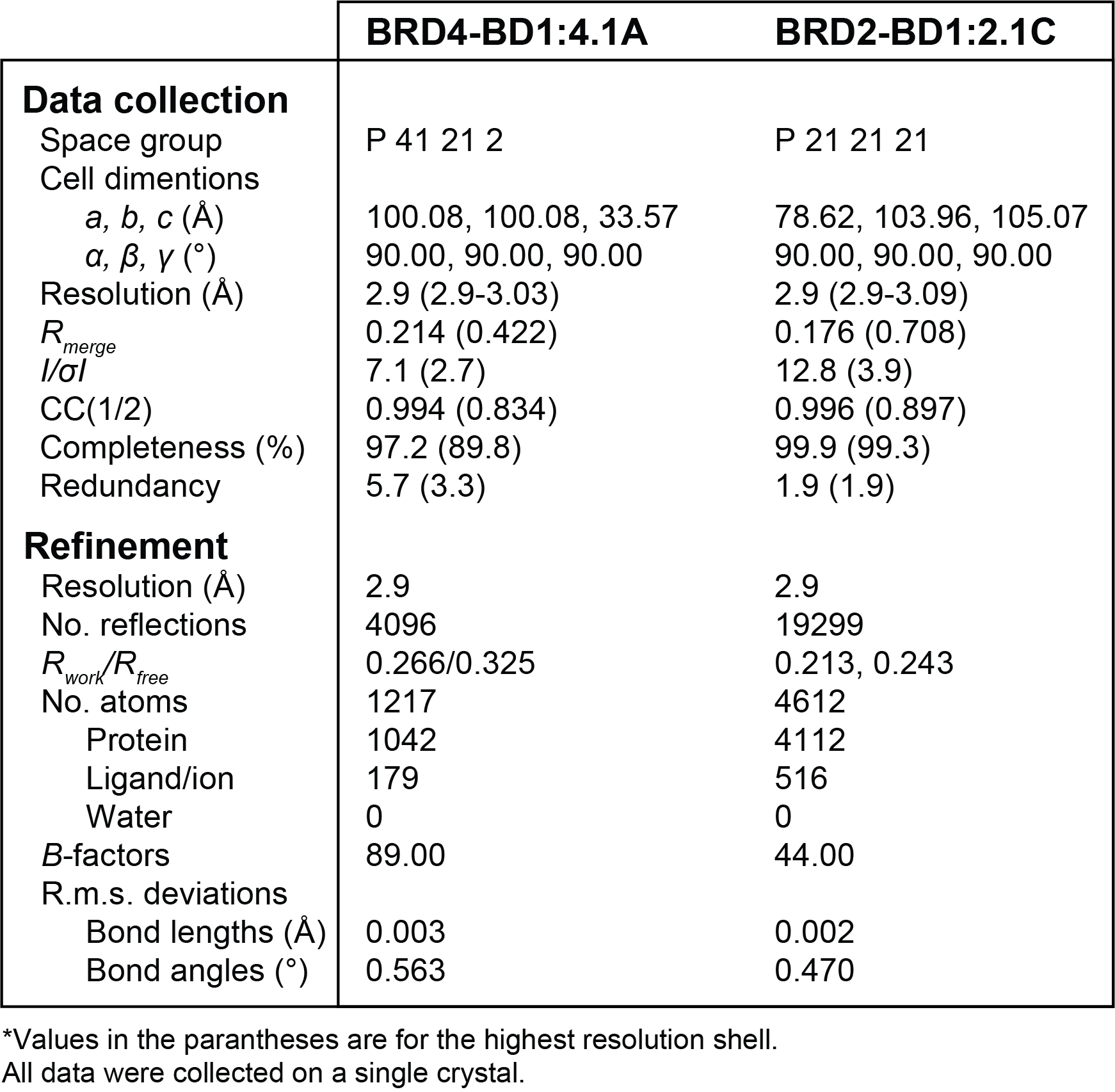

**Table S5.** Data collection and refinement statistics for the crystal structure of BRD2-BD1 in complex with 2.1C W11A (PDB ID: 9MPL), BRD3-BD1 in complex with 2.1C W11A (PDB ID: 9MPM), and BRD3-BD1 A128T mutant (PDB ID: 9MPN) solved using molecular replacement.

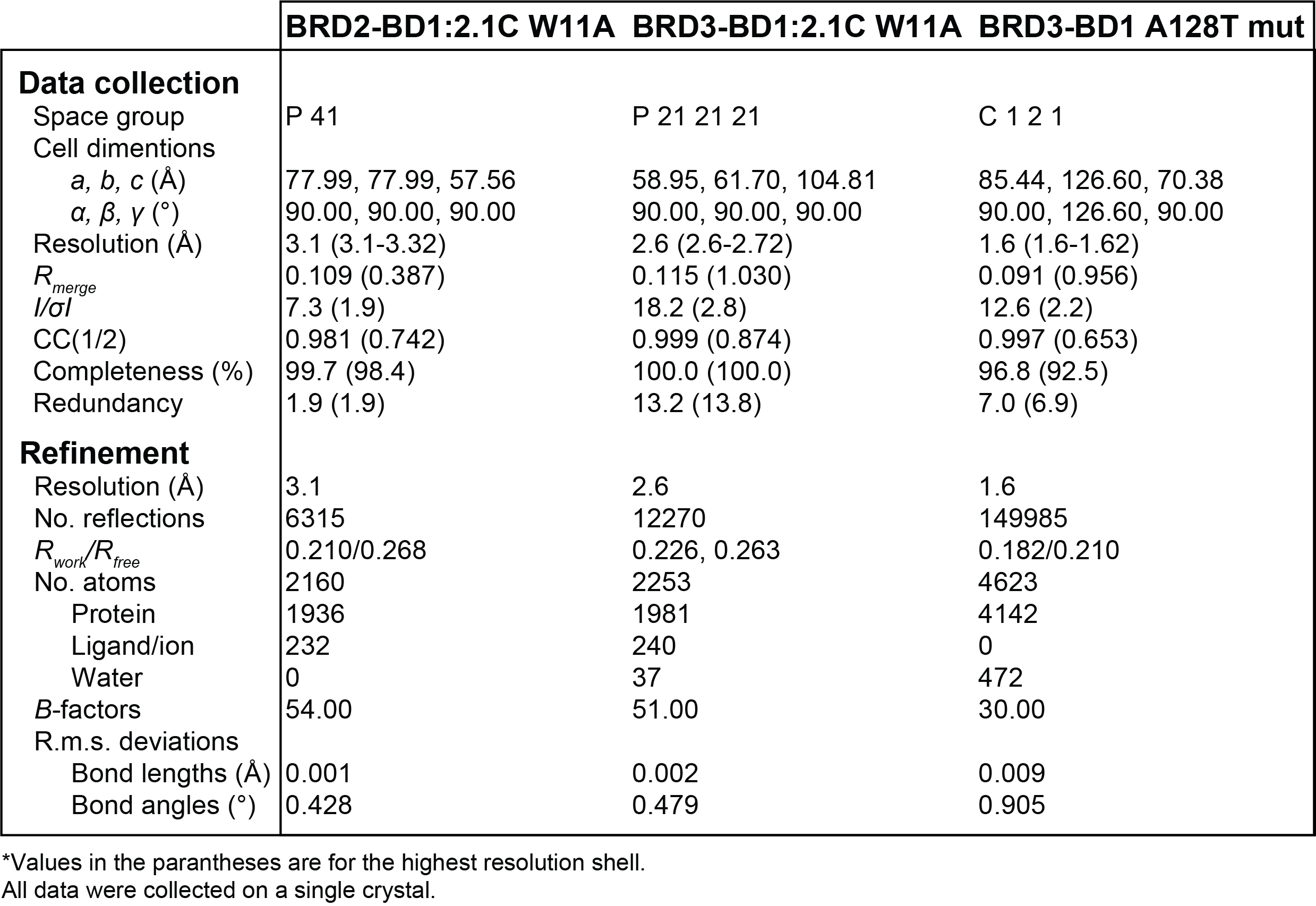

**Table S6.** Random coil deviation for Cα and Cβ shifts of BRD3-BD1.

| Amino acid | | Cα [ppm] | Cβ [ppm] | (Cα-Cβ)[ppm] | Secondary Structure |
| --- | --- | --- | --- | --- | --- |
| GLY | Expression tag residuals | 0 | 0 | 0 |  |
| PRO |  | -1.23 | -0.68 | -0.241 |  |
| LEU |  | -1.205 | -1.032 | -0.607 |  |
| GLY |  | -1.098 | 0 | -0.557 |  |
| SER |  | -1.138 | -0.739 | -0.665 |  |
| GLU- | 25 | -1.568 | -1.07 | -0.489 |  |
| VAL | 26 | -1.82 | -1.251 | -0.875 |  |
| SER | 27 | -1.903 | -0.344 | -0.709 |  |
| ASN | 28 | 0 | 0 | 0 |  |
| PRO | 29 | -0.664 | -0.717 | 0.036 |  |
| SER | 30 | -0.92 | -0.974 | -0.623 |  |
| LYS+ | 31 | -2.275 | -0.298 | -0.747 |  |
| PRO | 32 | -0.756 | -0.439 | -1.488 |  |
| GLY | 33 | -2.171 | 0 | -2.402 | STRAND |
| ARG+ | 34 | -2.698 | 2.021 | -2.985 | STRAND |
| LYS+ | 35 | -2.043 | 0.021 | -4.895 | STRAND |
| THR | 36 | -5.142 | 2.76 | -1.254 | STRAND |
| ASN | 37 | 2.176 | -4.027 | 0.094 |  |
| GLN | 38 | 1.857 | -0.125 | 3.978 | HELIX |
| LEU | 39 | 2.356 | -1.392 | 2.84 | HELIX |
| GLN | 40 | 1.858 | -0.933 | 3.735 | HELIX |
| TYR | 41 | 2.77 | -1.895 | 3.614 | HELIX |
| MET | 42 | 0.395 | -2.991 | 3.522 | HELIX |
| GLN | 43 | 2.192 | -0.323 | 2.469 | HELIX |
| ASN | 44 | 0.47 | -1.037 | 2.255 | HELIX |
| VAL | 45 | 1.154 | -1.589 | 4.324 | HELIX |
| VAL | 46 | 5.772 | -2.951 | 6.241 | HELIX |
| VAL | 47 | 4.451 | -2.806 | 6.25 | HELIX |
| LYS+ | 48 | 1.943 | -0.826 | 5.159 | HELIX |
| THR | 49 | 3.763 | -1.688 | 3.725 | HELIX |
| LEU | 50 | 2.002 | -0.954 | 4.292 | HELIX |
| TRP | 51 | 2.181 | -2.289 | 2.028 | HELIX |
| LYS+ | 52 | -2.064 | -0.722 | 1.315 | HELIX |
| HIS | 53 | 1.882 | 1.065 | 1.118 | HELIX |
| GLN | 54 | 1.491 | -2.389 | 0.665 |  |
| PHE | 55 | -5.991 | -3.288 | 0.127 |  |
| ALA | 56 | 0.099 | 0.896 | 0.778 |  |
| TRP | 57 | 2.194 | -3.639 | 2.562 | HELIX |
| PRO | 58 | 0.279 | -2.371 | 2.778 |  |
| PHE | 59 | -2.272 | -2.123 | 0.924 |  |
| TYR | 60 | -2.297 | -2.569 | -0.682 |  |
| GLN | 61 | -2.702 | -0.534 | -0.905 |  |
| PRO | 62 | -1.423 | -0.604 | -0.138 |  |
| VAL | 63 | 1.158 | -1.415 | -0.104 |  |
| ASP- | 64 | -3.623 | -1.557 | 0.58 |  |
| ALA | 65 | 0.509 | -0.724 | 0.876 |  |
| ILE | 66 | 0.627 | -2.834 | 2.282 | HELIX |
| LYS+ | 67 | 1.041 | -1.111 | 1.723 | HELIX |
| LEU | 68 | -1.78 | -1.335 | 1.354 | HELIX |
| ASN | 69 | -0.584 | -2.939 | -0.068 |  |
| LEU | 70 | -3.459 | -1.345 | 0.08 |  |
| PRO | 71 | 0 | 0 | 0 |  |
| ASP- | 72 | -1.628 | -1.606 | 1.894 | HELIX |
| TYR | 73 | 4.811 | -0.893 | 2.743 | HELIX |
| HIS | 74 | 0.524 | -2.023 | 3.019 | HELIX |
| LYS+ | 75 | -0.319 | -1.124 | 0.975 | HELIX |
| ILE | 76 | -0.258 | 0.168 | 0.259 |  |
| ILE | 77 | -5.364 | -5.763 | -0.122 |  |
| LYS+ | 78 | -1.264 | -0.926 | -0.67 |  |
| ASN | 79 | -2.169 | -0.099 | -0.87 |  |
| PRO | 80 | -1.064 | -0.862 | -1.077 |  |
| MET | 81 | -3.111 | -2.153 | -2.073 | STRAND |
| ASP- | 82 | -2.628 | 2.43 | -1.611 | STRAND |
| MET | 83 | 2.339 | 1.155 | -0.748 |  |
| GLY | 84 | 1.629 | 0 | 2.434 | HELIX |
| THR | 85 | 4.49 | 0 | 3.403 | HELIX |
| ILE | 86 | 3.38 | -0.71 | 4.073 | HELIX |
| LYS+ | 87 | 3.092 | -0.547 | 3.504 | HELIX |
| LYS+ | 88 | 1.225 | -1.558 | 3.575 | HELIX |
| ARG+ | 89 | 2.543 | -1.761 | 3.395 | HELIX |
| LEU | 90 | 1.713 | -1.385 | 3.135 | HELIX |
| GLU- | 91 | 0.583 | -1.42 | 1.454 | HELIX |
| ASN | 92 | -1.833 | -1.093 | 1.547 | HELIX |
| ASN | 93 | 0.059 | -3.318 | 1.106 | HELIX |
| TYR | 94 | 1.268 | 0.587 | 1.216 | HELIX |
| TYR | 95 | -1.727 | -1.316 | 0.32 |  |
| TRP | 96 | 0.273 | -0.418 | -1.876 | STRAND |
| SER | 97 | -4.861 | 1.048 | -0.838 |  |
| ALA | 98 | 1.189 | -1.516 | -0.338 |  |
| SER | 99 | 2.19 | 0 | 1.408 | HELIX |
| GLU- | 100 | 0.26 | 0.931 | 2.011 | HELIX |
| CYS | 101 | 2.796 | -1.717 | 2.747 | HELIX |
| MET | 102 | 3.009 | -1.39 | 4.183 | HELIX |
| GLN | 103 | 1.545 | -2.092 | 4.34 | HELIX |
| ASP- | 104 | 2.759 | -2.224 | 4.787 | HELIX |
| PHE | 105 | 4.514 | -1.228 | 4.847 | HELIX |
| ASN | 106 | 2.826 | -0.989 | 4.944 | HELIX |
| THR | 107 | 3.462 | -1.813 | 3.957 | HELIX |
| MET | 108 | 3.045 | 0.265 | 4.43 | HELIX |
| PHE | 109 | 2.276 | -2.959 | 4.518 | HELIX |
| THR | 110 | 3.897 | -1.641 | 4.391 | HELIX |
| ASN | 111 | 0.725 | -1.674 | 4.453 | HELIX |
| CYS | 112 | 3.09 | -2.333 | 3.152 | HELIX |
| TYR | 113 | -2.432 | -4.066 | 3.063 | HELIX |
| ILE | 114 | 1.18 | -0.952 | 2.168 | HELIX |
| TYR | 115 | 2.861 | 0.122 | 0.669 |  |
| ASN | 116 | -1 | 1.863 | 0.245 |  |
| LYS+ | 117 | -1.179 | -2.037 | -0.162 |  |
| PRO | 118 | 0.817 | -0.701 | 0.367 |  |
| THR | 119 | -2.848 | -1.573 | 0.05 |  |
| ASP- | 120 | -0.553 | -0.461 | 0.86 |  |
| ASP- | 121 | 2.245 | -1.702 | 2.111 | HELIX |
| ILE | 122 | 0.4 | -2.078 | 4.172 | HELIX |
| VAL | 123 | 3.62 | -2.47 | 4.225 | HELIX |
| LEU | 124 | 1.54 | -2.567 | 4.425 | HELIX |
| MET | 125 | 2.841 | -0.238 | 3.188 | HELIX |
| ALA | 126 | 1.828 | -0.551 | 2.842 | HELIX |
| GLN | 127 | 1.315 | -1.754 | 2.668 | HELIX |
| ALA | 128 | 0.846 | -1.709 | 2.449 | HELIX |
| LEU | 129 | 0.902 | -0.82 | 2.493 | HELIX |
| GLU- | 130 | 0.869 | -2.332 | 2.841 | HELIX |
| LYS+ | 131 | 2.21 | -1.39 | 3.489 | HELIX |
| ILE | 132 | 1.572 | -2.095 | 4.017 | HELIX |
| PHE | 133 | 2.779 | -2.005 | 3.76 | HELIX |
| LEU | 134 | 1.153 | -1.675 | 3.584 | HELIX |
| GLN | 135 | 1.355 | -1.786 | 2.147 | HELIX |
| LYS+ | 136 | -1.239 | -1.71 | 3.129 | HELIX |
| VAL | 137 | 3.26 | -2.516 | 2.8 | HELIX |
| ALA | 138 | 0.246 | -1.907 | 1.909 | HELIX |
| GLN | 139 | -3.13 | -0.927 | -0.834 |  |
| MET | 140 | -1.13 | 1.322 | -1.614 | STRAND |
| PRO | 141 | -1.01 | -0.824 | -0.482 |  |
| GLN | 142 | 0.476 | -0.716 | -0.824 |  |
| GLU- | 143 | -3.537 | -0.058 | -1.013 | STRAND |
| GLU- | 144 | -1.868 | -1.116 | -3.147 | STRAND |
| VAL | 145 | -3.475 | 1.736 | -2.181 | STRAND |
| GLU- | 146 | -1.227 | -0.646 | -2.239 | STRAND |
| LEU | 147 | -0.764 | 0.16 | -0.753 |  |

**Table S7.** Random coil deviation for Cα and Cβ shifts of BRD3-BD1 A128T.

| Amino Acid | | Cα [ppm] | Cβ [ppm] | (Cα-Cβ) [ppm] | Secondary Structure |
| --- | --- | --- | --- | --- | --- |
| GLY | Expression tag residuals | 0 | 0 | 0 |  |
| PRO |  | -0.288 | 0.441 | -0.325 |  |
| LEU |  | -0.288 | -0.042 | -0.363 |  |
| GLY |  | -0.114 | 0 | -0.32 |  |
| SER |  | -0.444 | 0.155 | -0.423 |  |
| GLU- | 25 | -0.608 | -0.051 | -0.613 |  |
| VAL | 26 | -0.903 | -0.219 | -0.932 |  |
| SER | 27 | -1.224 | 0.33 | -0.841 |  |
| ASN | 28 | -0.286 | 0 | -0.614 |  |
| PRO | 29 | 0.258 | 0.259 | -0.108 |  |
| SER | 30 | -0.107 | -0.071 | -0.441 |  |
| LYS+ | 31 | -1.287 | 0 | -0.56 |  |
| PRO | 32 | 0.204 | 0.561 | -0.946 |  |
| GLY | 33 | -1.193 | 0 | -2.132 | STRAND |
| ARG+ | 34 | -1.787 | 3.059 | -2.689 | STRAND |
| LYS+ | 35 | -1.165 | 0.863 | -4.929 | STRAND |
| THR | 36 | -4.237 | 3.675 | -1.265 | STRAND |
| ASN | 37 | 3.07 | -3.076 | 0.088 |  |
| GLN | 38 | 2.788 | 0.757 | 3.995 | HELIX |
| LEU | 39 | 3.265 | -0.543 | 2.873 | HELIX |
| GLN | 40 | 2.807 | 0.026 | 3.725 | HELIX |
| TYR | 41 | 3.668 | -0.918 | 3.571 | HELIX |
| MET | 42 | 1.298 | -2.047 | 3.484 | HELIX |
| GLN | 43 | 3.136 | 0.615 | 2.463 | HELIX |
| ASN | 44 | 1.343 | -0.179 | 2.159 | HELIX |
| VAL | 45 | 2.033 | -0.402 | 4.137 | HELIX |
| VAL | 46 | 6.632 | -1.823 | 5.98 | HELIX |
| VAL | 47 | 5.321 | -1.728 | 6.11 | HELIX |
| LYS+ | 48 | 2.909 | 0.084 | 5.145 | HELIX |
| THR | 49 | 4.758 | -0.803 | 3.74 | HELIX |
| LEU | 50 | 2.952 | 0.117 | 4.262 | HELIX |
| TRP | 51 | 3.122 | -1.268 | 1.959 | HELIX |
| LYS+ | 52 | -1.105 | 0.243 | 1.376 | HELIX |
| HIS | 53 | 2.969 | 1.884 | 0.635 |  |
| GLN | 54 | 2.169 | 0 | 0.261 |  |
| PHE | 55 | -5.191 | -2.72 | -0.366 |  |
| ALA | 56 | 1.048 | 1.844 | -0.061 |  |
| TRP | 57 | 3.085 | 0 | 1.623 | HELIX |
| PRO | 58 | 1.152 | -1.428 | 1.813 |  |
| PHE | 59 | -1.371 | -1.146 | 0.869 |  |
| TYR | 60 | -1.362 | -1.614 | -0.568 |  |
| GLN | 61 | -1.73 | 0 | -0.806 |  |
| PRO | 62 | -0.64 | 0.3 | -0.077 |  |
| VAL | 63 | 1.92 | -0.52 | -0.253 |  |
| ASP- | 64 | -2.806 | -0.547 | 0.436 |  |
| ALA | 65 | 1.402 | 0.275 | 0.768 |  |
| ILE | 66 | 1.503 | -1.933 | 2.184 | HELIX |
| LYS+ | 67 | 1.93 | -0.059 | 1.612 | HELIX |
| LEU | 68 | -0.936 | -0.348 | 1.212 | HELIX |
| ASN | 69 | 0.272 | -1.962 | -0.275 |  |
| LEU | 70 | -2.471 | 0 | 0.406 |  |
| PRO | 71 | 1.221 | -0.235 | -0.415 |  |
| ASP- | 72 | -0.782 | -0.551 | 2.296 | HELIX |
| TYR | 73 | 5.744 | 0.081 | 2.639 | HELIX |
| HIS | 74 | 1.412 | -1.072 | 2.944 | HELIX |
| LYS+ | 75 | 0.502 | -0.182 | 0.941 |  |
| ILE | 76 | 0.655 | 1.001 | 0.175 |  |
| ILE | 77 | -4.533 | -4.72 | -0.177 |  |
| LYS+ | 78 | -0.358 | 0.015 | -0.5 |  |
| ASN | 79 | -1.313 | 0 | -0.709 |  |
| PRO | 80 | -0.18 | 0.261 | -0.896 |  |
| MET | 81 | -2.153 | -1.218 | -2.153 | STRAND |
| ASP- | 82 | -1.711 | 3.372 | -1.646 | STRAND |
| MET | 83 | 3.26 | 2.18 | -0.479 |  |
| GLY | 84 | 2.565 | 0 | 2.842 | HELIX |
| THR | 85 | 4.88 | 0 | 3.815 | HELIX |
| ILE | 86 | 4.275 | 0.275 | 4.141 | HELIX |
| LYS+ | 87 | 4.007 | 0.464 | 3.417 | HELIX |
| LYS+ | 88 | 2.107 | -0.6 | 3.482 | HELIX |
| ARG+ | 89 | 3.473 | -0.724 | 3.319 | HELIX |
| LEU | 90 | 2.65 | -0.404 | 3.126 | HELIX |
| GLU- | 91 | 1.45 | -0.676 | 1.441 | HELIX |
| ASN | 92 | -0.903 | -0.046 | 1.513 | HELIX |
| ASN | 93 | 0.906 | -2.363 | 1.022 | HELIX |
| TYR | 94 | 2.126 | 1.472 | 1.103 | HELIX |
| TYR | 95 | -0.81 | -0.196 | 0.208 |  |
| TRP | 96 | 1.168 | 0.583 | -1.974 | STRAND |
| SER | 97 | -3.92 | 1.974 | -0.974 | STRAND |
| ALA | 98 | 2.107 | -0.28 | -0.116 |  |
| SER | 99 | 3.158 | 0 | 1.593 | HELIX |
| GLU- | 100 | 1.159 | 1.926 | 2.315 | HELIX |
| CYS | 101 | 3.786 | -0.769 | 2.725 | HELIX |
| MET | 102 | 3.999 | -0.388 | 4.162 | HELIX |
| GLN | 103 | 2.482 | -1.061 | 4.291 | HELIX |
| ASP- | 104 | 3.693 | -1.25 | 4.775 | HELIX |
| PHE | 105 | 5.471 | -0.368 | 4.892 | HELIX |
| ASN | 106 | 3.707 | -0.187 | 5.013 | HELIX |
| THR | 107 | 4.359 | -0.948 | 3.975 | HELIX |
| MET | 108 | 3.952 | 1.229 | 4.376 | HELIX |
| PHE | 109 | 3.094 | -2.003 | 4.415 | HELIX |
| THR | 110 | 4.754 | -0.67 | 4.281 | HELIX |
| ASN | 111 | 1.603 | -0.718 | 4.378 | HELIX |
| CYS | 112 | 4.031 | -1.358 | 3.11 | HELIX |
| TYR | 113 | -1.577 | -3.196 | 3.025 | HELIX |
| ILE | 114 | 2.084 | 0.017 | 2.142 | HELIX |
| TYR | 115 | 3.868 | 1.127 | 0.796 |  |
| ASN | 116 | 0.152 | 2.571 | 0.009 |  |
| LYS+ | 117 | -0.295 | 0 | -0.445 |  |
| PRO | 118 | 1.688 | 0.31 | -0.006 |  |
| THR | 119 | -1.844 | -0.744 | 0.13 |  |
| ASP- | 120 | 0.403 | 0.291 | 0.973 | HELIX |
| ASP- | 121 | 3.099 | -0.807 | 2.183 | HELIX |
| ILE | 122 | 1.362 | -1.169 | 4.23 | HELIX |
| VAL | 123 | 4.702 | -1.551 | 4.205 | HELIX |
| LEU | 124 | 2.307 | -1.523 | 4.606 | HELIX |
| MET | 125 | 4.139 | 0.404 | 3.319 | HELIX |
| ALA | 126 | 2.851 | 0.46 | 3.022 | HELIX |
| GLN | 127 | 2.193 | -0.748 | 3.61 | HELIX |
| THR | 128 | 4.895 | -0.603 | 3.555 | HELIX |
| LEU | 129 | 2.102 | -0.124 | 3.685 | HELIX |
| GLU- | 130 | 1.91 | -1.422 | 3.077 | HELIX |
| LYS+ | 131 | 3.164 | -0.51 | 3.521 | HELIX |
| ILE | 132 | 2.757 | -0.801 | 3.962 | HELIX |
| PHE | 133 | 3.488 | -1.167 | 3.69 | HELIX |
| LEU | 134 | 2.072 | -0.784 | 3.559 | HELIX |
| GLN | 135 | 2.318 | -0.848 | 2.154 | HELIX |
| LYS+ | 136 | -0.337 | -0.778 | 3.084 | HELIX |
| VAL | 137 | 4.201 | -1.443 | 2.743 | HELIX |
| ALA | 138 | 1.179 | -0.964 | 1.857 | HELIX |
| GLN | 139 | -2.106 | 0.11 | -0.078 |  |
| MET | 140 | -0.16 | 0 | -0.889 |  |
| PRO | 141 | -0.144 | 0.148 | 0.213 |  |
| GLN | 142 | 1.365 | 0.273 | -0.91 |  |
| GLU- | 143 | -2.632 | 0.898 | -1.096 | STRAND |
| GLU- | 144 | -0.94 | -0.09 | -3.221 | STRAND |
| VAL | 145 | -2.587 | 2.696 | -2.239 | STRAND |
| GLU- | 146 | -0.306 | 0.279 | -1.911 | STRAND |
| LEU | 147 | 0.134 | 0 | -0.226 |  |

**Table S8.** Random coil deviation for Cα and Cβ shifts of BRD3-BD1 Q139S.

| Amino acid | | Cα[ppm] | Cβ[ppm] | (Cα-Cβ) [ppm] | Secondary Structure |
| --- | --- | --- | --- | --- | --- |
| GLY | Expression tag residuals | 0 | 0 | 0 |  |
| PRO |  | -0.257 | 0.445 | -0.309 |  |
| LEU |  | -0.283 | -0.059 | -0.349 |  |
| GLY |  | -0.12 | 0 | -0.31 |  |
| SER |  | -0.442 | 0.145 | -0.414 |  |
| GLU- | 25 | -0.599 | -0.065 | -0.614 |  |
| VAL | 26 | -0.903 | -0.183 | -0.877 |  |
| SER | 27 | -0.952 | 0.426 | -0.79 |  |
| ASN | 28 | -0.273 | 0 | -0.55 |  |
| PRO | 29 | 0.258 | 0.256 | -0.106 |  |
| SER | 30 | -0.111 | -0.065 | -0.445 |  |
| LYS+ | 31 | -1.291 | 0 | -0.56 |  |
| PRO | 32 | 0.209 | 0.551 | -0.945 |  |
| GLY | 33 | -1.201 | 0 | -2.142 | STRAND |
| ARG+ | 34 | -1.786 | 3.097 | -2.705 | STRAND |
| LYS+ | 35 | -1.169 | 0.863 | -4.948 | STRAND |
| THR | 36 | -4.253 | 3.677 | -1.267 | STRAND |
| ASN | 37 | 3.048 | -3.112 | 0.062 |  |
| GLN | 38 | 2.805 | 0.849 | 3.988 | HELIX |
| LEU | 39 | 3.246 | -0.601 | 2.837 | HELIX |
| GLN | 40 | 2.81 | 0.101 | 3.7 | HELIX |
| TYR | 41 | 3.672 | -0.871 | 3.528 | HELIX |
| MET | 42 | 1.311 | -2.021 | 3.48 | HELIX |
| GLN | 43 | 3.165 | 0.601 | 2.469 | HELIX |
| ASN | 44 | 1.351 | -0.159 | 2.239 | HELIX |
| VAL | 45 | 2.052 | -0.59 | 4.19 | HELIX |
| VAL | 46 | 6.567 | -1.851 | 6.059 | HELIX |
| VAL | 47 | 5.367 | -1.749 | 6.105 | HELIX |
| LYS+ | 48 | 2.876 | 0.095 | 5.119 | HELIX |
| THR | 49 | 4.651 | -0.809 | 3.694 | HELIX |
| LEU | 50 | 2.92 | 0.08 | 4.25 | HELIX |
| TRP | 51 | 3.131 | -1.32 | 1.928 | HELIX |
| LYS+ | 52 | -1.27 | 0.237 | 1.209 | HELIX |
| HIS | 53 | 2.803 | 2.12 | 0.419 |  |
| GLN | 54 | 2.08 | 0 | -0.009 |  |
| PHE | 55 | -5.072 | -2.282 | -0.511 |  |
| ALA | 56 | 1.025 | 1.847 | -0.185 |  |
| TRP | 57 | 3.057 | 0 | 1.604 | HELIX |
| PRO | 58 | 1.117 | -1.459 | 1.801 |  |
| PHE | 59 | -1.35 | -1.12 | 0.883 |  |
| TYR | 60 | -1.288 | -1.592 | -0.553 |  |
| GLN | 61 | -1.733 | 0 | -0.767 |  |
| PRO | 62 | -0.643 | 0.23 | -0.035 |  |
| VAL | 63 | 2.056 | -0.445 | -0.171 |  |
| ASP- | 64 | -2.806 | -0.664 | 0.47 |  |
| ALA | 65 | 1.304 | 0.253 | 0.748 |  |
| ILE | 66 | 1.437 | -1.899 | 2.104 | HELIX |
| LYS+ | 67 | 1.906 | -0.018 | 1.567 | HELIX |
| LEU | 68 | -0.924 | -0.364 | 1.187 | HELIX |
| ASN | 69 | 0.255 | -1.943 | -0.254 |  |
| LEU | 70 | -2.4 | 0 | 0.424 |  |
| PRO | 71 | 1.289 | -0.186 | -0.362 |  |
| ASP- | 72 | -0.686 | -0.524 | 2.542 | HELIX |
| TYR | 73 | 5.657 | -0.656 | 2.903 | HELIX |
| HIS | 74 | 1.318 | -1.24 | 3.154 | HELIX |
| LYS+ | 75 | 0.518 | -0.073 | 0.881 |  |
| ILE | 76 | 0.648 | 1.155 | 0.093 |  |
| ILE | 77 | -4.519 | -4.713 | -0.21 |  |
| LYS+ | 78 | -0.301 | 0.017 | -0.475 |  |
| ASN | 79 | -1.3 | 0 | -0.683 |  |
| PRO | 80 | -0.179 | 0.253 | -0.909 |  |
| MET | 81 | -2.181 | -1.185 | -2.161 | STRAND |
| ASP- | 82 | -1.715 | 3.339 | -1.654 | STRAND |
| MET | 83 | 3.251 | 2.162 | -0.458 |  |
| GLY | 84 | 2.591 | 0 | 2.858 | HELIX |
| THR | 85 | 4.895 | 0 | 3.818 | HELIX |
| ILE | 86 | 4.282 | 0.315 | 4.135 | HELIX |
| LYS+ | 87 | 4.009 | 0.465 | 3.497 | HELIX |
| LYS+ | 88 | 2.14 | -0.839 | 3.638 | HELIX |
| ARG+ | 89 | 3.481 | -0.91 | 3.471 | HELIX |
| LEU | 90 | 2.657 | -0.385 | 3.125 | HELIX |
| GLU- | 91 | 1.444 | -0.498 | 1.367 | HELIX |
| ASN | 92 | -0.917 | -0.034 | 1.441 | HELIX |
| ASN | 93 | 0.901 | -2.363 | 1.002 | HELIX |
| TYR | 94 | 2.121 | 1.496 | 1.12 | HELIX |
| TYR | 95 | -0.794 | -0.266 | 0.214 |  |
| TRP | 96 | 1.186 | 0.642 | -1.951 | STRAND |
| SER | 97 | -3.961 | 1.908 | -0.979 | STRAND |
| ALA | 98 | 2.087 | -0.3 | -0.107 |  |
| SER | 99 | 3.16 | 0 | 1.595 | HELIX |
| GLU- | 100 | 1.153 | 1.916 | 2.311 | HELIX |
| CYS | 101 | 3.775 | -0.76 | 2.694 | HELIX |
| MET | 102 | 3.811 | -0.5 | 4.111 | HELIX |
| GLN | 103 | 2.479 | -1.009 | 4.232 | HELIX |
| ASP- | 104 | 3.644 | -1.253 | 4.738 | HELIX |
| PHE | 105 | 5.487 | -0.342 | 4.826 | HELIX |
| ASN | 106 | 3.69 | -0.063 | 4.96 | HELIX |
| THR | 107 | 4.379 | -0.919 | 3.936 | HELIX |
| MET | 108 | 3.99 | 1.232 | 4.441 | HELIX |
| PHE | 109 | 3.208 | -2.06 | 4.495 | HELIX |
| THR | 110 | 4.769 | -0.69 | 4.346 | HELIX |
| ASN | 111 | 1.585 | -0.727 | 4.4 | HELIX |
| CYS | 112 | 4.09 | -1.339 | 3.113 | HELIX |
| TYR | 113 | -1.55 | -3.147 | 3.116 | HELIX |
| ILE | 114 | 2.023 | -0.298 | 2.22 | HELIX |
| TYR | 115 | 3.797 | 1.054 | 0.755 |  |
| ASN | 116 | -0.008 | 2.791 | -0.109 |  |
| LYS+ | 117 | -0.271 | 0 | -0.61 |  |
| PRO | 118 | 1.638 | 0.398 | -0.093 |  |
| THR | 119 | -2.003 | -0.755 | -0.015 |  |
| ASP- | 120 | 0.384 | 0.42 | 0.903 |  |
| ASP- | 121 | 3.189 | -0.803 | 2.176 | HELIX |
| ILE | 122 | 1.416 | -1.156 | 4.199 | HELIX |
| VAL | 123 | 4.519 | -1.513 | 4.218 | HELIX |
| LEU | 124 | 2.465 | -1.586 | 4.344 | HELIX |
| MET | 125 | 3.763 | 0.815 | 3.14 | HELIX |
| ALA | 126 | 2.732 | 0.31 | 2.84 | HELIX |
| GLN | 127 | 2.18 | -0.97 | 2.691 | HELIX |
| ALA | 128 | 1.631 | -0.87 | 2.464 | HELIX |
| LEU | 129 | 1.814 | 0.072 | 2.488 | HELIX |
| GLU- | 130 | 1.823 | -1.397 | 2.825 | HELIX |
| LYS+ | 131 | 3.13 | -0.382 | 3.441 | HELIX |
| ILE | 132 | 2.41 | -1.181 | 3.961 | HELIX |
| PHE | 133 | 3.671 | -1.109 | 3.724 | HELIX |
| LEU | 134 | 2.003 | -0.797 | 3.563 | HELIX |
| GLN | 135 | 2.304 | -0.804 | 2.104 | HELIX |
| LYS+ | 136 | -0.286 | -0.691 | 2.966 | HELIX |
| VAL | 137 | 3.912 | -1.472 | 2.712 | HELIX |
| ALA | 138 | 1.378 | -0.969 | 2.167 | HELIX |
| SER | 139 | -1.056 | 0.175 | 0.292 |  |
| MET | 140 | -0.24 | 0 | -0.613 |  |
| PRO | 141 | -0.161 | 0.208 | 0.14 |  |
| GLN | 142 | 1.339 | 0.311 | -0.974 | STRAND |
| GLU- | 143 | -2.667 | 0.915 | -1.094 | STRAND |
| GLU- | 144 | -0.921 | -0.192 | -3.199 | STRAND |
| VAL | 145 | -2.588 | 2.697 | -2.197 | STRAND |
| GLU- | 146 | -0.295 | 0.282 | -1.903 | STRAND |
| LEU | 147 | 0.154 | 0 | -0.211 |  |

**Table S9.** Random coil deviation for Cα and Cβ shifts of BRD3-BD1 V145Q.

| Amino Acid | | Cα[ppm] | Cβ[ppm] | (Cα-Cβ)[ppm] | Secondary Structure |
| --- | --- | --- | --- | --- | --- |
| GLY | Expression tag residuals | 0 | 0 | 0 |  |
| PRO |  | -0.267 | 0.435 | -0.31 |  |
| LEU |  | -0.288 | -0.059 | -0.351 |  |
| GLY |  | -0.123 | 0 | -0.314 |  |
| SER |  | -0.443 | 0.148 | -0.419 |  |
| GLU- | 25 | -0.609 | -0.066 | -0.618 |  |
| VAL | 26 | -0.907 | -0.187 | -0.886 |  |
| SER | 27 | -0.962 | 0.434 | -0.8 |  |
| ASN | 28 | -0.283 | 0 | -0.555 |  |
| PRO | 29 | 0.264 | 0.251 | -0.096 |  |
| SER | 30 | -0.175 | -0.156 | -0.433 |  |
| LYS+ | 31 | -1.293 | 0 | -0.551 |  |
| PRO | 32 | 0.193 | 0.533 | -0.939 |  |
| GLY | 33 | -1.183 | 0 | -2.13 | STRAND |
| ARG+ | 34 | -1.837 | 3.029 | -2.7 | STRAND |
| LYS+ | 35 | -1.191 | 0.86 | -4.911 | STRAND |
| THR | 36 | -4.272 | 3.544 | -1.236 | STRAND |
| ASN | 37 | 2.961 | -3.197 | 0.078 |  |
| GLN | 38 | 2.749 | 0.856 | 3.944 | HELIX |
| LEU | 39 | 3.263 | -0.519 | 2.791 | HELIX |
| GLN | 40 | 2.789 | 0.09 | 3.68 | HELIX |
| TYR | 41 | 3.663 | -0.897 | 3.533 | HELIX |
| MET | 42 | 1.306 | -2.035 | 3.474 | HELIX |
| GLN | 43 | 3.137 | 0.617 | 2.465 | HELIX |
| ASN | 44 | 1.332 | -0.201 | 2.166 | HELIX |
| VAL | 45 | 2.043 | -0.402 | 4.222 | HELIX |
| VAL | 46 | 6.672 | -2.016 | 6.083 | HELIX |
| VAL | 47 | 5.346 | -1.771 | 6.197 | HELIX |
| LYS+ | 48 | 2.896 | 0.111 | 5.127 | HELIX |
| THR | 49 | 4.695 | -0.784 | 3.744 | HELIX |
| LEU | 50 | 2.911 | -0.056 | 4.292 | HELIX |
| TRP | 51 | 3.152 | -1.278 | 1.988 | HELIX |
| LYS+ | 52 | -1.198 | 0.236 | 1.226 | HELIX |
| HIS | 53 | 2.785 | 2.104 | 0.441 |  |
| GLN | 54 | 2.077 | 0 | 0.033 |  |
| PHE | 55 | -5.053 | -2.394 | -0.483 |  |
| ALA | 56 | 1.015 | 1.883 | -0.162 |  |
| TRP | 57 | 3.042 | 0 | 1.583 | HELIX |
| PRO | 58 | 1.146 | -1.43 | 1.804 |  |
| PHE | 59 | -1.359 | -1.152 | 0.903 |  |
| TYR | 60 | -1.258 | -1.599 | -0.533 |  |
| GLN | 61 | -1.733 | 0 | -0.807 |  |
| PRO | 62 | -0.73 | 0.3 | -0.119 |  |
| VAL | 63 | 1.986 | -0.42 | -0.235 |  |
| ASP- | 64 | -2.802 | -0.721 | 0.45 |  |
| ALA | 65 | 1.288 | 0.262 | 0.747 |  |
| ILE | 66 | 1.48 | -1.816 | 2.111 | HELIX |
| LYS+ | 67 | 1.921 | -0.089 | 1.579 | HELIX |
| LEU | 68 | -0.909 | -0.34 | 1.222 | HELIX |
| ASN | 69 | 0.265 | -1.96 | -0.258 |  |
| LEU | 70 | -2.431 | 0 | 0.377 |  |
| PRO | 71 | 1.015 | -0.321 | -0.406 |  |
| ASP- | 72 | -0.654 | -0.532 | 2.288 | HELIX |
| TYR | 73 | 5.707 | 0.057 | 2.637 | HELIX |
| HIS | 74 | 1.252 | -1.132 | 2.847 | HELIX |
| LYS+ | 75 | 0.506 | 0 | 0.858 |  |
| ILE | 76 | 0.564 | 0.881 | 0.237 |  |
| ILE | 77 | -4.527 | -5.05 | -0.046 |  |
| LYS+ | 78 | -0.325 | 0.018 | 2.556 | HELIX |
| ASN | 79 | -1.292 | -8.781 | 2.238 | HELIX |
| PRO | 80 | -0.173 | 0.259 | 2.011 |  |
| MET | 81 | -2.178 | -1.155 | -2.175 | STRAND |
| ASP- | 82 | -1.711 | 3.36 | -1.676 | STRAND |
| MET | 83 | 3.262 | 2.196 | -0.482 |  |
| GLY | 84 | 2.558 | 0 | 2.834 | HELIX |
| THR | 85 | 4.879 | 0 | 3.787 | HELIX |
| ILE | 86 | 4.277 | 0.354 | 4.149 | HELIX |
| LYS+ | 87 | 4.006 | 0.361 | 3.43 | HELIX |
| LYS+ | 88 | 2.15 | -0.571 | 3.544 | HELIX |
| ARG+ | 89 | 3.5 | -0.766 | 3.367 | HELIX |
| LEU | 90 | 2.648 | -0.467 | 3.109 | HELIX |
| GLU- | 91 | 1.438 | -0.508 | 1.404 | HELIX |
| ASN | 92 | -0.877 | -0.029 | 1.451 | HELIX |
| ASN | 93 | 0.895 | -2.361 | 1.017 | HELIX |
| TYR | 94 | 2.127 | 1.485 | 1.096 | HELIX |
| TYR | 95 | -0.823 | -0.213 | 0.177 |  |
| TRP | 96 | 1.125 | 0.625 | -2.01 | STRAND |
| SER | 97 | -3.908 | 2.012 | -1.027 | STRAND |
| ALA | 98 | 2.059 | -0.281 | 0.488 |  |
| SER | 99 | 3.155 | -1.89 | 2.211 | HELIX |
| GLU- | 100 | 1.176 | 1.927 | 2.952 | HELIX |
| CYS | 101 | 3.783 | -0.778 | 2.7 | HELIX |
| MET | 102 | 3.864 | -0.426 | 4.116 | HELIX |
| GLN | 103 | 2.48 | -1.017 | 4.25 | HELIX |
| ASP- | 104 | 3.702 | -1.261 | 4.746 | HELIX |
| PHE | 105 | 5.47 | -0.308 | 4.828 | HELIX |
| ASN | 106 | 3.691 | -0.053 | 4.931 | HELIX |
| THR | 107 | 4.382 | -0.889 | 3.92 | HELIX |
| MET | 108 | 3.973 | 1.228 | 4.399 | HELIX |
| PHE | 109 | 3.199 | -1.982 | 4.461 | HELIX |
| THR | 110 | 4.768 | -0.688 | 4.319 | HELIX |
| ASN | 111 | 1.599 | -0.722 | 4.392 | HELIX |
| CYS | 112 | 4.051 | -1.347 | 3.098 | HELIX |
| TYR | 113 | -1.558 | -3.134 | 3.004 | HELIX |
| ILE | 114 | 2.033 | -0.005 | 2.118 | HELIX |
| TYR | 115 | 3.808 | 1.067 | 0.665 |  |
| ASN | 116 | 0.01 | 2.794 | -0.106 |  |
| LYS+ | 117 | -0.274 | 0 | -0.579 |  |
| PRO | 118 | 1.666 | 0.344 | -0.086 |  |
| THR | 119 | -2.028 | -0.723 | -0.011 |  |
| ASP- | 120 | 0.377 | 0.426 | 0.872 |  |
| ASP- | 121 | 3.186 | -0.784 | 2.167 | HELIX |
| ILE | 122 | 1.448 | -1.132 | 4.194 | HELIX |
| VAL | 123 | 4.519 | -1.514 | 4.196 | HELIX |
| LEU | 124 | 2.436 | -1.539 | 4.268 | HELIX |
| MET | 125 | 3.612 | 0.817 | 3.022 | HELIX |
| ALA | 126 | 2.727 | 0.43 | 2.423 | HELIX |
| GLN | 127 | 2.177 | 0 | 2.316 | HELIX |
| ALA | 128 | 1.708 | -0.765 | 2.128 | HELIX |
| LEU | 129 | 1.832 | 0.098 | 2.474 | HELIX |
| GLU- | 130 | 1.795 | -1.421 | 2.825 | HELIX |
| LYS+ | 131 | 3.131 | -0.393 | 3.46 | HELIX |
| ILE | 132 | 2.442 | -1.197 | 3.982 | HELIX |
| PHE | 133 | 3.674 | -1.108 | 3.744 | HELIX |
| LEU | 134 | 2.04 | -0.77 | 3.56 | HELIX |
| GLN | 135 | 2.296 | -0.793 | 2.086 | HELIX |
| LYS+ | 136 | -0.357 | -0.715 | 3.053 | HELIX |
| VAL | 137 | 4.182 | -1.529 | 2.725 | HELIX |
| ALA | 138 | 1.165 | -0.942 | 1.87 | HELIX |
| GLN | 139 | -2.21 | -0.002 | -0.088 |  |
| MET | 140 | -0.162 | 0 | -0.832 |  |
| PRO | 141 | -0.095 | 0.031 | 0.368 |  |
| GLN | 142 | 1.391 | -0.002 | -0.776 |  |
| GLU- | 143 | -2.682 | 0.914 | -0.799 |  |
| GLU- | 144 | -0.483 | -0.289 | -3.219 | STRAND |
| GLN | 145 | -2.705 | 3.162 | -2.147 | STRAND |
| GLU- | 146 | -0.167 | 0.212 | -1.971 | STRAND |
| LEU | 147 | 0.333 | 0 | -0.023 |  |

**FIGURES**

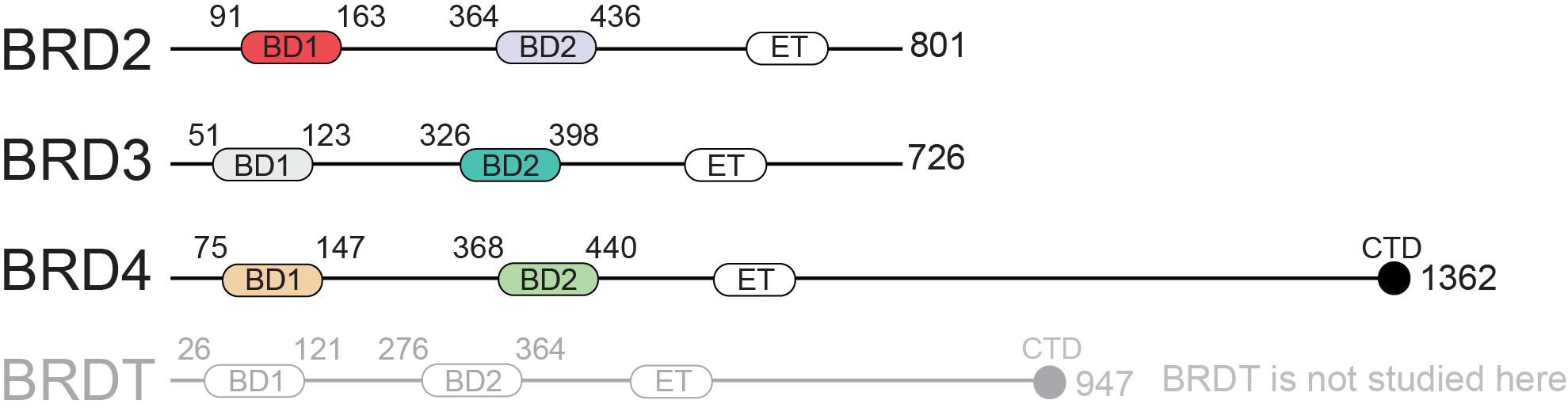

**Figure S1. Domain architecture of the Bromodomain and ExtraTerminal domain (BET) family of proteins.** The domain topologies of the human BET proteins are shown. The positioning of the first (BD1) and second (BD2) bromodomains, extraterminal domains (ET), and the *C*-terminal domain (CTD) found in BRD4 and BRDT is shown. The BDs used in this study are coloured differently to distinguish them and the colouring used to display structures of the BDs throughout this paper is consistent with this scheme.

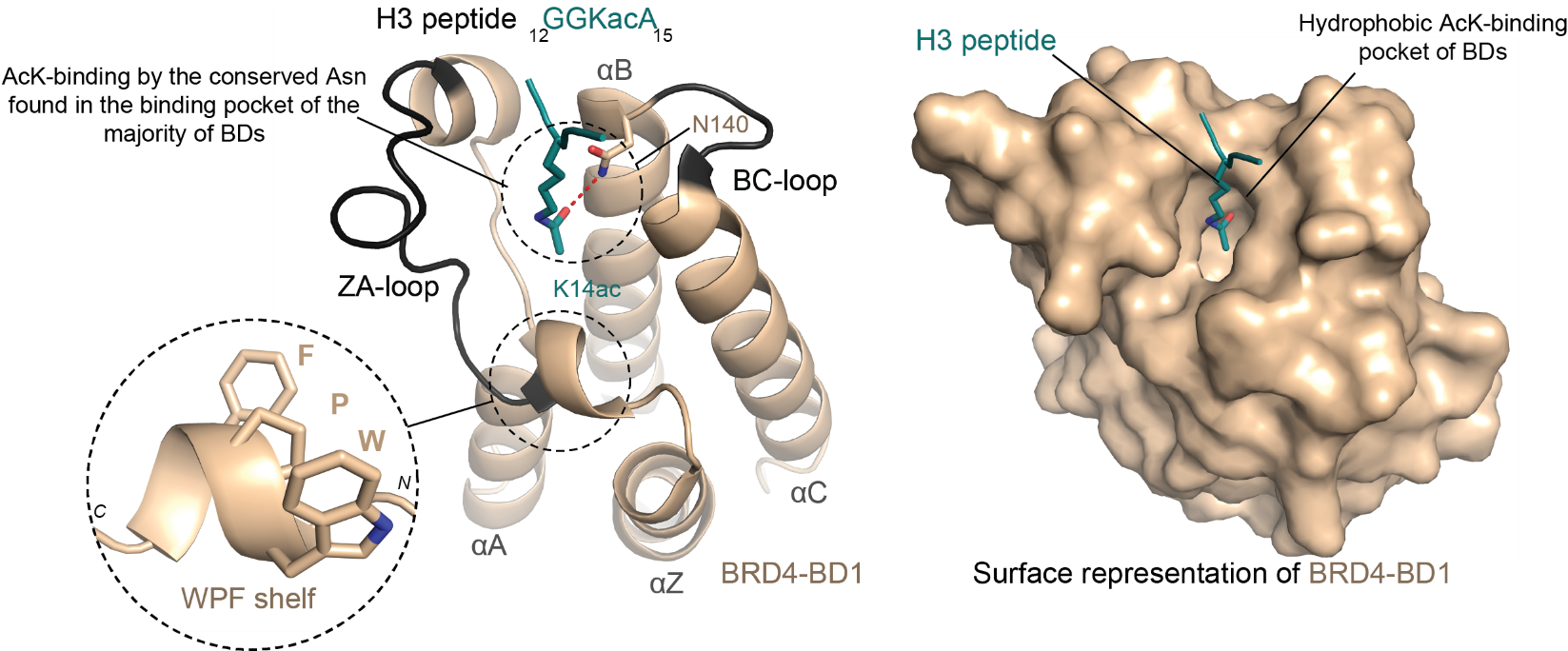

**Figure S2. Overview of bromodomain (BD) structure and acetyllysine (AcK) binding using the first bromodomain of BRD4 (BRD4-BD1) as an example.** The crystal structure of a bromodomain (BRD4-BD1; PDB ID: 3JVK) in complex with an acetylated histone H3 peptide^1^. *Left*: Ribbon representation of BRD4-BD1 with the four α-helices (αZ, αA, αB, and αD) labelled. The helices are connected by two flexible loop regions: the ZA loop that links the αZ and αA helices and the BC loop that links the αB and αC helices. The acetylated H3 peptide occupying the BD-binding pocket is circled and displayed in *turquoise* with the AcK residue represented as sticks. The hydrogen bond that forms between the acetyl group of AcK residues and the sidechain amide of a conserved asparagine in the binding pocket of BDs is indicated by the *red* dashed line. The “WPF shelf”, which is located at the surface of the AcK-binding pocket and is involved in engaging native BD substrates, is also circled and shown as an inset. *Right*: Surface representation of BRD4-BD1, illustrating the AcK-binding cavity.

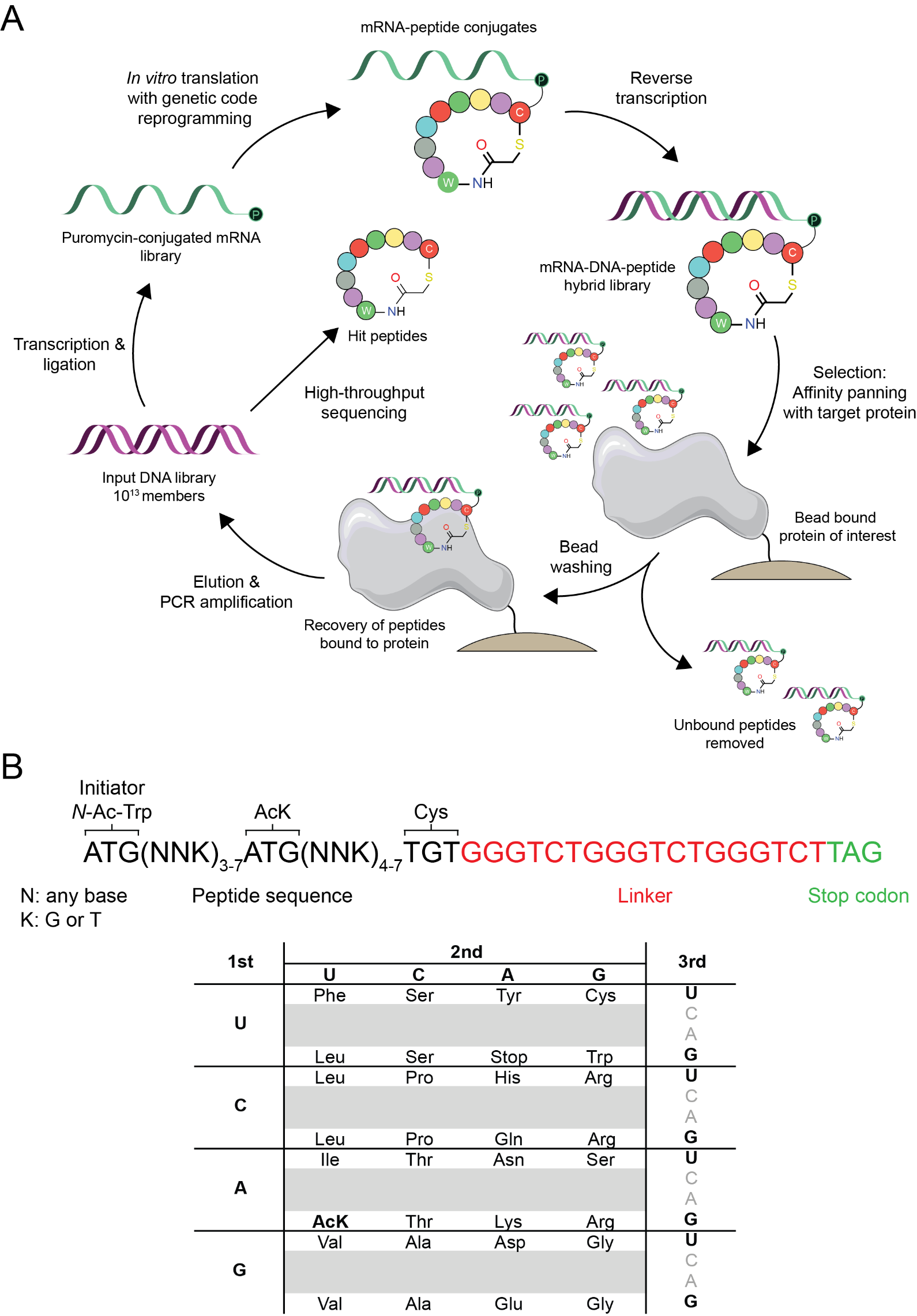

**Figure S3. Overview of the RaPID mRNA display system. A.** Schematic of the RaPID selection process. **B.** Library design and codon assignment used in the RaPID selections in this study.

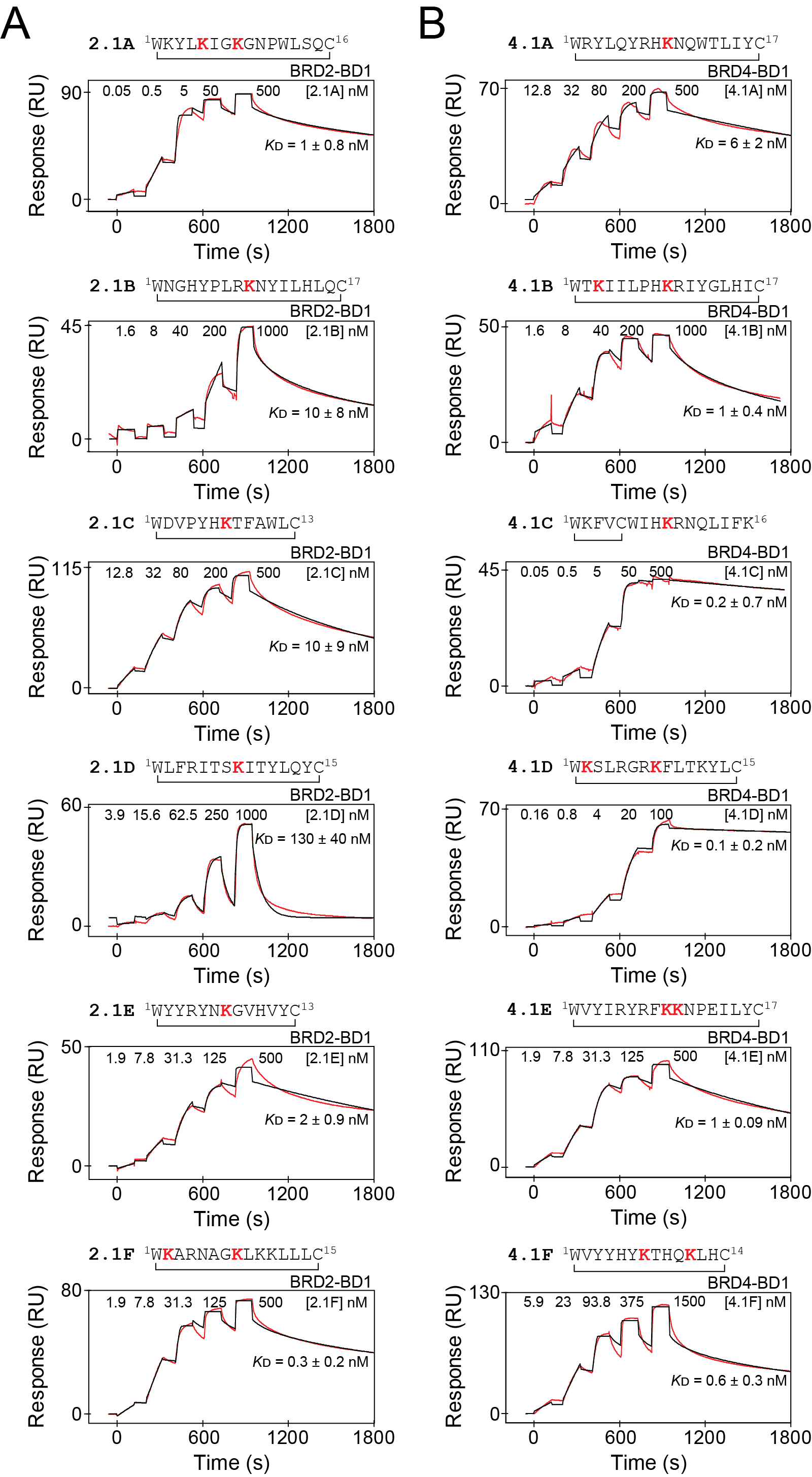

**Figure S4. SPR binding analyses for CPs selected against BRD2-BD1 and BRD4-BD1.** Representative SPR sensorgrams (*red*) for binding of CPs to the domain against which they were selected. **A** shows CPs selected against BRD2-BD1 and **B** shows peptides selected against BRD4-BD1. Fits to a 1:1 binding model (*black*) are shown for each sensorgram. The *K*_D_ values for the interactions are provided and further summarised in **Table S3**.

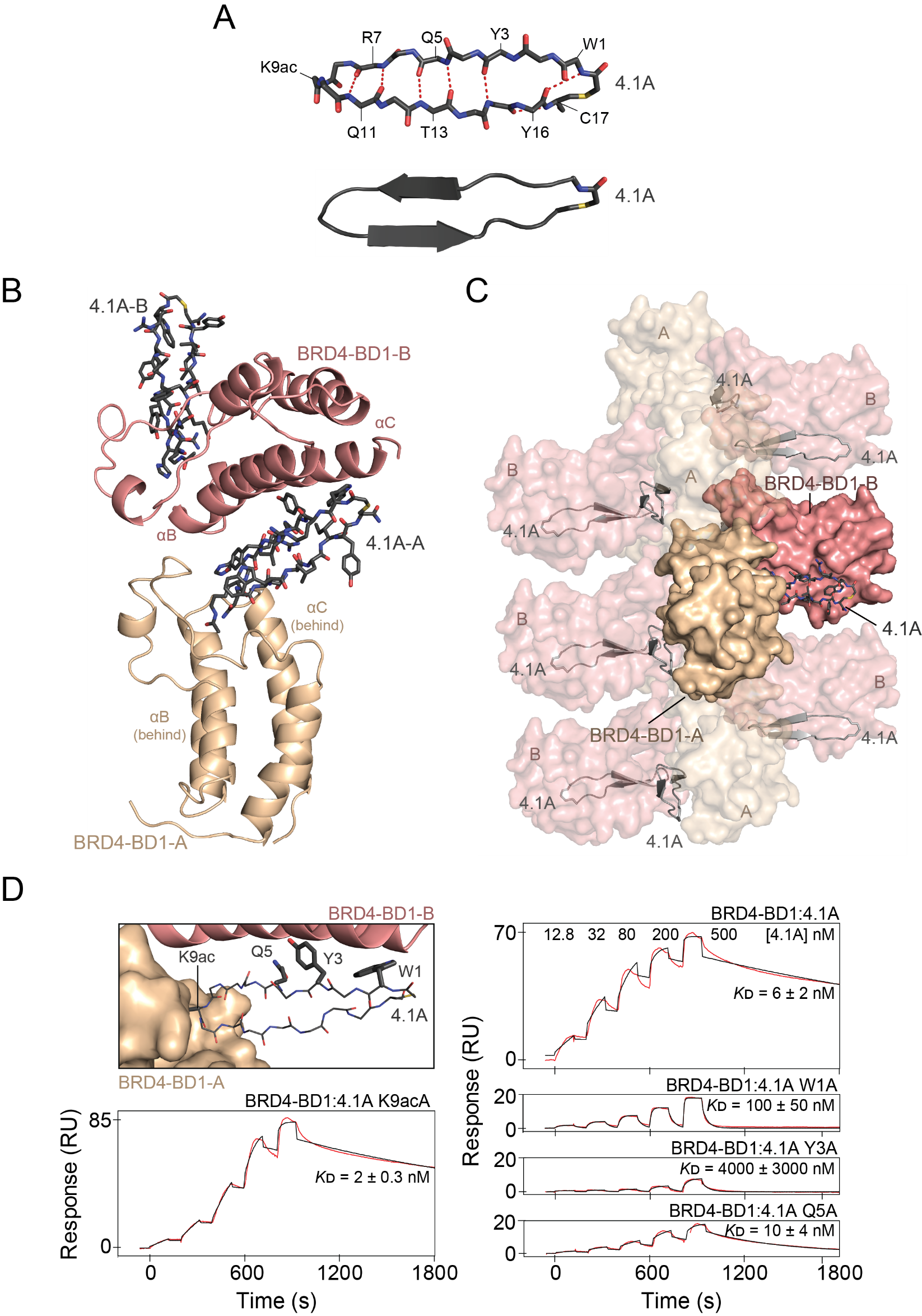

**Figure S5. Structural details of the complex formed between BRD4-BD1 and 4.1A. A.** Backbone structure of 4.1A in complex with BRD4-BD1. The intra-mainchain hydrogen bonds are indicated by *red dashed lines*. The cartoon structure of the peptide is shown below. **B.** Ribbon representation of the two BRD4-BD1 molecules in the structure presented in **Fig. 2A**. The αB and αC helices are labelled and the second 4.1A that binds the AcK-binding pocket of BRD4-BD1-B is shown as sticks. **C.** A section of the filamentous structure formed by the BRD4-BD1:4.1A complex from **Fig. 2B** shown from a different orientation. **D.** *Top left*: A section of the X-ray crystal structure of BRD4-BD1:4.1A. Residues mutated to alanine are displayed as thick sticks. *Bottom left* and *right*: Representative SPR sensorgrams (*red*) for the binding of the 4.1A alanine mutants to BRD4-BD1. The SPR data for the BRD4-BD1 interaction with ‘wild-type’ 4.1A are also shown for comparison. Fits to a 1:1 binding model (*black*) are shown. The *K*_D_ values are provided as the geometric mean (± standard error) of at least three independent measurements.

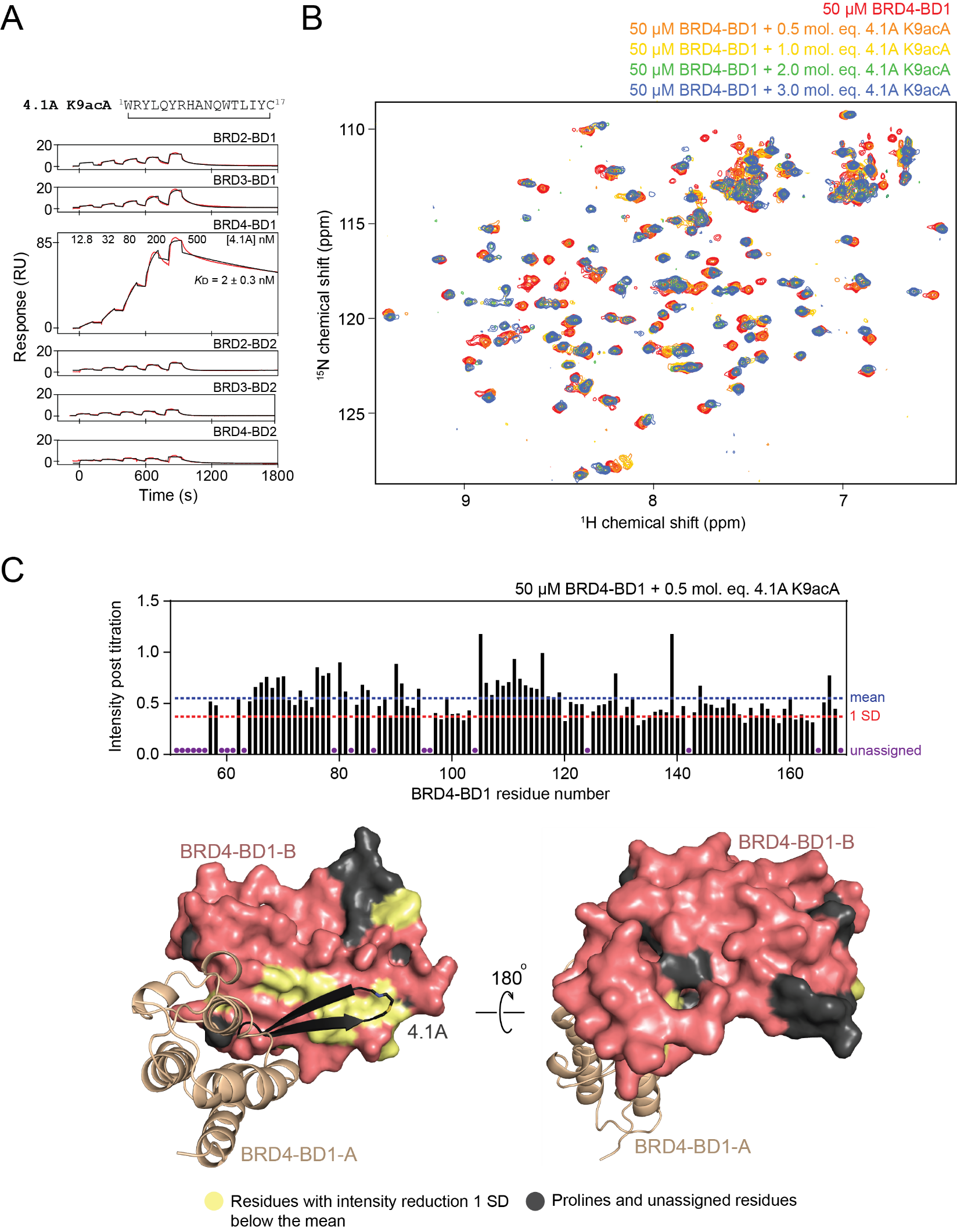

**Figure S6. 4.1A-K9acA binds predominantly to the αB-αC surface of BRD4-BD1.** **A.** Representative SPR sensorgrams (*red*) for the binding of 4.1A K9ac to all BDs of BRD2, BRD3, and BRD4. Fits to a 1:1 binding model (*black*) are shown for each sensorgram. The *K*_D_ for the interaction between 4.1A K9ac and BRD4-BD1 is provided. The remainder of the BDs display no binding to 4.1A K9ac. **B.** ^15^N-HSQC spectra of BRD4-BD1 alone (*red*; 50 μM) and in the presence of increasing concentrations of 4.1A-K9acA (*blue*; up to 3 molar equivalents). **C.** *Top*: Quantitation of ^15^N-HSQC signal intensity for BRD4-BD1 after the addition of 0.5 molar equivalents of 4.1A-K9acA. The mean intensity across all assignable signals in the titration is marked by the *blue dashed line* and the intensity one standard deviation (SD) below the mean is indicated by the *red dashed line*. Prolines or unassigned residues in BRD4-BD1 are indicated by *purple circles*. *Bottom*: Residues for which signals are reduced in intensity by at least 1 SD from the mean are mapped onto the structure of BRD4-BD1 (*salmon*) in *pale yellow*. Prolines and unassigned residues are coloured *dark grey*.

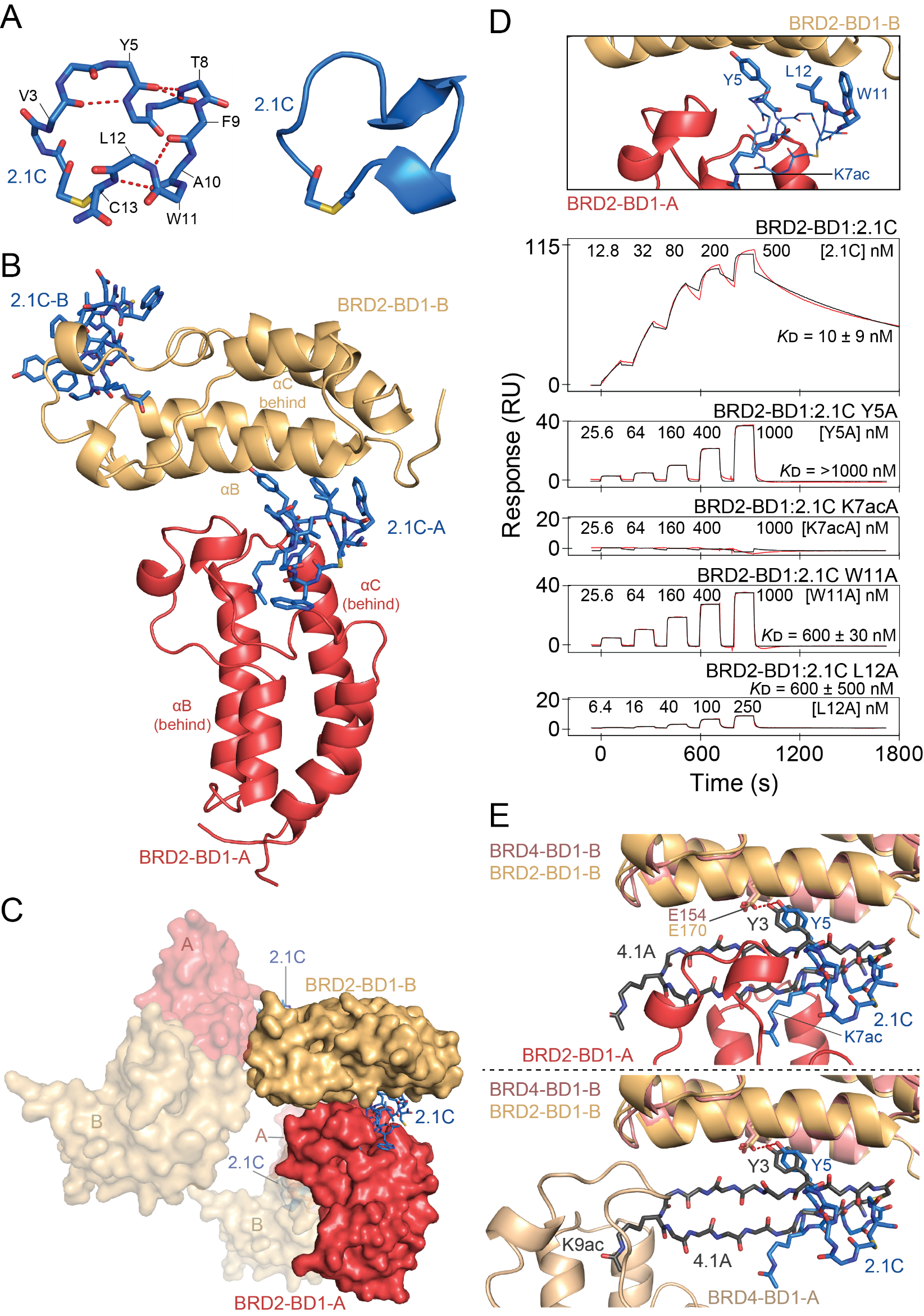

**Figure S7. Structural details of the complex formed between BRD2-BD1 and 2.1C. A.** Backbone structure of 2.1C when in complex with BRD2-BD1. Intra-mainchain hydrogen bonds are indicated by *red dashed lines*. **B.** Ribbon representation of the two BRD2-BD1 molecules in the structure presented in **Fig. 3A**. The αB and αC helices are labelled and the second 2.1C that binds the AcK-binding pocket of BRD2-BD1-B is also shown. **C.** A section of the filamentous structure formed by the BRD2-BD1:2.1C complex, shown in a different orientation to **B**. **D.** *Top*: A section of the X-ray crystal structure of BRD2-BD1:2.1C with the residues mutated to alanine for testing by SPR displayed in thick sticks. *Bottom*: Representative SPR sensorgrams (*red*) for the binding of the 2.1C alanine mutants to BRD2-BD1. SPR data for the BRD2-BD1 2.1C interaction (from **Fig. 1C**) is shown for comparison. Fits to a 1:1 binding model (*black*) are shown. The *K*_D_ values for interactions are provided as the geometric mean (± standard error) of at least three independent measurements. **E.** Overlay of the second BDs and peptides in the X-ray crystal structures of the BRD2-BD1:2.1C and BRD4-BD1:4.1A complexes, showing the shared interaction of a BD glutamate (E154 in BRD4-BD1 and E170 in BRD2-BD1) with a peptide tyrosine (Y3 in BRD4-BD1 and Y5 in BRD2-BD1). The same view is shown in both panels, but the *top* panel shows the position of BRD2-BD1-A from the BRD2-BD1:2.1C complex, whereas the *bottom* panel shows the position of BRD4-BD1-A from the BRD4-BD1:4.1A complex.

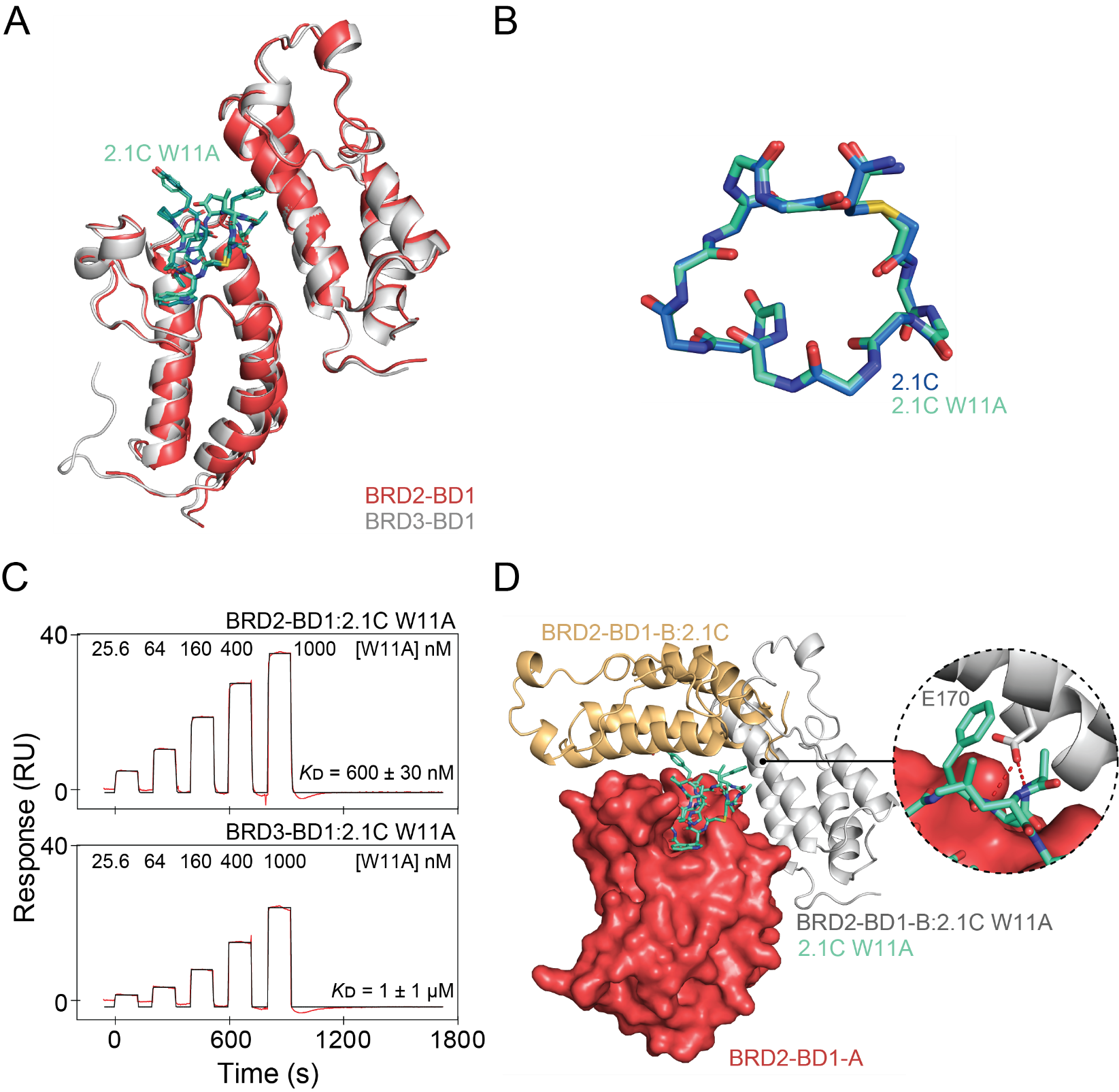

**Fig S8. Characterisation of the 2.1C W11A peptide mutant. A**. Overlay of the BRD2-BD1:2.1C-W11A (BDs as *coral* ribbons, 2.1C-W11A in *teal* sticks; 3.1 Å resolution, PDB ID: 9MPL) and BRD3-BD1:2.1A-W11A (BDs as *grey* ribbons; 2.7 Å resolution, PDB ID: 9MPM) complexes. **B.** Overlay of the backbone structures of 2.1C and 2.1C W11A. **C.** Representative SPR sensorgrams (*red*) for the binding of 2.1C W11A to BRD2-BD1 and BRD3-BD1. Fits of a 1:1 binding mode (*black*) are shown. The *K*_D_ values are provided as the geometric mean (± standard error) of at least two independent measurements. **D**. Comparison of the orientation of BRD2-BD1-B when in complex with 2.1C (*light orange*) and 2.1C-W11A (*grey*). *Left inset*: Close up of the hydrogen bonds (*red dashed lines*) formed between E170 of BRD2-BD1-B and the backbone of 2.1C-W11A.

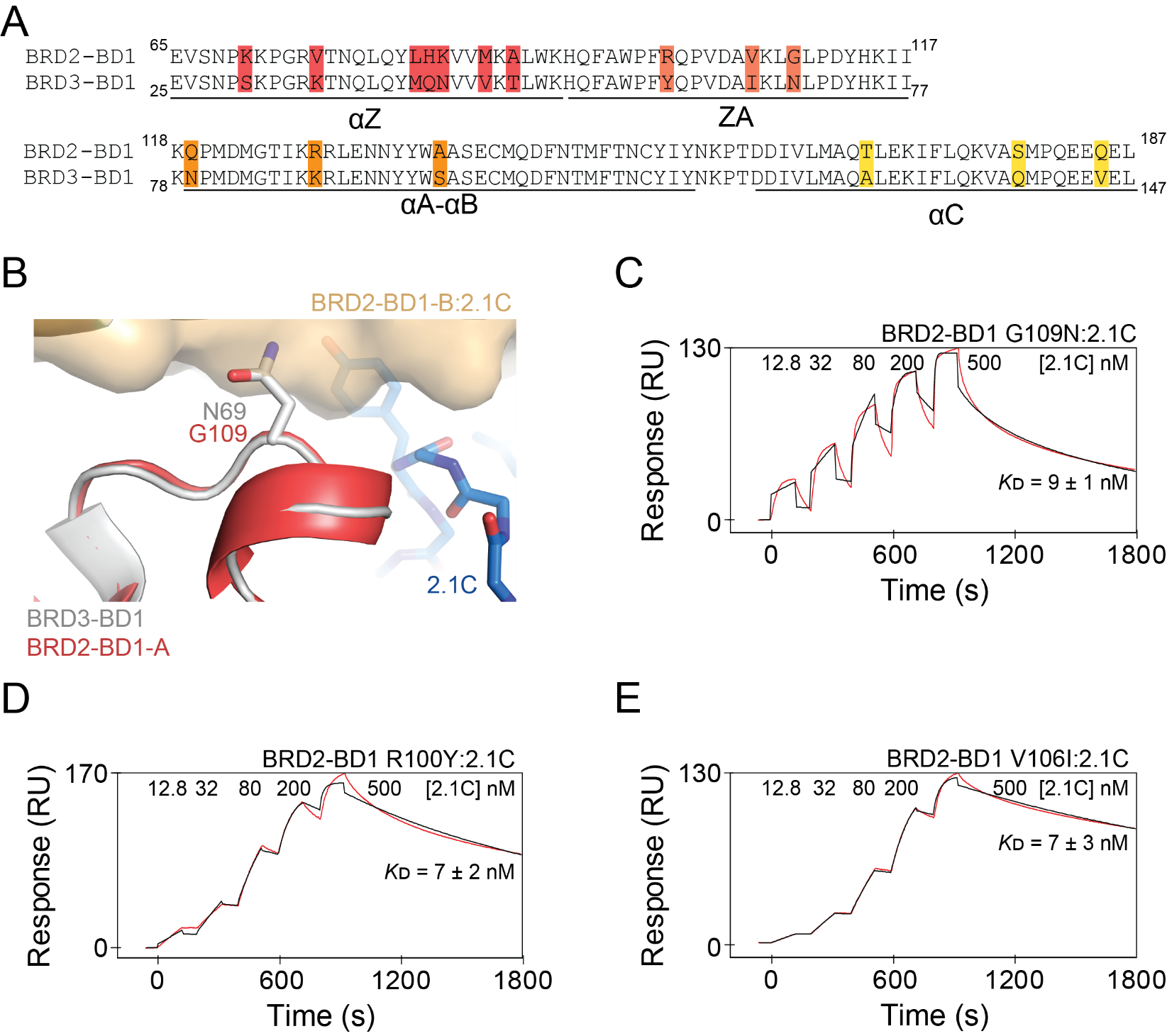

**Figure S9. Mutations in the ZA loop of BRD2-BD1 do not impair binding to 2.1C.** **A.** Sequence alignment of BRD2-BD1 and BRD3-BD1. Non-conserved residues, which were mutated to make the chimeras in **Fig. 3E,** are highlighted and coloured according to their location in the BD. **B**. Overlay of BRD3-BD1 (*grey*) onto the structure of BRD2-BD1 in complex with 2.1C. N69 of BRD3-BD1, which could potentially clash with the BD:BD interaction interface in the BRD2-BD1:2.1C complex, is represented as sticks. **C.** Representative SPR sensorgrams (*red*) for the binding of 2.1C to BRD2-BD1 G109N. **D.** Representative SPR sensorgram (*red*) for the binding of 2.1C to BRD2-BD1 R100Y. **E.** Representative SPR sensorgrams (*red*) for the binding of 2.1C to BRD2-BD1 V106I. For all SPR data in the Figure, fits to a 1:1 binding model (*black*) are shown and *K*_D_ values are provided as the geometric mean (± standard error) of at least three independent measurements.

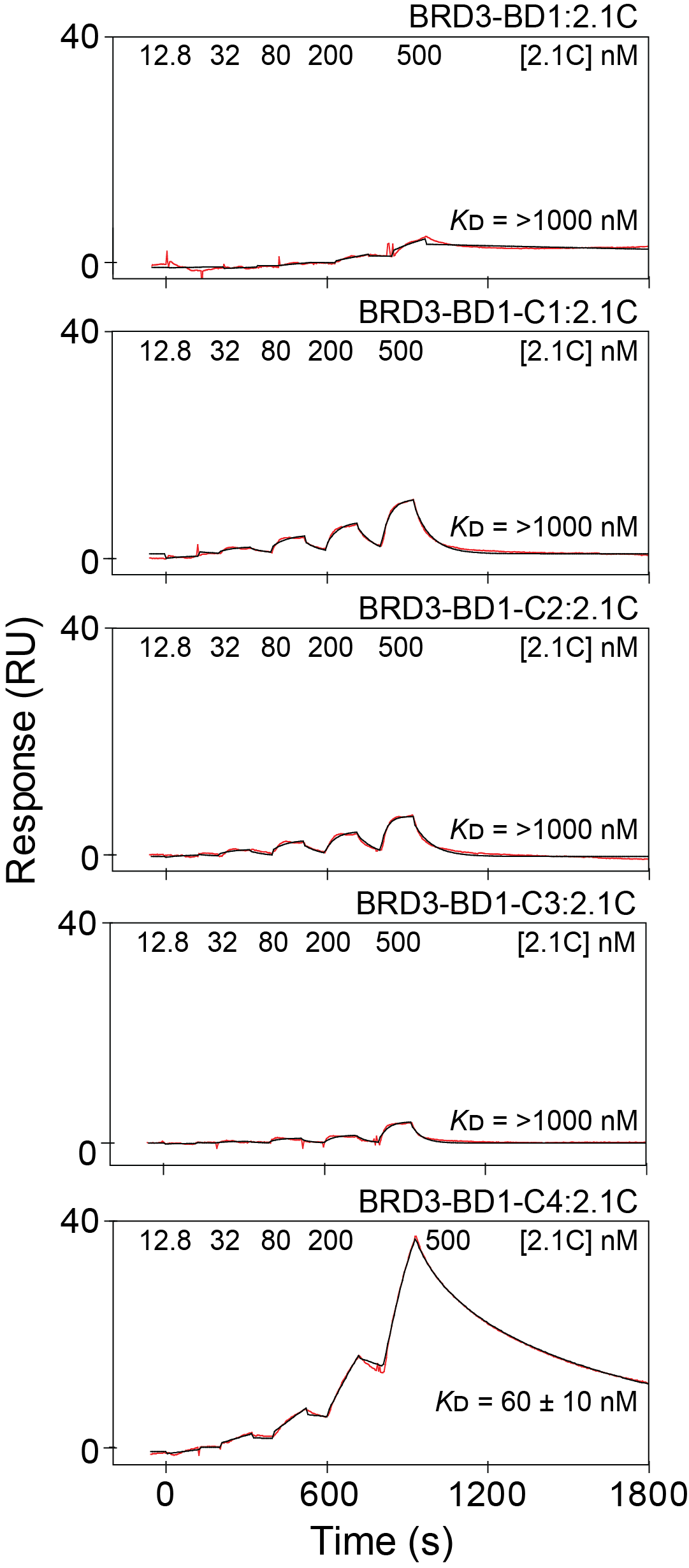

**Figure 10.** Representative SPR sensorgram (*red*) for the binding of 2.1C to BRD2-BD1 and BRD3-BD1 chimeras (shown in **Fig. 3E**. Fits to a 1:1 binding model (*black*) are shown and *K*_D_ values are provided as the geometric mean (± standard error) of at least three independent measurements.

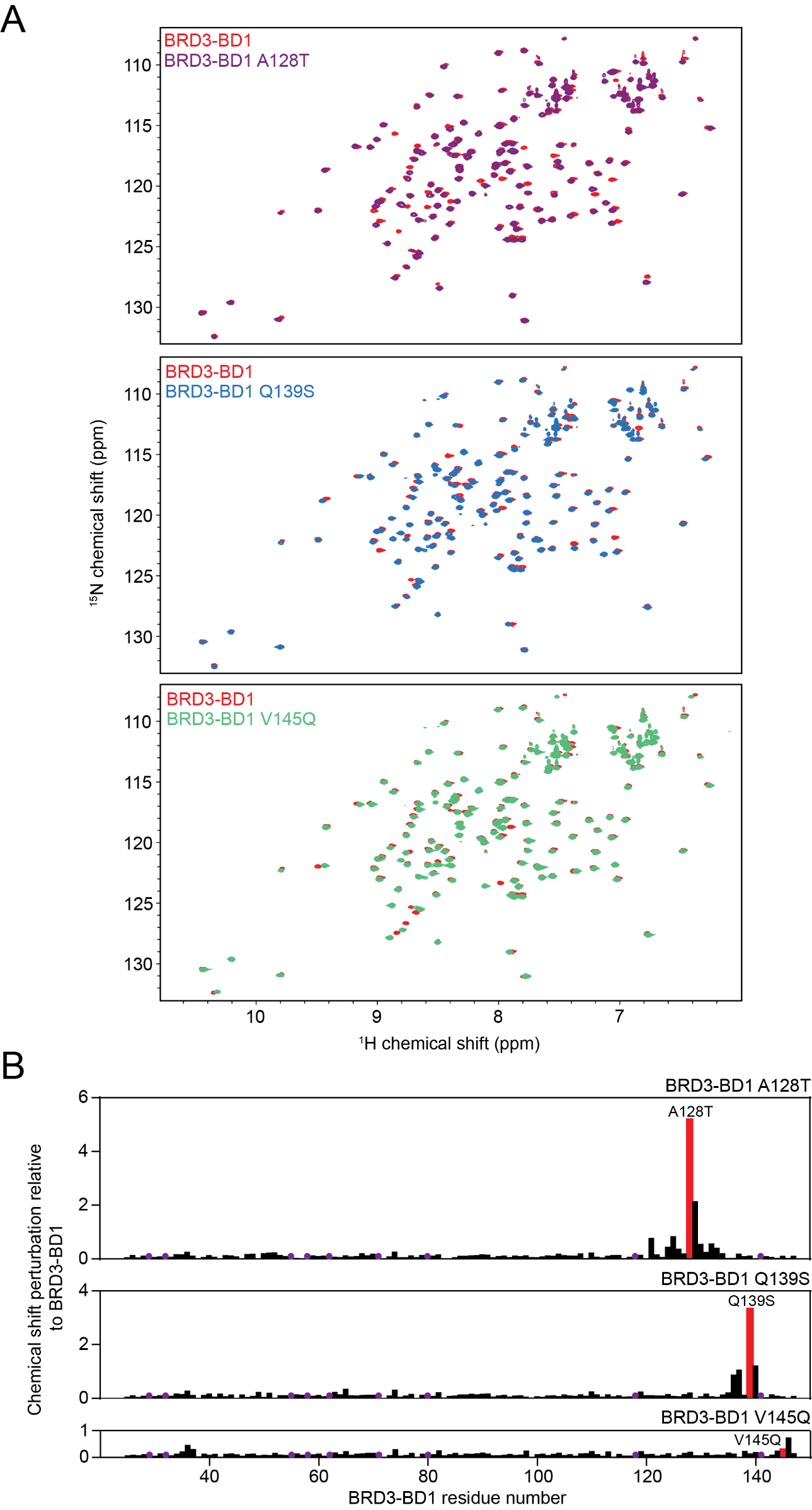

**Figure S11. The fold of BRD3-BD1 aC point mutants is not perturbed. A.** ^15^N-HSQC spectra of BRD3-BD1 (*red*) overlayed with the spectra of BRD3-BD1 A128T (*purple*), BRD3-BD1 Q139S (*blue*), and BRD3-BD1 V145Q (*green*). **B.** Quantitation of the chemical shift perturbations (CSPs) for each mutant relative to BRD3-BD1. The CSP for each mutated residue is coloured *red*. Prolines and unassigned residues are indicated by purple circles.

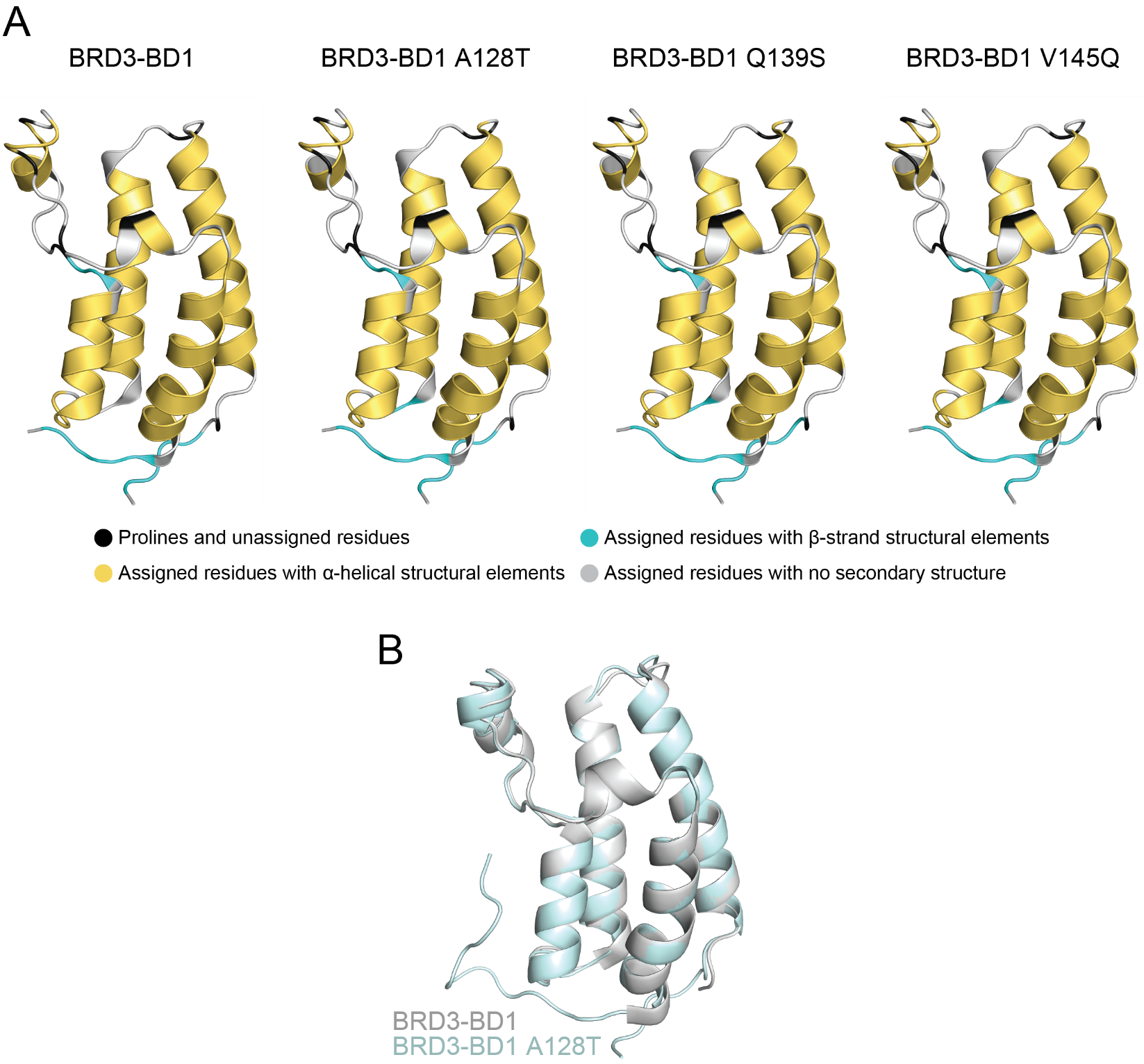

**Figure S12. Secondary structure analysis of BRD3-BD1 mutants. A.** Secondary structure analysis of BRD3-BD1, BRD3-BD1 A128T, BRD3-BD1 Q139S, and BRD3-BD1 V145Q generated from calculating the random coil deviations for CA and CB shifts (provided in **Tables S6-9**). **B**. Overlay of the crystal structure of BRD3-BD1 A128T (*pale cyan*, 1.6 Å resolution, PDB ID: 9MPN) and wildtype BRD3-BD1.

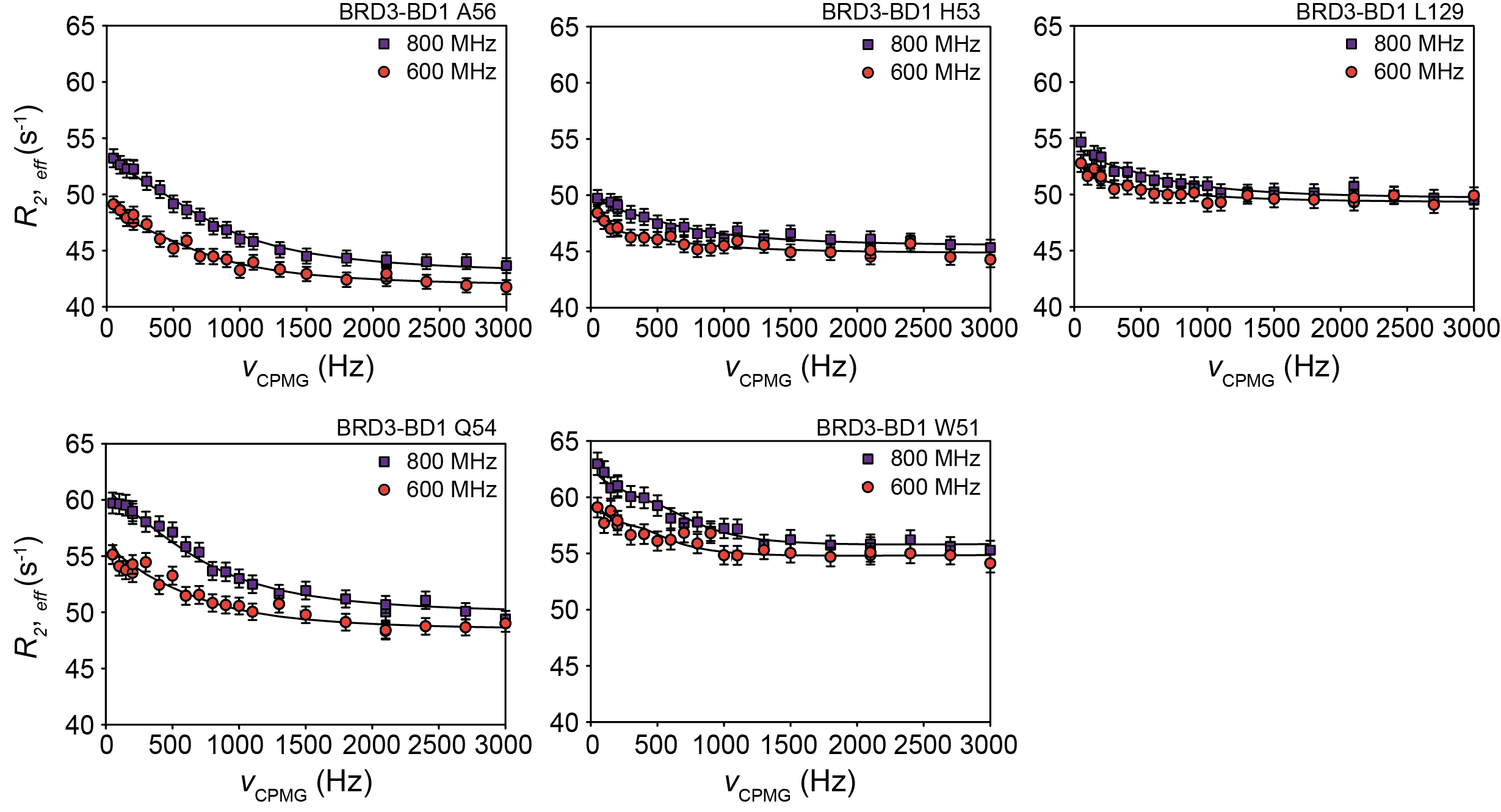

**Figure S13. Representative relaxation dispersion profiles of BRD3-BD1 at site 1.** Effective relaxation rates (*R_2, eff_*) plotted as a function of CPMG frequency (ν_CPMG_). ^1^H^N^ CPMG relaxation dispersion data were acquired at two magnet fields (600 MHz and 800 MHz) at 15 °C and globally fitted to a two-state exchange model using ChemEx. The exchange rate (*k_ex_*) and the minor population (*p_B_*) at this site were estimated to be 4025 ± 243 s^-1^ and 0.77 ± 0.11%, respectively.

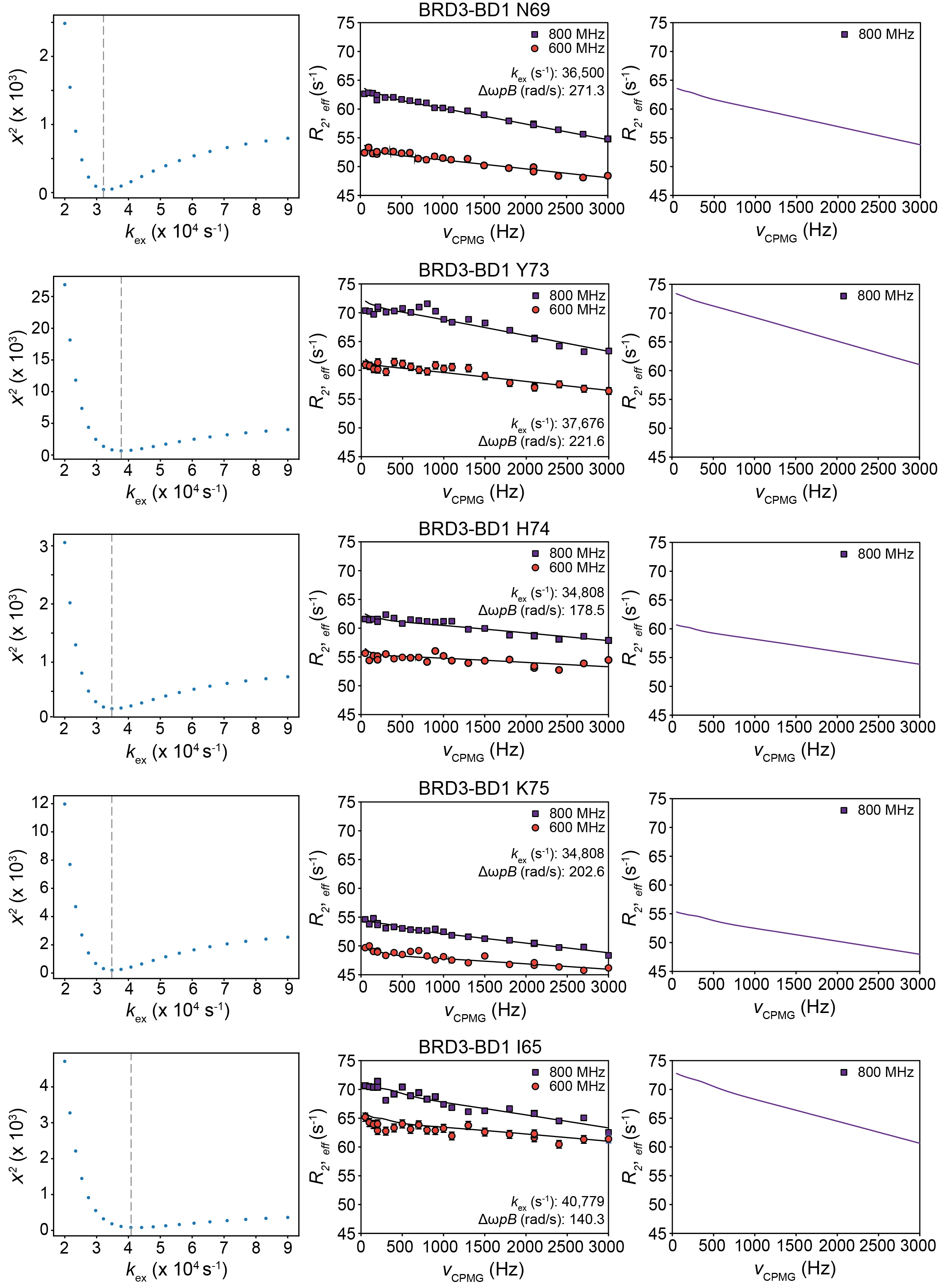

**Figure S14. Relaxation dispersion data analysis of BRD3-BD1 at site 2.** (Left panel) The χ^2^ was evaluated as a function of the exchange rate (*k_ex_*) while keeping the minor population (*p_B_*) and the chemical shift difference of major and minor conformation (Δω) fixed. This analysis was performed individually for each residue, yielding global minima in the range of ~ 34800-41000 s^-1^. For χ^2^ fitting, the ^1^H^N^ CPMG relaxation dispersion data acquired on the 600 MHz and 800 MHz at 15 °C were provided as input and the analysis was performed by ChemEx with a two-state exchange model. (Middle panel) ^1^H^N^ CPMG relaxation dispersion data at two magnetic fields, as fitted during χ^2^ evaluation. The mean exchange rate (*k_ex_*) at this site was estimated to be 36900 ± 2477 s^-1^. (Right panel) The CPMG relaxation dispersion data for each residue at 800 MHz was simulated separately with the same exchange parameters from χ^2^ analysis.

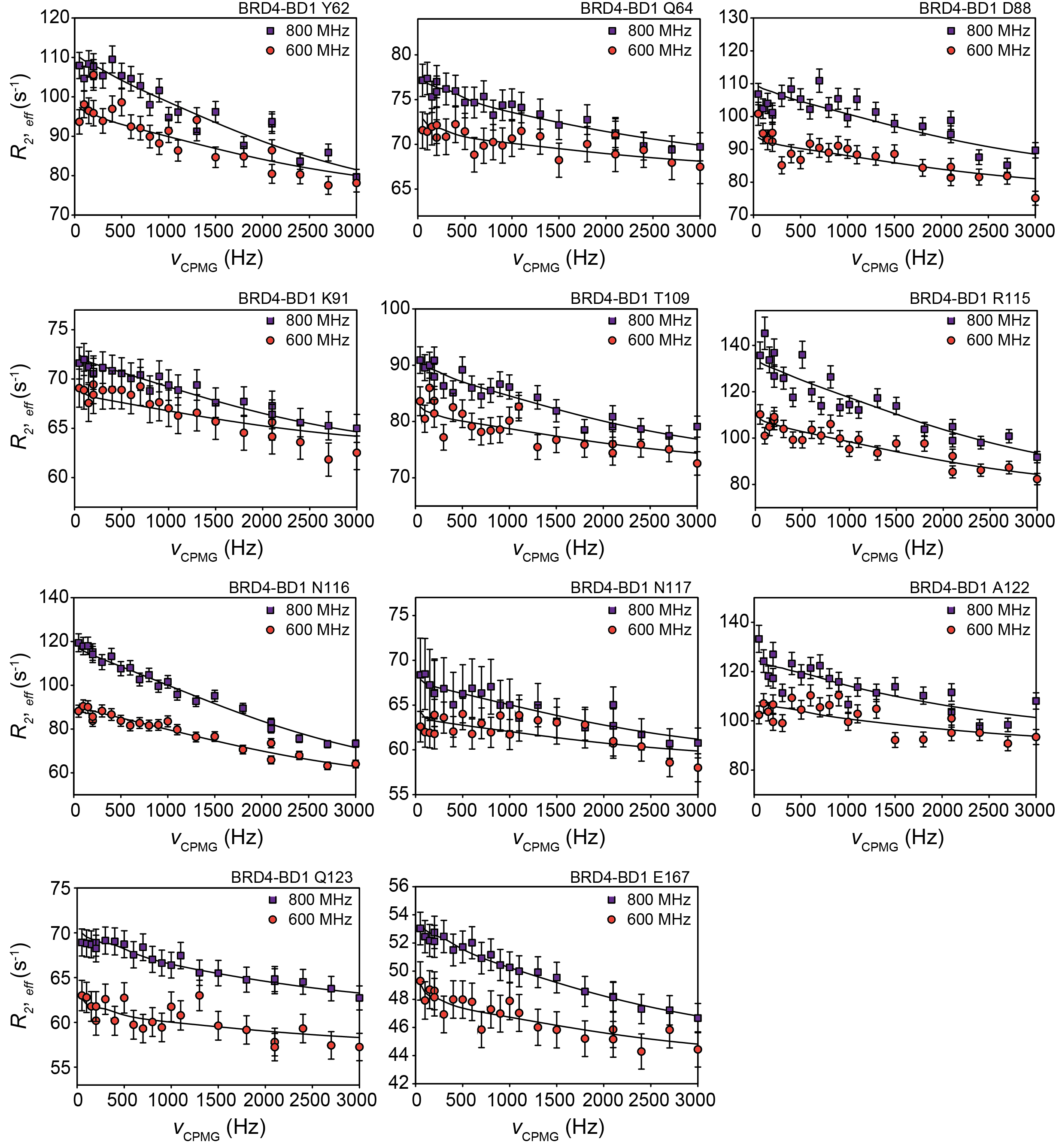

**Figure S15. Representative relaxation dispersion profiles of BRD4-BD1.** Effective relaxation rates (*R_2, eff_*) plotted as a function of CPMG frequency (ν_CPMG_). The ^1^H^N^ CPMG relaxation dispersion data were acquired at two magnet fields at 15 °C and globally fitted to a two-state exchange model using ChemEx. The exchange rate (*k_ex_*) at this site was estimated to be 14691 ± 1386 s^-1^.

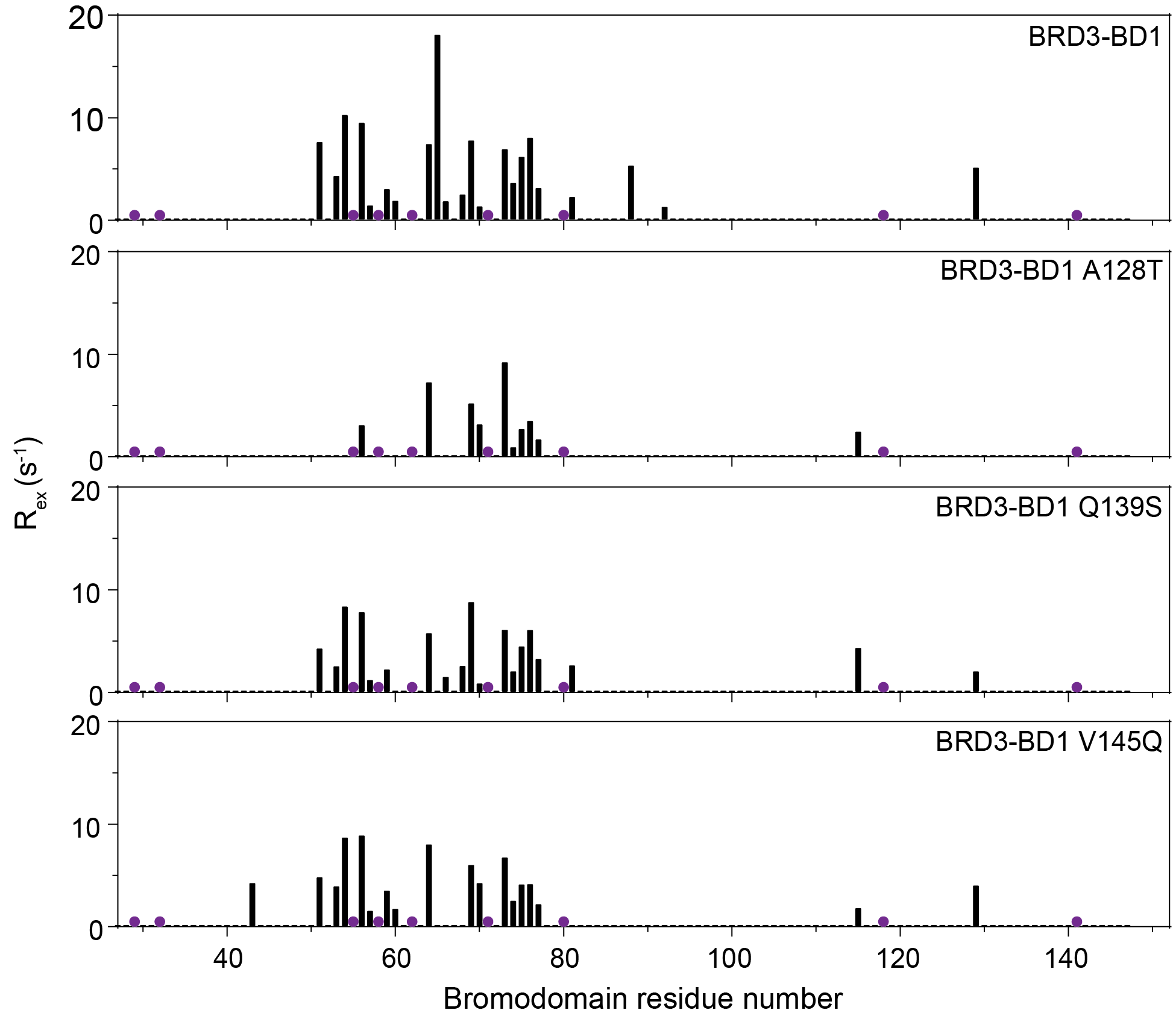

**Figure S16. Magnitude of dispersion (R_ex_) comparisons for BRD3-BD1, BRD3-BD1 A128T, BRD3-BD1 Q139S and BRD3-BD1 V145Q.** For each construct, R_ex_ (difference between the effective relaxation rates at low and high CPMG frequencies) plotted as a function of the residue number. The ^1^H^N^ CPMG relaxation dispersion data collected on the 800 MHz at 15 °C were used for this analysis. Missing assignments, overlapped/ambiguous peaks and prolines are indicated in purple spheres.

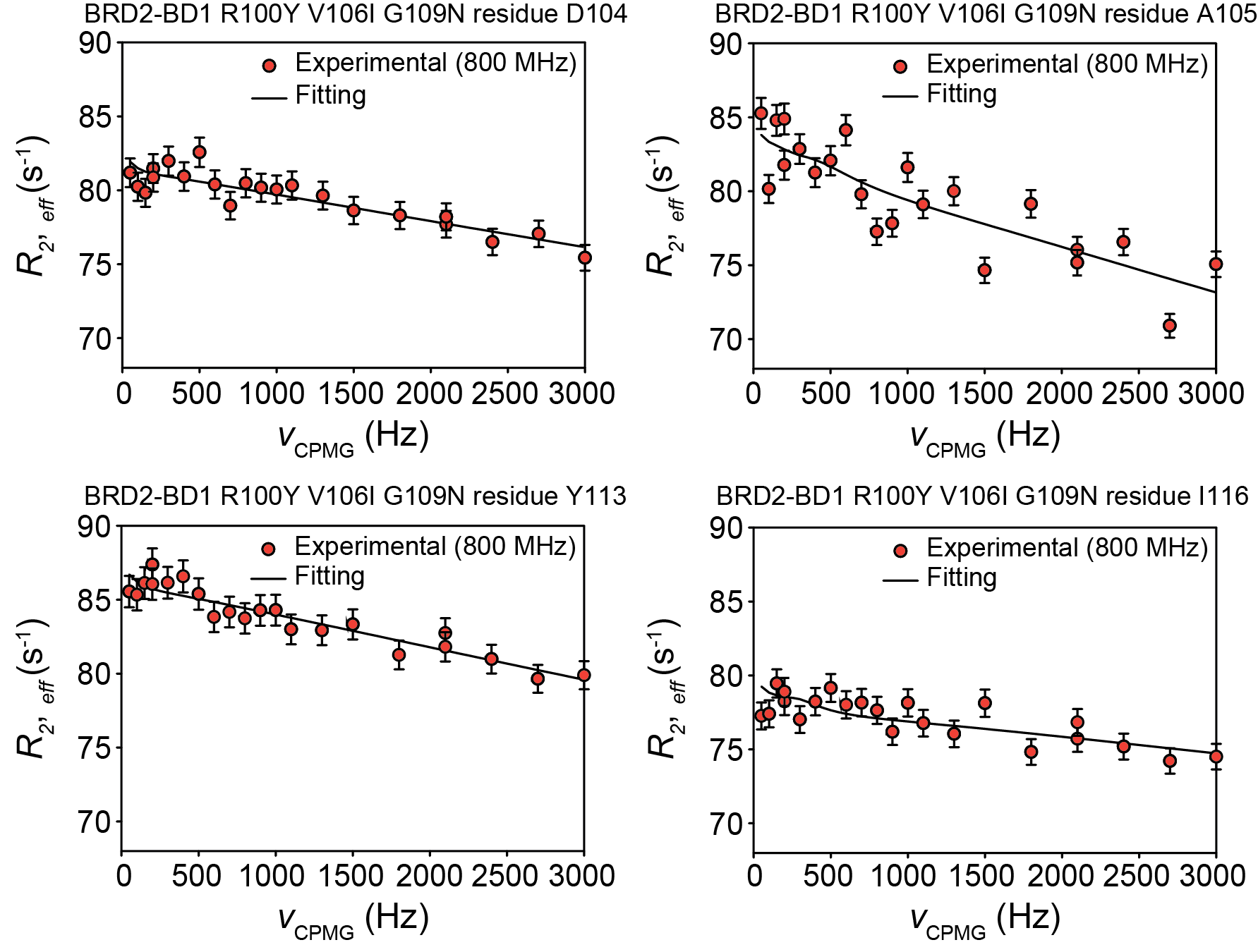

**Figure S17. Representative relaxation dispersion profiles of BRD2-BD1 R100Y V106I G109N.** Effective relaxation rates (*R_2, eff_*) plotted as a function of CPMG frequency (ν_CPMG_). The ^1^H^N^ CPMG relaxation dispersion data was acquired on the 800 MHz at 15 °C and globally fitted to a two-state exchange model using ChemEx. The mean exchange rate (*k_ex_*) was estimated to be 34900 s^-1^, calculated from 100 Monte Carlo simulations.

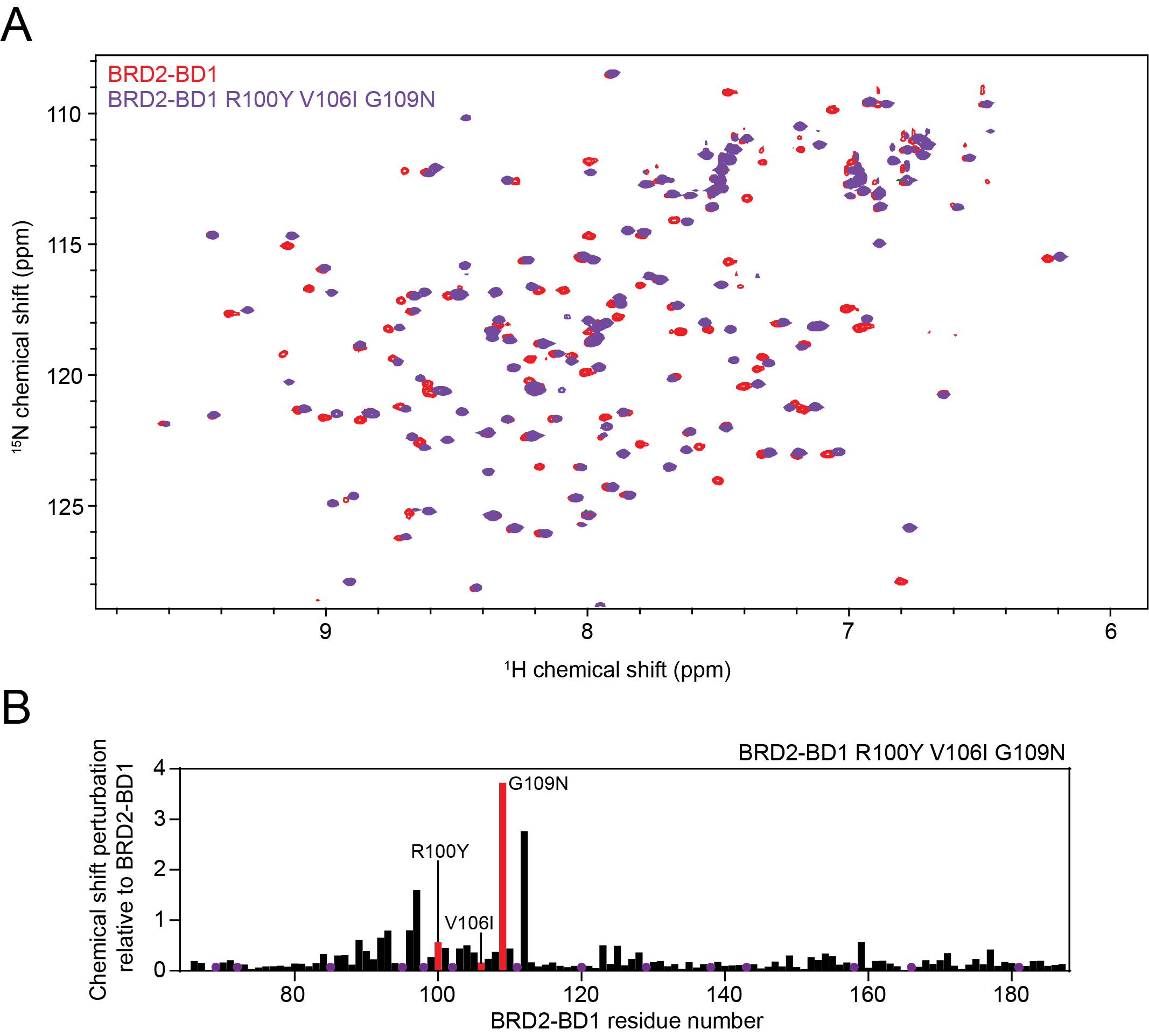

**Figure S18. Chemical shift perturbations analysis of BRD2-BD1 R100Y V106I G109N relative to BRD2-BD1. A.** 2D ^15^N-HSQC overlay of BRD2-BD1 (*red*) against BRD2-BD1 R100Y V106I G109N (*purple*). **B.** Quantitation of the chemical shift perturbations (CSPs) between BRD2-BD1 and BRD2-BD1 R100Y V106I G109N. Prolines and unassigned residues are indicated by purple circles.

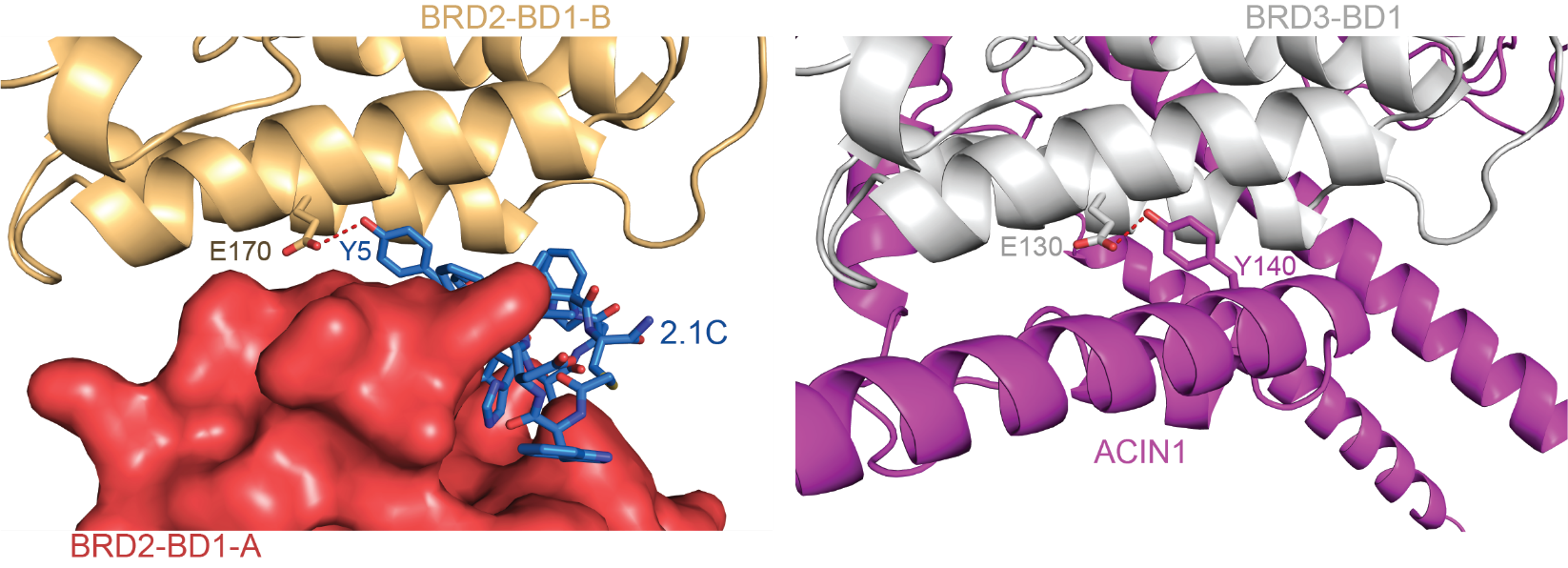

**Figure S19. Predicted structure of a BRD3-BD1:ACIN1 complex.** *Left.* Close-up of the X-ray crystal structure of BRD2-BD1 (*coral* and *light orange*) bound to 2.1C (*blue*). The hydrogen bond between BRD2-BD1 E170 and 2.1C Y5 is indicated by sticks and a dashed line. *Right*. Close-up of the Alphafold2-multimer prediction of the interaction made by BRD3-BD1 (*grey*) and a portion of ACIN1 (*magenta*) with the corresponding hydrogen bond (E130 of BRD3-BD1 and Y140 of ACIN1) indicated. The prediction was originally discovered in a search of the ProData database of human protein-protein interaction predictions (prodata.swmed.edu./humanPPI)^2^.

**REFERENCES**

1.	Divakaran, A. *et al.* Molecular Basis for the N-Terminal Bromodomain-and-Extra-Terminal-Family Selectivity of a Dual Kinase-Bromodomain Inhibitor. *J Med Chem* **61**, 9316–9334 (2018).

2.	Zhang, J. *et al.* Computing the Human Interactome. *bioRxiv* 2024.10.01.615885 (2024) doi:10.1101/2024.10.01.615885.
